# A molecular timescale for the origin of red algal-derived plastids

**DOI:** 10.1101/2020.08.20.259127

**Authors:** Jürgen F. H. Strassert, Iker Irisarri, Tom A. Williams, Fabien Burki

## Abstract

In modern oceans, eukaryotic phytoplankton is dominated by lineages with red algal-derived plastids such as diatoms, dinoflagellates, and coccolithophores. These lineages and countless others representing a huge diversity of forms and lifestyles all belong to four algal groups: cryptophytes, ochrophytes, haptophytes, and myzozoans. Despite the ecological importance of these groups, we still lack a comprehensive understanding of their evolution and how they obtained their plastids. Over the last years, new hypotheses have emerged to explain the acquisition of red algal-derived plastids by serial endosymbiosis, but the chronology of these putative independent plastid acquisitions remains untested. Here, we have established a timeframe for the origin of red algal-derived plastids under scenarios of serial endosymbiosis, using a taxon- and gene-rich phylogenomic dataset combined to Bayesian molecular clock analyses. We find that the hypotheses of serial endosymbiosis are chronologically possible, as the stem lineages of all red plastid-containing groups overlapped in time. This period in the Meso- and Neoproterozoic Eras set the stage for the later expansion to dominance of red algal-derived primary production in the contemporary oceans, which has profoundly altered the global geochemical and ecological conditions of the Earth.

## Introduction

Plastids (= chloroplasts) are organelles that allow eukaryotes to perform oxygenic photosynthesis. Oxygenic photosynthesis (hereafter simply photosynthesis) evolved in cyanobacteria around 2.4 billion years ago (bya), leading to the Great Oxidation Event—a rise of oxygen that profoundly transformed the Earth’s atmosphere and shallow ocean^1,2^. Eukaryotes later acquired the capacity to photosynthesise with the establishment of plastids by endosymbiosis. Plastids originated from primary endosymbiosis between a cyanobacterium and a heterotrophic eukaryotic host, leading to primary plastids in the first photosynthetic eukaryotes. There are three main lineages with primary plastids: red algae, green algae (including land plants), and glaucophytes—altogether forming a large group known as Archaeplastida^3,4^. Subsequently to the primary endosymbiosis, plastids spread to other eukaryote groups from green and red algae by eukaryote-to-eukaryote endosymbioses, i.e., the uptake of primary plastid-containing algae by eukaryotic hosts. These higher order endosymbioses resulted in complex plastids surrounded by additional membranes, some even retaining the endosymbiont nucleus (the nucleomorph), and led to the diversification of many photosynthetic lineages of global ecological importance, especially those with red algal-derived plastids (e.g., diatoms, dinoflagellates, and apicomplexan parasites)^5^.

The evolution of complex red algal-derived plastids has been difficult to decipher, mainly because the phylogeny of host lineages does not straightforwardly track the phylogeny of plastids. From the plastid perspective, phylogenetic and cell biological evidence strongly support a common origin of all complex red plastids^6–11^. This is at the centre of the chromalveolate hypothesis^12^, which proposed that the series of events needed to establish a plastid are better explained by a single secondary endosymbiosis in the common ancestor of alveolates, stramenopiles, cryptophytes, and haptophytes: the four major groups known to harbour complex red plastids. From the host side, however, the phylogenetic relationships of these four groups have become increasingly difficult to reconcile with a single origin of all complex red algal-derived plastids in a common ancestor. Indeed, over a decade of phylogenomic investigations have consistently shown that all red plastid-containing lineages are most closely related to a series of plastid-lacking lineages, often representing several paraphyletic taxa, which would require extensive plastid losses under the chromalveolate hypothesis (at least ten)^5^. This situation is further complicated by the fact that no cases of complete plastid loss have been demonstrated, except in a few parasitic taxa^13,14^.

The current phylogeny of eukaryotes has given rise to a new framework for explaining the distribution of complex red plastids. This framework, unified under the rhodoplex hypothesis, invokes the process of serial endosymbiosis, specifically a single secondary endosymbiosis between a red alga and a eukaryotic host, followed by successive higher-order—tertiary, quaternary—endosymbioses spreading plastids to unrelated groups^15^. Several models compatible with the rhodoplex hypothesis have been proposed, differing in the specifics of the plastid donor and recipient lineages^16–19^ (Fig. 1). However, these models of serial endosymbiosis remain highly speculative, in particular because we do not know if they are chronologically possible—did the plastid donor and recipient lineages co-exist? Addressing this important issue requires a reliable timeframe for eukaryote evolution, which has been challenging to obtain owing to a combination of complicating factors, notably: 1) uncertain phylogenetic relationships among the major eukaryote lineages; 2) the lack of genome-scale data for the few microbial groups with a robust fossil record; and 3) a generally poor understanding of methodological choices on the dates estimated for early eukaryote evolution.

**Fig. 1.**
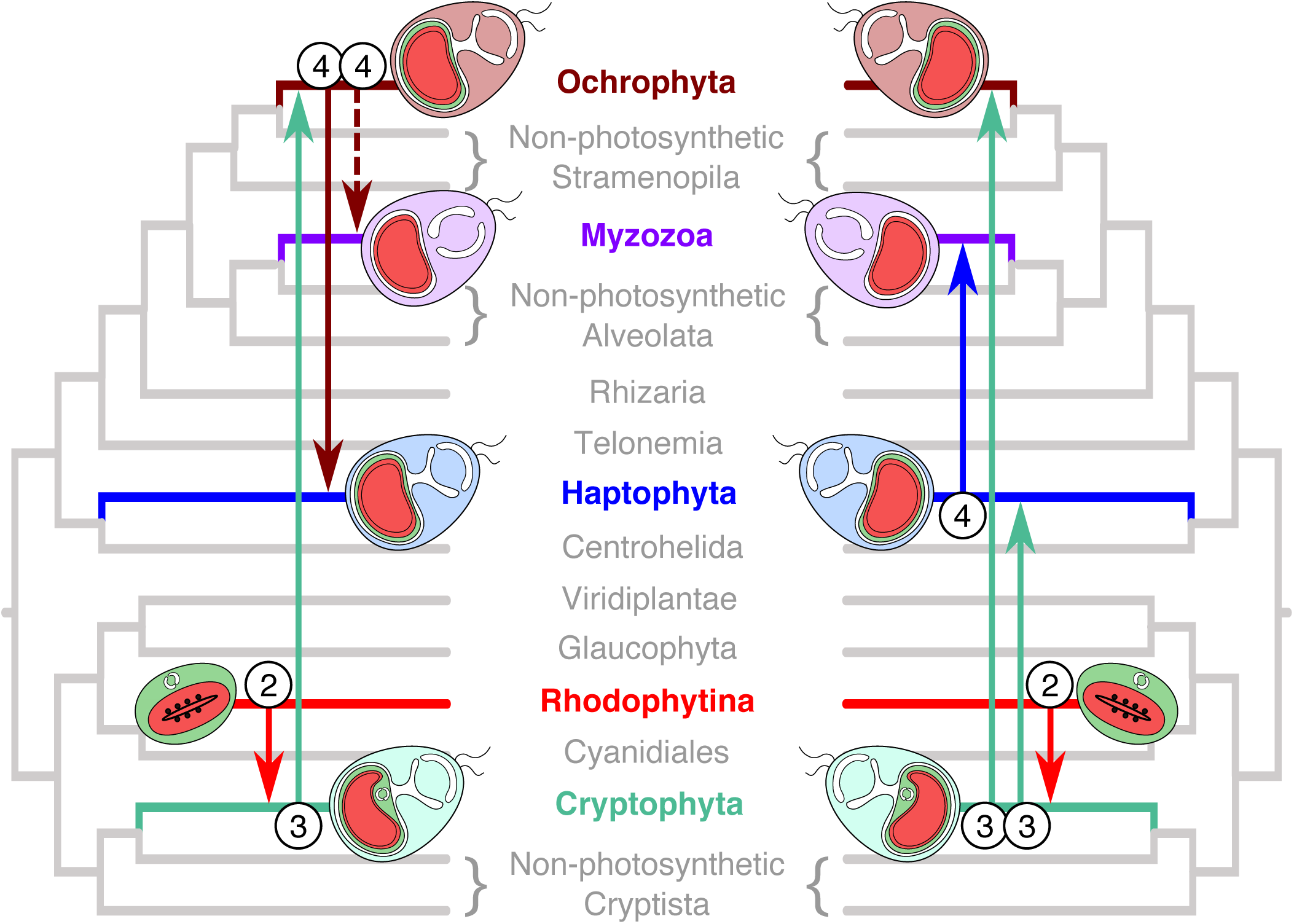
Serial plastid endosymbioses models as proposed by Stiller *et al*.^18^ (left) and Bodyl *et al*.^50^ (right). The tree topology shown here is based on the results obtained in our study. Further models have been suggested but are not compatible with this topology^15,83,84^. Numbers denote the level of endosymbiosis events. Note, Myzozoa were not included by Stiller et al.^18^ and the dashed line indicates engulfment of an ochrophyte by the common ancestor of Myzozoa as suggested by Ševčĺková *et al*.^19^.

Recent molecular clock analyses placed the origin of primary plastids in an ancestor of Archaeplastida in the Paleoproterozoic Era, between 2.1–1.6 bya^20^. The origin of red algae has been estimated to be in the late Mesoproterozoic to early Neoproterozoic (1.3–0.9 bya)^20^, after a relatively long lag following the emergence of Archaeplastida. However, an earlier appearance in the late Paleoproterozoic has also been proposed based on molecular analyses^21,22^. The earliest widely accepted fossil for the crown-group Archaeplastida is the multicellular filamentous red alga *Bangiomorpha* deposited ∼1.2 bya^23^. Recently, older filamentous fossils (*Rafatazmia* and *Ramathallus*) interpreted as crown-group red algae were recovered in the ∼1.6 Ga old Vindhyan formation in central India^24^. This taxonomic interpretation pushed back the oldest commonly accepted red algal fossil record by 400 million years, and consequently the origin of red algae and Archaeplastida well into the Paleoproterozoic Era. Evidence for the diversification of red algal plastid-containing lineages comes much later in the fossil record, which is apparent in the well-documented Phanerozoic continuous microfossil records starting from about 300 million years ago (mya). This period marks the diversification of some of the most ecologically important algae in modern oceans such as diatoms (ochrophytes), dinoflagellates (myzozoans), and coccolithophorids (haptophytes). If the fossil record is taken at face value, there is therefore a gap of over one billion years between the first appearance of crown-group eukaryotes interpreted as red algae in the early Mesoproterozoic and the later rise to ecological prominence of red algal-derived plastid-containing lineages^25^.

In this study, we combined phylogenomics and molecular clock analyses to investigate the chronology of the origin and spread of complex red plastids among distantly-related eukaryote lineages in order to test the general rhodoplex hypothesis^15^. We assembled a broad gene- and taxon-rich dataset (320 nuclear genes, 733 taxa), incorporating 33 well-established fossil calibrations, to estimate the timing of early eukaryote diversification. We explored the effect of a range of Bayesian molecular clock implementations, relaxed clock models and prior calibration densities, as well as two alternative roots for the eukaryote tree. Our analyses show that the hypotheses of serial endosymbiosis are chronologically possible, as most red algal plastid acquisitions likely occurred in an overlapping timeframe during the Mesoproterozoic and Neoproterozoic Eras, setting the stage for the subsequent evolution of the most successful algae on Earth.

## Results

### The phylogeny of eukaryotes

Molecular clock analyses rely on robust tree topologies. To obtain our reference topology, we derived two sub-datasets from the full dataset of 320 protein-coding genes and 733 taxa (Supplementary Fig. 1) to allow computationally intensive analyses: a 136-OTU dataset and a 63-OTU dataset (see Material and Methods). The 136-OTU dataset was used in Maximum Likelihood (ML) inference using the best fitting site-heterogeneous LG+C60+G+F-PMSF model (hereafter simply *ML-c60*) based on a concatenated alignment, as well as with a supertree method consistent with the Multi Species Coalescent model (MSC) in a version of the alignment where the taxa within OTUs were retained individually. The concatenated 63-OTU dataset was analysed by Bayesian inference using the site heterogeneous CAT+GTR+G (hereafter simply *catgtr*) and LG+C60+G (*BI-c60*) models, as well as in posterior predictive analyses (PPA) to compare the fit of both models.

The *ML-c60* and MSC-derived trees are in good overall agreement with current knowledge of the broad eukaryote evolution. The *ML-c60* tree recovered many existing supergroups with maximal bootstrap support (Supplementary Fig. 2), namely the TSAR assemblage^26^, Haptista, Cryptista, Discoba, Amoebozoa and Obazoa^27,28^. Archaeplastida was also recovered monophyletic, albeit with lower bootstrap support (86%), but this supergroup has previously lacked support in phylogenomic analyses (see Discussion). The relationships among these supergroups are also consistent with published work, most notably the recurrent affinity between Cryptista and Archaeplastida (CA clade), the branching of Haptista with *Ancoracysta twista* in ML analyses, and the placement of this group deep in the tree (here sister to the CA clade with 95% bootstrap support). The MSC analyses mostly recapitulate the same observations (Supplementary Fig. 3), although with the exceptions of TSAR and Archaeplastida due to the unresolved positions of telonemids as well as red algae and Cryptista, respectively. Taken together, the MSC analyses either supported the results of the concatenated ML analysis, or were inconclusive rather than conflicting.

**Fig. 2.**
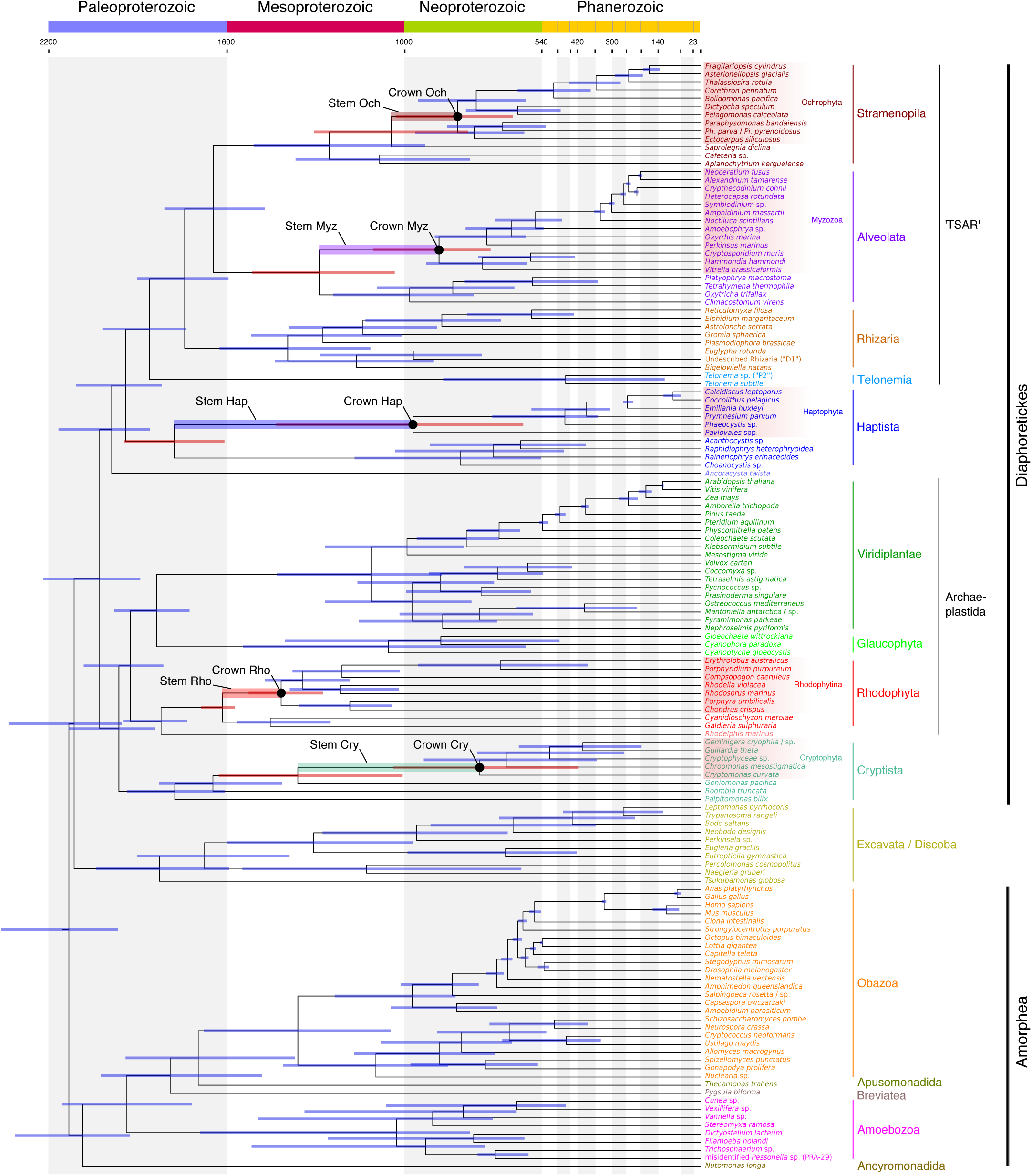
Time-calibrated phylogeny of extant eukaryotes. Divergence times were inferred with MCMCTree under an autocorrelated relaxed clock model and 33 fossil calibration points as soft-bound uniform priors (Table 1). The tree topology was reconstructed using IQ-TREE under the LG+C60+G+F model and a constrained tree search following the OTU-reduced CAT+GTR+G topology (Supplementary Fig. 4). Approximate likelihood calculations on the 320 gene concatenation under LG+G and a birth-death tree prior were used. Bars at nodes are 95% HPD. Bars corresponding to the first and last common ancestors of extant red plastid-donating and -containing lineages are highlighted in red and their stems are shaded as indicated. Crowns denote the common ancestors of the extant members of these groups. An absolute time scale in Ma and a geological time scale are shown. The tree depicted here was rooted on Amorphea. An equivalent time-calibrated tree rooted on Excavata is shown in the Supplementary Fig. 6. Cry – Cryptophyta, Hap – Haptophyta, Myz – Myzozoa, Och – Ochrophyta, Rho – Rhodophytina.

**Fig. 3.**
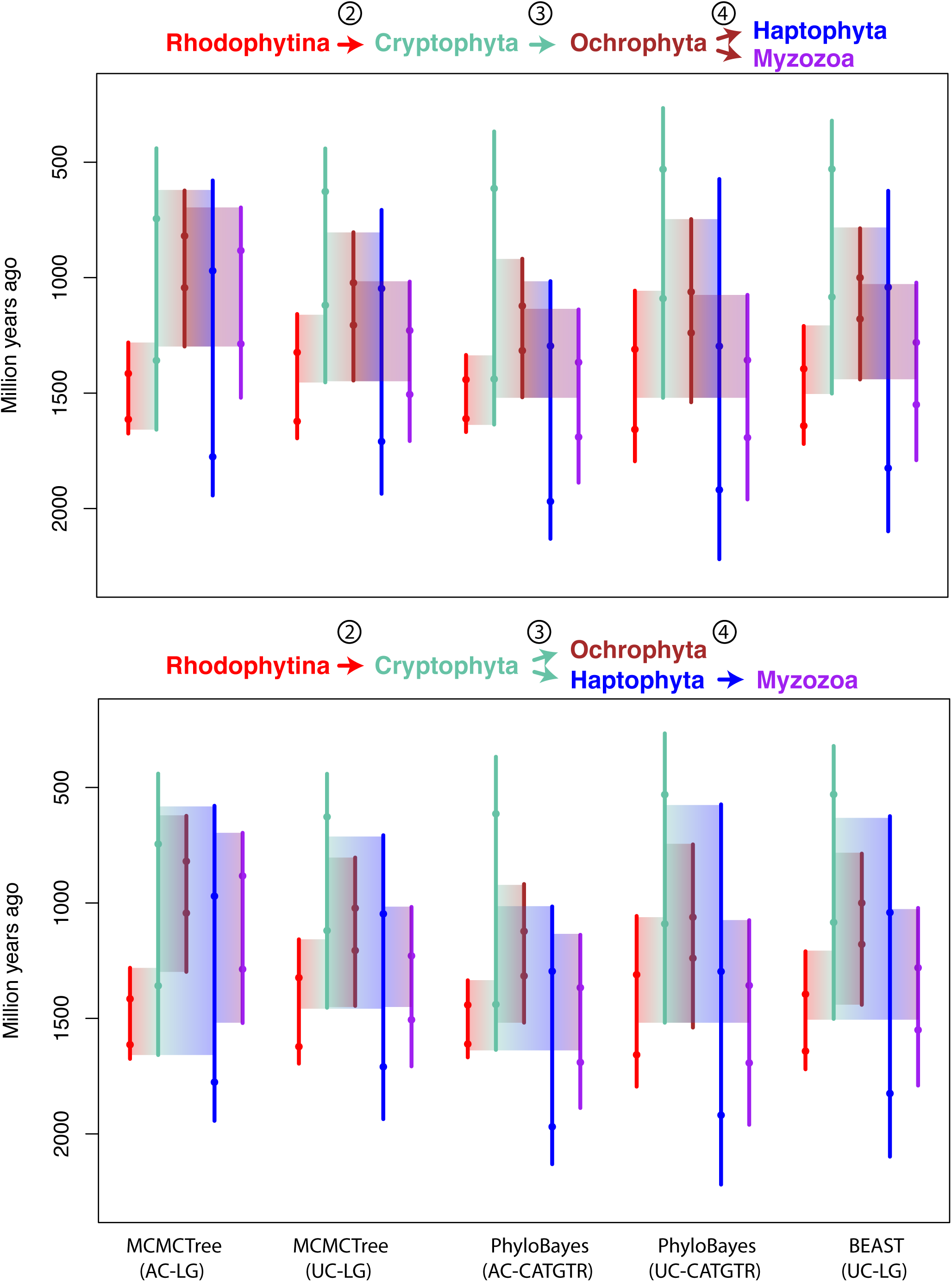
Summary of inferred timeframes for the spread of complex red plastids using different software and models. Vertical lines correspond to the lower and upper 95% HPD intervals from the nodes defining the branch of interest and dots indicate their posterior mean divergences. Faded boxes represent the temporal windows for the secondary endosymbioses, constrained by the HPDs and tree topology under the two proposed symbiotic scenarios. Numbers at arrows denote the level of endosymbiosis events (compare with Fig. 1). AC – autocorrelated clock model, UC – uncorrelated clock model.

The *catgtr* tree (Supplementary Fig. 4) received full posterior probabilities (PP) for all bifurcations and is globally consistent with the *ML-c60* analyses (Supplementary Fig. 2), albeit with one important difference for understanding the evolution of plastids. In this tree, the positions of Haptista and *A. twista* were paraphyletic with respect to TSAR. A Bayesian reanalysis of the 63-OTU dataset under the *BI-c60* model recapitulated the *ML-c60* topology, suggesting that these topological differences derived from the chosen evolutionary model and not from the use of Bayesian or ML inference (Supplementary Fig. 5). Posterior predictive tests showed that the *catgtr* model better accounts for compositional heterogeneity than *BI-c60* (although neither models did it fully; Supplementary Table 1) and the *catgtr* topology could not be rejected by ML (approximately unbiased test; p-AU = 0.182). Taken together, these results favoured the relationships described under the *catgtr* model. Therefore, the backbone topology for the divergence time estimation was inferred under a set of constraints defined by the best *catgtr* topology but using the 136-OTU dataset, which contains a broader taxon sampling allowing to more precisely place fossil calibrations.

**Table 1.** Calibrations used for dating the eukaryote tree of life.

| Clade | Age constraint (Ma) | Type | Eon | Refs. |
| --- | --- | --- | --- | --- |
| Rhodophyta<sup>a</sup> | 1,600 | Min | Proterozoic | 24 |
| Bangiophyceae/Florideophyceae | 1,600 | Max | Proterozoic | --b |
| Bangiophyceae/Florideophyceae | 1,047 | Min | Proterozoic | 23,85 |
| Metazoa | 833 | Max | Proterozoic | 86 |
| Ciliophora<sup>a</sup> | 740 | Min | Proterozoic | 87 |
| Euglyphidae<sup>a</sup> | 736 | Min | Proterozoic | 88,89 |
| Evosea/Tubulinea<sup>a</sup> | 736 | Min | Proterozoic | 89,90 |
| Tubulinea | 736 | Max | Proterozoic | --b |
| Chlorophyta<sup>a</sup> | 700 | Min | Proterozoic | 91 |
| Eumetazoa | 636 | Max | Proterozoic | 86 |
| Deuterostomia | 636 | Max | Proterozoic | 86 |
| Chordata | 636 | Max | Proterozoic | 86 |
| Bilateria | 636 | Max | Proterozoic | 86 |
| Arthropoda | 636 | Max | Proterozoic | 86 |
| Metazoa | 635 | Min | Proterozoic | 29 |
| Eumetazoa | 550 | Min | Proterozoic | 86 |
| Bilateria | 550 | Min | Proterozoic | 86 |
| Mollusca | 549 | Max | Proterozoic | 86 |
| Foraminifera<sup>a</sup> | 542 | Min | Proterozoic | 92,93 |
| Bacillariophyta | 541 | Max | Proterozoic | 94 |
| Embryophyta | 540 | Max | Phanerozoic | 20,95 |
| Mollusca | 532 | Min | Phanerozoic | 86 |
| Deuterostomia | 515 | Min | Phanerozoic | 86 |
| Chordata | 514 | Min | Phanerozoic | 86 |
| Arthropoda | 514 | Min | Phanerozoic | 86 |
| Embryophyta | 470 | Min | Phanerozoic | 96 |
| Angiosperms/Gymnosperms | 470 | Max | Phanerozoic | --b |
| Euglenales/Eutreptiales<sup>a</sup> | 450 | Min | Phanerozoic | 97 |
| Chytridiomycotaa | 410 | Min | Phanerozoic | 98,99 |
| Tubulinea | 405 | Min | Phanerozoic | 100 |
| Ascomycotaa | 400 | Min | Phanerozoic | 101 |
| Angiosperms/Gymnosperms | 385 | Min | Phanerozoic | 102 |
| Angiosperms | 385 | Max | Phanerozoic | --b |
| Basidiomycotaa | 360 | Min | Phanerozoic | 103 |
| Amniota | 333 | Max | Phanerozoic | 86 |
| Amniota | 318 | Min | Phanerozoic | 86 |
| Core dinoflagellates (excl. Noctilucales) | 300 | Max | Phanerozoic | 42 |
| Coccolithales/Isochrysidales | 260 | Max | Phanerozoic | 42 |
| Core dinoflagellates (excl. Noctilucales) | 235 | Min | Phanerozoic | 104 |
| Peridinales | 235 | Max | Phanerozoic | --b |
| Gonyaulacales | 235 | Max | Phanerozoic | --b |
| Coccolithales/Isochrysidales | 225 | Min | Phanerozoic | 105 |
| Calcidiscaceae/Coccolithaceae | 225 | Max | Phanerozoic | --b |
| Peridinales | 210 | Min | Phanerozoic | 104 |
| Gonyaulacales | 200 | Min | Phanerozoic | 104 |
| Bacillariophyta | 190 | Min | Phanerozoic | 94 |
| Pennales | 190 | Max | Phanerozoic | --b |
| Euarchontoglires | 165 | Max | Phanerozoic | 86 |
| Angiosperms | 130 | Min | Phanerozoic | 106 |
| Eudicotyledons (Tricoplastes) | 130 | Max | Phanerozoic | --b |
| Eudicotyledons (Tricoplastes) | 124 | Min | Phanerozoic | 107,108 |
| Aves sensu stricto | 87 | Max | Phanerozoic | 86 |
| Pennales | 75 | Min | Phanerozoic | 94 |
| Aves sensu stricto | 66 | Min | Phanerozoic | 86 |
| Calcidiscaceae/Coccolithaceae | 65 | Min | Phanerozoic | 109,110 |
| Euarchontoglires | 61 | Min | Phanerozoic | 86 |

<sup>a</sup>Max = 1,900 Ma was used; see for example Knoll<sup>111</sup> and Eme *et al.*<sup>41</sup>.
<sup>b</sup>Max based on Min age of ancestor.

### Timescale for eukaryote evolution

Our molecular clock analyses revealed congruent dates inferred under comparable analytical conditions (i.e., clock models, prior calibration densities, and root positions; see below and Material and Methods) and by all tested implementations (MCMCTree, PhyloBayes, BEAST). The fossils used as calibrations were chosen to span a wide diversity of lineages (Table 1), and we followed a conservative approach when interpreting the fossil record by choosing only fossils widely accepted by the paleontological community. We also included one generally uncontested biomarker to constrain the emergence of extant Metazoa in order to expand the otherwise sparse Proterozoic fossil record (24-isopropyl cholestane)^29^. Our analyses placed the root of the eukaryote tree well into the Paleoproterozoic Era (Fig. 2). This Era also saw the origin of primary plastids in the common ancestor of Archaeplastida, which likely took place between 2,137 and 1,807 mya. Crown group red algae were inferred in the Paleoproterozoic between 1,984 and 1,732 mya. These age ranges are provided as a conservative approach encompassing all performed analyses (with the exception of those using t-cauchy distributions with long tails, see below).

From red algae, plastids then spread to distantly-related groups of eukaryotes by at least one secondary endosymbiosis. Plastid phylogenies have consistently shown that this red algal donor lineage belonged to a stem lineage of Rhodophytina^30^, i.e., it lived after the split of Cyanidiophyceae and the rest of red algae (Fig. 2). Based on Fig. 2, we inferred a time range for this donor lineage to be between 1,675 and 1,281 mya. The origination period for the lineages currently harbouring red algal-derived plastids were inferred as follows: cryptophytes between 1,658 and 440 mya; ochrophytes between 1,298 and 622 mya; haptophytes between 1,943 and 579 mya; myzozoans between 1,520 and 696 mya (dates refer to the 95% HPD intervals in Fig. 2). Thus, the stems of all extant lineages containing red algal plastids—along which these plastids were acquired—overlap chronologically. This overlap was consistent across all performed analyses and defines time windows during which endosymbiotic transfers, as proposed by the serial endosymbiosis hypotheses, could reconcile plastid and nuclear phylogenies (Figs. 2 and 3; Supplementary Figs. 6, 7, and Supplementary Data 1). Note that PhyloBayes analyses with the autocorrelated clock model displayed the shortest confidence intervals, while the MCMCTree analyses with the same clock model recovered generally younger ages for the origin of ochrophytes and myzozoans (Fig. 3), but these differences did not alter the observation of time overlap for the putative plastid acquisitions.

To better understand the effect of methodological choices on divergence times, we performed a battery of sensitivity analyses with MCMCTree that tested different combinations of clock models, prior calibration densities, and the position of the eukaryote root (32 sensitivity analyses in total; Supplementary Data 1 and Material and Methods). These analyses indicated that the prior calibration distributions had the strongest effect on the inferred divergence times, followed by the clock model, and lastly the root position. Prior calibrations model the uncertainty of fossil ages and their proximity to the cladogenetic events being calibrated, and thus different distributions can be understood as more literal (skew-normal), loose (t-cauchy), or conservative (uniform) interpretations of the fossil record^31^. As expected from the prior distributions, we observed younger overall ages with skew-normal calibrations (mean of 905 Ma) compared to uniform (mean of 926 Ma) or t-cauchy distribution with short or long tails (means of 1,105 and 1,591 Ma, respectively; Supplementary Data 1). The excessively old ages and wide 95% HPD intervals (mean of 517 *vs*. 306 to 372 Ma) inferred with long-tailed t-cauchy distribution were considered biologically implausible, and thus disregarded in the following. The clock model had a modest impact on the posterior dates, with the autocorrelated clock model generally producing slightly younger ages and narrower intervals (mean ages 1,138 Ma; 95% HPD width 356 Ma) than the uncorrelated clock (mean age 1,125 Ma and 95% HPD width 416 Ma). The younger ages inferred under the autocorrelated clock model were more apparent when uniform calibrations were applied (mean ages of 904 *vs*. 947 Ma) and this trend was inverted in the case of skew-normal distributions (913 *vs*. 896 Ma). CorrTest indicated that branch lengths are most likely correlated (CorrScore = 0.99808, p < 0.001), suggesting that autocorrelated models might better model our dataset. Finally, the use of the two alternative roots, either on Amorphea or on Discoba, had the smallest effect on the posterior ages. Only marginal differences were observed on the overall node mean times of (1,134 *vs*. 1,132 Ma for the Amorphea and Discoba roots, respectively) and interval widths (381 *vs*. 388 Ma). The only exceptions were basal relationships within Discoba, which were noticeably older rooting the tree on this group (Fig. 2; Supplementary Fig. 6).

Further sensitivity analyses were performed with PhyloBayes to confirm the effects of the clock model choice (Supplementary Data 1). Additionally, we tested the effect of the substitution model by comparing LG+G with the *catgtr* mixture model. We observed a higher impact of the clock model than the evolutionary model. In this case, the uncorrelated clock model led to younger ages and wider HPD intervals than the autocorrelated clock (mean age of 942 *vs*. 1024 Ma, and 95% HPD width of 420 *vs*. 321 Ma, respectively). The *catgtr* model tended to infer wider intervals than the LG+G model (e.g., 95% HPD width of 356 *vs*. 321 Ma under the autocorrelated clock model). Finally, we tested a third commonly used Bayesian implementation (BEAST) using the LG+G substitution model, the Amorphea root, and uniform prior calibrations. The obtained mean posterior ages and 95% HPD intervals were in overall agreement with those obtained by MCMCTree and PhyloBayes (Supplementary Data 1) corroborating our inferred dates in these analyses.

## Discussion

### The eukaryote tree of life and the rhodoplex hypothesis

In the last 15 years, the tree of eukaryotes has been extensively rearranged based on phylogenomics. We assembled a dataset containing a dense eukaryotic-wide taxon sampling with 733 taxa and analysed subsamples of it with a variety of mixture models to resolve several important uncertainties that are key for understanding plastid evolution. Notably, we recovered the monophyly of Archaeplastida, which has previously been strongly supported by plastid evidencee.g.,^11,32^ but not by host (nuclear) phylogenetic markerse.g., ^33–36^. This topology is consistent with a recent exhaustive phylogenomic study of nuclear markers^4^ as well as with the long-held view of a single point of entry of photosynthesis in eukaryotes from cyanobacteria through the establishment of primary plastids^37^ (with the exception of the chromatophores in *Paulinella*^38^). The supergroup Cryptista, which includes the red algal plastid-containing cryptophytes, was placed as sister to Archaeplastida. The association of Cryptista and Archaeplastida has been suggested before, often even disrupting the monophyly of Archaeplastida^35,36^, but our analyses robustly recovered the sister relationship of these two major eukaryote clades. Another contentious placement has concerned the supergroup Haptista^26,35,36,39^, which includes the red algal plastid-containing haptophytes, as well as its possible relationship to the orphan lineage *Ancoracysta twista*^40^. The best-fitting *catgtr* model favoured the position of Haptista as sister to TSAR and placed *A. twista* deeper in the tree, unrelated to Haptista. We observed that the model (*catgtr vs. lgc60*) rather than the method or program is determinant in placing *A. twista* relative to Haptista—*lgc60* always placed these two lineages together, both in ML and Bayesian, but *catgtr* never did—and that variation in the taxon-sampling did not drastically modify this association.

Given this well-resolved tree of eukaryotes, we then mapped the position of the groups with red algal-derived plastids (Fig. 2). The rhodoplex hypothesis explains the distribution of these plastids by a series of endosymbioses, positing that plastids were separately acquired in the stem lineages of cryptophytes, ochrophytes, haptophytes, and myzozoans^15^. While the rhodoplex hypothesis remains speculative, it provides the benefit of reconciling the accumulating discrepancies between plastid and host phylogenies that existed under the previously prevalent chromalveolate hypothesis^12^. In the chromalveolate hypothesis, a red plastid was acquired by secondary endosymbiosis in the common ancestor of all red plastid-bearing lineages, and multiple subsequent losses were invoked to explain the patchy distribution of red plastids across the eukaryote tree. Our phylogeny is consistent with other recent analyses in rendering this scenario impossible^11,19,26,35,36^: since Cryptista branch as sister to Archaeplastida, but the other red plastid lineages branch elsewhere in the tree, the common ancestor of red plastid-bearing lineages corresponds to one of the earliest nodes in the tree (Fig. 2), which was ancestral to the cell that acquired primary plastids at the origin of Archaeplastida. As has been pointed out before^35^, this scenario would require a red alga to travel backwards in time to be engulfed by one of its distant ancestors. In contrast, our analyses indicate that the inferred red algal donor lineage (stem Rhodophytina) was contemporaneous with stem cryptophytes, haptophytes, and myzozoans, but red plastids were unlikely to have been acquired directly by ochrophytes, which in all likelihood had not yet diverged from other stramenopiles at the time (although their 95% HPDs marginally overlap). Moreover, all red algal plastid-containing lineages are most closely related to lineages without plastids (Fig. 2), and for which conclusive evidence for a past photosynthetic history does not exist. This situation would require many plastid losses if red plastids had been established early and vertically transmitted, indicating that the chromalveolate hypothesis no longer provides a compelling explanation for the distribution of red plastids in extant eukaryotes^5^.

### Timing the early evolution of eukaryotes

We present a detailed molecular clock analysis providing a timeframe for eukaryote evolution. Our estimates inferred an age for the Last Eukaryote Common Ancestor (LECA) between 2,386 and 1,958 mya, which is generally older than in other molecular clock analyses based on phylogenomic data^41–43^. However, an early Paleoproterozoic origin of extant eukaryotes fits with the oldest definitive crown-eukaryote fossils, the putative red algae *Rafatazmia chitrakootensis* and *Ramathallus lobatus* from 1,600 mya^24^, which imply that eukaryotes must have originated before this time. It also fits with the recent discoveries of multicellular eukaryotes in different groups of Proterozoic fossils indicating that eukaryotes were already complex in deep times. For example, the chlorophyte fossil *Proterocladus antiquus* in ∼1,000 Ma old rocks, taken as evidence for a much earlier appearance of multicellularity in this group of green algae^44^, is in line with our results (Fig. 2). Similarly, multicellular organic-walled microfossils with affinity to fungi were recently reported in the 1– 0.9 Ga old Grassy Bay Formation^45^, which pushes back the emergence of fungi by 500 Ma compared to previous studies^46^. In our analyses, fungi were estimated to originate even before (1,759 to 1,078 Ma), consistently with the presence of multicellular organisms around 1 bya. More generally, an early Paleoproterozoic origin of eukaryotes would also be in line with records of aggregative multicellularity appearing more than 2 bya, although the eukaryote affiliation of these fossils is debated^47,48^.

Our estimated dates placed the common ancestor of Archaeplastida also in the Paleoproterozoic Era, suggesting that eukaryotes acquired primary plastids, and thus photosynthesis, early on. This early origin of plastids is consistent with the putative photosynthetic crown-Archaeplastida acritarch *Tappania*, which offers circumstantial evidence that eukaryotes exploited photosynthesis from the onset of eukaryote evolution^49^. It is also in agreement with a recent molecular clock analysis of early photosynthetic eukaryotes that placed the origin of Archaeplastida at ∼1,900 mya^20^. Furthermore, total group red algae are generally thought to contain some of the best minimum constraints for the existence of fully photosynthetic eukaryotes, most convincingly the late Mesoproterozoic *Bangiomorpha* resembling extant large bangiophytes^23^, but also the earlier *R. chitrakootensis* and *R. lobatus*, probably also multicellular from 1,600 Ma^24^. Thus, both the fossil record and molecular clock inferences support a Paleoproterozoic origin of primary plastids, and an early Mesoproterozoic origin of red algae.

For any endosymbiotic relationship to be established, the endosymbiont and the host must live at the same time and in the same place in order to interact. While the time intervals for plastid acquisition are sometimes relatively wide in our analyses (Fig. 2), they can be further constrained by overlaying the direction of plastid transfers proposed in the different models congruent with the rhodoplex hypothesis (Fig. 1). Of these models, only those by Stiller *et al*.^18^ and Bodyl *et al*.^50^ are compatible with our tree (Fig. 2), because the other models assumed the monophyly of haptophytes with cryptophytes and the acquisition of plastids in an ancestor of that group. In both compatible models, the secondary engulfment of a red alga by a stem cryptophyte can be constrained by the minimum age of the plastid donor lineage, i.e., the age of the last Rhodophytina common ancestor (Fig. 2). This would indicate a rather early plastid acquisition by cryptophytes and a relatively long time span before the diversification of extant cryptophytes. In fact, if a red plastid-bearing cryptophyte was subsequently acquired by tertiary endosymbiosis by a stem ochrophyte, as both models proposed, this acquisition likely took place at latest 819 mya (95% HPD: 633–1,017), thus providing a maximum constraint for the age of crown cryptophytes. The model of Bodyl *et al*.^50^ also proposed that cryptophytes were acquired by an ancestor of haptophytes in a parallel tertiary endosymbiosis, thus resulting in further constraints on the cryptophyte stem lineage. However, phylogenetic analyses of plastid-targeted proteins have instead supported an ochrophyte origin for the haptophyte plastid^18^. Following the same logic, the most likely time ranges of plastid acquisitions in all four red algal plastid-containing lineages and for the models and data compatible with our tree are presented in Table 2. Strikingly, this approach to chronologically constraining red plastid endosymbioses revealed that the serial transfers all took place in a relatively short time window between 650 and 1,079 million years from the initial secondary endosymbiosis in a stem cryptophyte to the establishment of plastids in the ancestors of all modern-day red algal plastid-containing lineages (narrowest and widest 95% HPD widths for the totality of the four endosymbioses; Table 2).

**Table 2.** Inferred time intervals for the acquisition of red algal-derived plastids by serial endosymbioses. Time intervals have been calculated from overlapping 95% HPD intervals, constrained by the order of symbioses in the models of Stiller *et al*.^18^ or Bodyl *et al*.^50^. Numbers are in million years. Cry – Cryptophyta, Hap – Haptophyta, Myz – Myzozoa, Och – Ochrophyta, Rho – Rhodophytina.

|  | MCMCTREE |  |  |  |  |  | PHYLOBAYES |  |  |  |  |  | BEAST |  |  |
| --- | --- | --- | --- | --- | --- | --- | --- | --- | --- | --- | --- | --- | --- | --- | --- |
|  | AUTOCORRELATED |  |  | UNCORRELATED |  |  | AUTOCORRELATED |  |  | UNCORRELATED |  |  | UNCORRELATED |  |  |
| Stiller *et al.* |  |  |  |  |  |  |  |  |  |  |  |  |  |  |  |
|  | max | min | width | max | min | width | max | min | width | max | min | width | max | min | width |
| Rho => Cry | 1,658 | 1,281 | 377 | 1,453 | 1,158 | 296 | 1,636 | 1,335 | 301 | 1,517 | 1,057 | 461 | 1,502 | 1,210 | 292 |
| Cry => Och | 1,298 | 623 | 675 | 1,446 | 804 | 642 | 1,518 | 918 | 600 | 1,517 | 747 | 771 | 1,441 | 786 | 654 |
| Och => Hap | 1,298 | 623 | 675 | 1,446 | 804 | 642 | 1,518 | 1051 | 467 | 1,517 | 747 | 771 | 1,441 | 786 | 654 |
| Och => Myz | 1,298 | 696 | 602 | 1,446 | 1,017 | 428 | 1,518 | 1,138 | 380 | 1,517 | 1,075 | 443 | 1,441 | 1,022 | 419 |
| Bodył *et al.* |  |  |  |  |  |  |  |  |  |  |  |  |  |  |  |
|  | max | min | width | max | min | width | max | min | width | max | min | width | max | min | width |
| Rho => Cry | 1,658 | 1,281 | 377 | 1,453 | 1,158 | 296 | 1,636 | 1,335 | 301 | 1,517 | 1,057 | 461 | 1,502 | 1,210 | 292 |
| Cry => Och | 1,298 | 623 | 675 | 1,446 | 804 | 642 | 1,518 | 918 | 600 | 1,517 | 747 | 771 | 1,441 | 786 | 654 |
| Cry => Hap | 1,658 | 579 | 1,079 | 1,453 | 707 | 747 | 1,636 | 1051 | 585 | 1,517 | 573 | 944 | 1,502 | 624 | 878 |
| Hap => Myz | 1,520 | 696 | 824 | 1,453 | 1,017 | 436 | 1,636 | 1,138 | 498 | 1,517 | 1,075 | 443 | 1,502 | 1,022 | 480 |

### The origin and rise of algae

The fossil and biomarker records document the increasing abundance and rise to ecological prominence of primary endosymbiotic algae in the oceans, including red algae, ∼659–645 mya^51^. Red plastid-bearing algae, notably diatoms, coccolithophores and dinoflagellates, started to expand later after the Permo-Triassic transition ∼250 mya and have since remained major primary producers in the oceans^52^. By contrast, our molecular clock analyses indicate that the events underlying the evolutionary origin of these algae, including the primary plastid endosymbiosis, the origin of red algae, and the subsequent spread of red plastids across the eukaryote tree, all pre-dated the ecological expansion of these groups by at least ∼0.5–1 Ga. This long time lag suggests that the selective pressures underpinning the establishment of primary and complex plastids in the Palaeo- and Mesoproterozoic Eras are distinct from those that drove their expansion ∼1 Ga later.

The Proterozoic ocean was poor in essential inorganic nutrients, such as phosphate and nitrogen, which may have limited the expansion of larger eukaryotic algae into the open marine realm^52^. Yet, these oligotrophic environments may have been favourable to the origin and early evolution of plastid-containing lineages. Predatory behaviours have been demonstrated in green algae and non-photosynthetic direct relatives of red algae, suggesting that mixotrophy was a key intermediate stage in early evolution of Archaeplastida^53,54^. More generally, mixotrophy is increasingly recognised as the default lifestyle for many, perhaps most single-cell algae, and is clearly advantageous over both autotrophy or heterotrophy in communities limited by nutrients^55,56^. Thus, oligotrophic Proterozoic waters could have selected for the increased fitness that mixotrophy provided, but these nutrient-poor environments could not sustain a radiation of eukaryotic algae. The rise to ecological dominance of algae with red plastids during the Mesozoic might have been driven by profound environmental changes such as the increase in coastlines associated with the breakup of the supercontinent Pangea, providing newly flooded continental margins with high-nutrient habitats^57^. Importantly, these environmental changes would have happened after ∼1 Ga of evolution since the origin of red algal-derived plastids, providing ample opportunities for the evolution of a genetic toolkit that would prove beneficial with the rise of more favourable habitats. One such example of beneficial genetic innovation is cell protection by a variety of armour plating, which is a convergent feature of many ecologically successful algae with complex red plastids. These armours aid the phytoplankton to protect from grazing and thus represent an additional condition that may have favoured the late Mesozoic expansion to ecological dominance of some groups with red algal-derived plastids^57^.

### Conclusion

Algae powered by red algal-derived plastids are among the most evolutionary and ecologically successful eukaryotes on Earth. Yet, we still lack a comprehensive understanding of how, and how many times, red plastids were established. In recent years, hypotheses of serial endosymbiosis have flourished to explain how disparate groups of eukaryotes obtained their red plastids. In the present study, we used molecular clocks applied to a broad phylogenomic dataset to test whether the serial endosymbiosis hypotheses are chronologically possible. Our results indicate that all putative plastid donor and recipient lineages most likely overlapped during Earth history, thus in principle allowing plastids to be passed between distantly related hosts. Furthermore, we showed that the timeframe from the initial secondary endosymbiosis with a red alga to the establishment of all complex red plastids was relatively short, likely spanning between 650 to 1,079 million years mainly during the Mesoproterozoic Era. This relatively short timeframe represents a novel insight into the diversification of photosynthetic eukaryotes during the Mesoproterozoic and the origin of the most ecologically important modern-day algae. More generally, specific serial endosymbiosis hypotheses, if validated, will provide useful relative constraints for better understanding the overall timescale of eukaryote diversification in future paleobiological studies.

## Methods

### Phylogenomic dataset construction

Two publicly available phylogenomic datasets were used as starting points in this study: a 263 genes, 234 taxa dataset^26,35^, and a 351 genes, 64 taxa dataset^58^. Non-overlapping genes between these two datasets (134 genes) were identified by BLAST^59^ analyses, allowing the merging of both datasets to bring the total number of initial genes to 397. We expanded the sampling of species with publicly available genomic/transcriptomic data to obtain a comprehensive eukaryote-wide dataset, with particular attention to taxa most relevant to this study (sources: ensemblgenomes.org, imicrobe.us/#/projects/104, ncbi.nlm.nih.gov, onekp.com, and few publications that did not provide a link to a public sequence database; now all available in EukProt^60^). The procedure to add taxa was as follows: 1) For each taxon, protein sequences were clustered with CD-HIT^61^ using an identity threshold of 85%; 2) Homologous sequences were retrieved by BLASTP searches using all 397 genes as queries (e-value: 1e-20; coverage cutoff: 0.5); 3) In three rounds, gene trees were constructed and carefully inspected in order to detect and remove putative paralogs and contaminants. For that, sequences were aligned with MAFFT v. 7.310^62^ using either the -auto option (first round) or MAFFT L-INS-i with default settings (second and third round). Ambiguously aligned positions were filtered using trimAL v. 1.4^63^ with a gap threshold of 0.8 (all three rounds), followed by Maximum Likelihood (ML) single-gene tree reconstruction with either FastTree v. 2.1.10^64^ using -lg -gamma plus options for more accurate performances (first round) or RAxML v. 8.2.10^65^ with PROTGAMMALGF and 100 rapid bootstrap searches (second and third round). To facilitate the detection of contaminants and paralogs, all taxa were renamed following NCBI’s taxonomy (manually refined—the custom taxonomy is available in Supplementary Fig. 1; Supplementary Data 2), and colour-coded using in-house scripts. Multiple copies from the same taxon were assigned to a unique colour allowing to more easily detect contaminants and paralogs in FigTree v. 1.4.3 (http://tree.bio.ed.ac.uk/software/figtree/). After the three rounds of gene tree inspection, we discarded 77 genes (74 out of the 134 genes added from Brown *et al*.^58^) due to suspicious clustering of major groups (e.g., duplication of the entire Sar clade in FTSJ1). The resulting dataset comprised 320 genes and 733 eukaryote taxa with ≥5% data; Supplementary Data 2.

For each curated gene, sequence stretches without clear homology (e.g., poor quality stretches of amino acids, or leftover untranslated regions) were removed with PREQUAL v. 1.01^66^ employing a posterior probability threshold of 0.95 (ignoring some fast-evolving taxa). Sequences were aligned using MAFFT G-INS-i with a variable scoring matrix to avoid over-alignment (--unalignlevel 0.6) and trimmed with BMGE v. 1.12^67^ using -g 0.2, -b 5, -m BLOSUM75 parameters. Partial sequences belonging to the same taxon that did not show evidence for paralogy or contamination on the gene trees were merged. All 320 trimmed gene alignments were concatenated with SCaFos v. 1.25^68^ into a supermatrix of 733 taxa and 62,723 aligned amino acid positions (62,552 distinct patterns; gaps and undetermined characters: ∼35%; Supplementary Data 2). This dataset was subjected to ML analysis in IQ-TREE^69^ with the site-homogeneous model LG+G+F and ultrafast bootstrap approximation (UFBoot^70^; 1,000 replicates) employing the -bb and -bnni flags (IQ-TREE versions 1.6.3 to 1.6.9 have been used in this study). This large tree (Supplementary Fig. 1) was used to select a reduced taxon-sampling that maintained phylogenetic diversity but allowed downstream analyses with more sophisticated models. The taxa selection aimed to 1) retain all major lineages of eukaryotes; 2) preferentially keep slowly-evolving (shorter branches) representatives; 3) preferentially discard taxa with more missing data; and 4) allow the precise taxonomic placement of fossil calibrations. In order to increase sequence coverage, some monophyletic strains or species/genera complexes were combined to form a chimeric operational taxonomic unit (OTU; Supplementary Data 2). This strategy led to a dataset containing 136 OTUs, for which the raw sequences were again filtered, aligned, trimmed, and concatenated as described above, forming a supermatrix with 73,460 aligned amino acid positions (71,540 distinct patterns; gaps and undetermined characters: ∼17%; Supplementary Data 2).

### Phylogenomic analyses

The reduced dataset (136 OTUs) was subjected to ML analysis in IQ-TREE using the best-fit site-heterogeneous model LG+C60+G+F with the PMSF approach to calculate non-parametric bootstrap support (100 replicates; Supplementary Fig. 2). This dataset was also analysed under the Multi Species Coalescent (MSC) approach implemented in ASTRAL-III^71^ to account for incomplete lineage sorting (here, taxa were not combined into OTUs). In order to improve the phylogenetic signal in the single-gene trees used in ASTRAL, a partial filtering method (i.e., Divvier^72^ using the -partial option) was applied followed by trimming of highly incomplete positions with trimAL (-gt 0.05). ML single-gene trees were inferred with IQ-TREE under BIC-selected models including site-homogeneous models (such as LG) and empirical profile models (C10-C60). Branch support was inferred with 1,000 replicates of ultrafast bootstrap (-bbni) and branches with <10 support were collapsed. Branch support for the MSC tree (Supplementary Fig. 3) was calculated as quartet supports^73^, and multilocus bootstrapping (1,000 UFboot2 bootstraps per gene).

Finally, we created a taxon-subset of the 136-OTU dataset containing only 63 OTUs with 73,460 aligned amino acid positions (67,700 distinct patterns; gaps and undetermined characters: ∼13%; Supplementary Data 2), to be able to obtain chains convergence with the computationally very demanding CAT+GTR+G model in PhyloBayes-MPI v. 1.8. Three independent Markov Chain Monte Carlo (MCMC) chains were run for 2,600 cycles (all sampled). The initial 500 cycles were removed (as burnin) from each chain before generating a consensus tree using the bpcomp option. Global convergence was achieved in all combinations of the chains (Supplementary Fig. 4) with maxdiff reaching 0. The 63-OTU dataset was also used in posterior predictive analyses (Supplementary Table 1) to informatively select the best topology. For that, the CAT+GTR+G tree was compared to a tree obtained using PhyloBayes-MPI v. 1.8 under LG+C60+G+F; Supplementary Fig. 5). The reference topology used in the dating analyses corresponded to a ML analyses of the 136-OTU dataset under LG+C60+G+F but constrained by the relationships obtained in the CAT+GTR+G tree.

### Molecular dating

Bayesian molecular dating was performed with MCMCTree^74^ within the PAML package v. 4.9h^75^, PhyloBayes-MPI v. 1.8, and BEAST v. 1.10.4^76^. We used a total of 33 node calibrations based on fossil evidence (retrieved April 2019; Table 1) and the tree topology of Supplementary Fig. 4. MCMCTree was used to perform a set of sensitivity analyses (two chains for each experimental condition) in order to understand the effect of different clock models (uncorrelated or autocorrelated), tree roots, and prior calibration densities. A uniform birth-death tree prior was assumed and analyses were run with either (i) uncorrelated (clock = 2) or (ii) autocorrelated (clock = 3) relaxed clock models. The tree was rooted either on (i) Amorphea^77^ or (ii) Discoba (Excavata)^78^. Four prior calibration distributions were tested, following dos Reis *et al*.^31^: (a) uniform (i.e., maxima and minima), (b) skew-normal (α = 10 and β scale parameters chosen so that the 97.5% cumulative probabilities coincide with the maxima; calculated with MCMCTreeR https://github.com/PuttickMacroevolution/MCMCtreeR), and truncated-cauchy distributions with either (c) short (p = 0; c = 0.1; pL = 0.01) or (d) long (p = 0; c = 10; pL = 0.01) distribution tails. Skew-normal distributions represent ‘literal’ interpretations of the fossil record, assuming minima are close to the real node ages, whereas truncated-cauchy distributions represent more ‘loose’ interpretations that assume older divergences than minimum bounds^31^. In all cases, the root was calibrated using a uniform distribution 1.6–3.2 Ga. All maxima and minima were treated as soft bounds with a default 2.5% prior probability beyond their limits. MCMCTree analyses were run on the entire concatenated alignment using approximate likelihood calculations^79^. Data were analysed as a single partition under the LG+G model. The prior on the mean (or ancestral) rates (‘rgene_gamma’) were set as diffuse gamma Dirichlet priors indicating severe among-lineage rate heterogeneity and mean rates of 0.02625 and 0.0275 amino acid replacements site-1 10^8 Myr-1, for respectively the Amorphea (α = 2, β = 72.73) and Excavata (α = 2, β = 76.19) roots. The average rate was calculated as mean root-to-tip paths on the corresponding ML trees with the two roots. The rate drift parameter (‘sigma2_gamma’) was set to indicate considerable rate heterogeneity across lineages (α = 2, β = 2). A 100 million years time unit was assumed. Two independent MCMC chains were run for each analysis, each consisting of 20.2 million generations, of which the first 200,000 were excluded as burnin. Convergence of chains was checked *a posteriori* using Tracer v. 1.7.1^80^ and all parameters reached ESS > 200. A total of 32 analyses were run, corresponding to all possible combinations of two root positions, two clock models, four calibration distributions, and two MCMC chains per experimental condition.

For computational tractability, PhyloBayes and BEAST were run on a subset of the 10 most clock-like genes (selected with SortaDate^81^). PhyloBayes analyses were run under (i) uncorrelated and (ii) autocorrelated relaxed clock models and using either the (i) site-homogeneous LG+G or (ii) site-heterogeneous CAT+GTR models. In this case, calibrations were set as uniform priors with soft bounds and assumed a birth-death tree prior and the same tree topology (rooted on Amorphea). Two independent MCMC chains were run until convergence, assessed by PhyloBayes’ built-in tools (bpcomp and tracecomp). For comparative purposes, we run an additional BEAST analysis with similar parameterizations (among those available in BEAST): the uncorrelated relaxed clock, a fixed tree topology rooted on Amorphea, uniform prior calibrations, a Yule tree prior and the LG+G evolutionary model. Two independent MCMC chains were run for 200 million generations, the first 10% being discarded as burnin. Convergence of chains was checked with Tracer and all parameters reached ESS > 200. CorrTest^82^ was used to test the autocorrelation of branch lengths.

## Acknowledgments

This work was supported by a grant from Science for Life Laboratory available to FB, which covered the salary of JFHS and II. II was also supported by a postdoctoral Juan de la Cierva-Incorporación fellowship (IJCI-2016-29566) from the Spanish Ministry of Economy and Competitiveness (MINECO). TAW was supported by a Royal Society University Research Fellowship and NERC Grant NE/P00251X/1. Computations were performed on resources provided by the Swedish National Infrastructure for Computing (SNIC) at Uppsala Multidisciplinary Center for Advanced Computational Science (UPPMAX) under Projects 2017-7-65, 2017-7-355, 2018-3-147, 2018-3-288, 2018-8-187, 2018-8-192, and 2019-3-305.

## Author contributions

FB conceived, designed and supervised the project. JFHS assembled and curated the data, and performed phylogenomic analyses. TAW analysed data under the Multi Species Coalescent model. JFHS compiled the fossil records, and II designed and performed molecular dating analyses. Artwork was made by JFHS (Figs. 1, 2, S1–S6) and II (Figs. 3, S7). All authors drafted the manuscript and read and approved the final version.

## Additional information

Supplementary Information: Supplementary Figs. 1–7 and Supplementary Table 1

Supplementary Data 1: Molecular clock results and violin plots

Supplementary Data 2: Sequence data and taxonomy files

