## Supplementary Data 1 for "A molecular timescale for the origin of red algal-derived plastids": README.pdf

**Molecular_clock_results.xlsx** -> Results of divergence times from MCMCTree analyses (mean posterior, 95% HPD intervals) and comparison with Phylobayes and BEAST, including comparison of 10 key nodes as well as mean divergence times and 95% HPD interval widths.

**Violin_plots_all_nodes.pdf** -> Divergence times for all nodes under a broad range of conditions (MCMCTree analyses). Violin plots show the full posterior distribution. Mean ages are indicated with a white circle. For node assignment, see page 18. Note that the ages for node 138 of the tree rooted on Amorphea (RAM) do not correspond to node 138 of the tree rooted on Excavata (REX). Node 138 separates ingroup and outgroup, i.e., it refers to the common ancestor of all eukaryotes except Amorphea or Excavata for trees rooted on these groups, and therefore their ages are not comparable between differently rooted trees. All other node numbers in REX have been adjusted in order to facilitate comparison.
