## Supplementary Data 1 for "A molecular timescale for the origin of red algal-derived plastids": Violin_plots_all_nodes.pdf

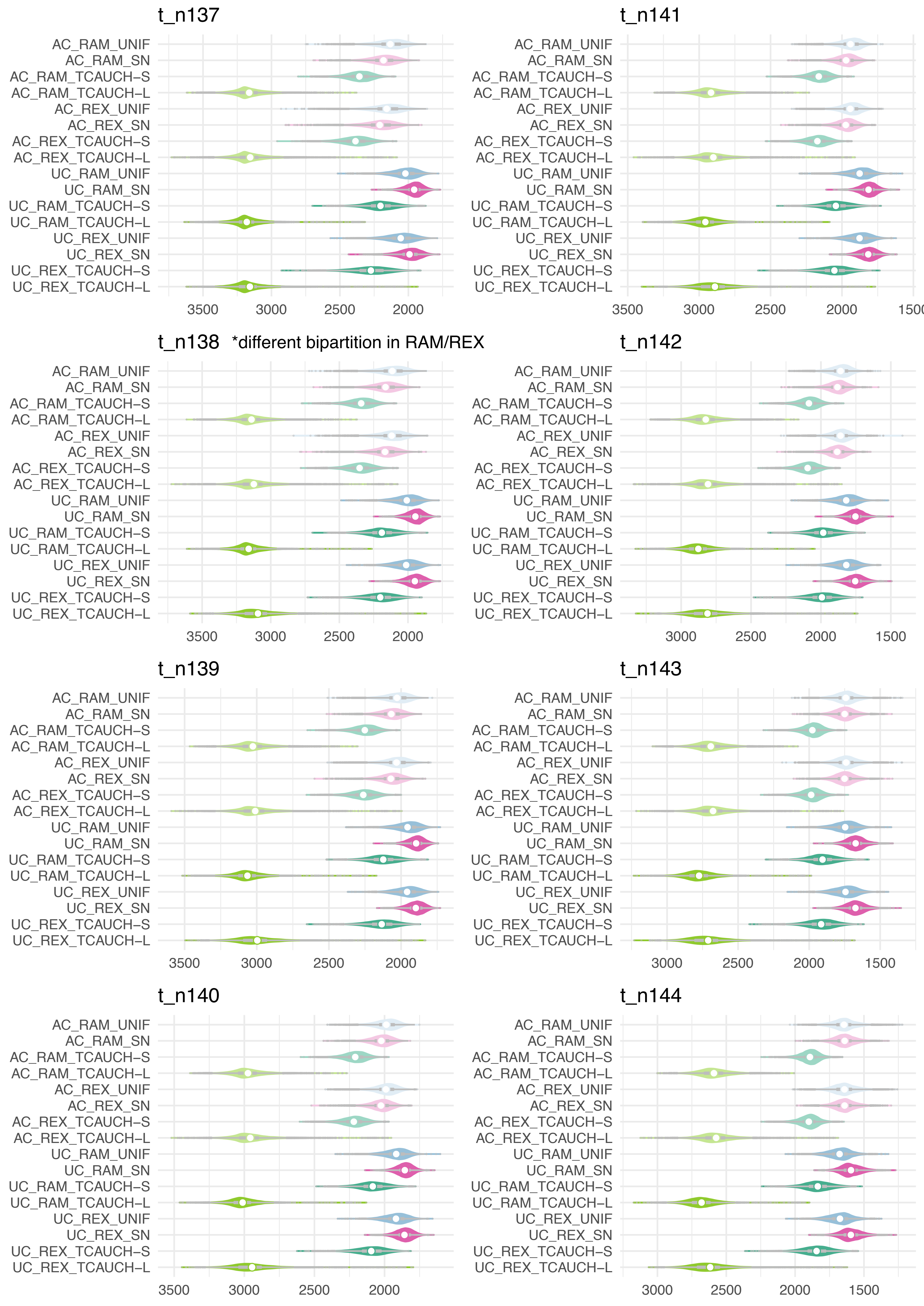

t_n137
AC_RAM_UNIF
AC_RAM_SN
AC_RAM_TCAUCH–S
AC_RAM_TCAUCH–L
AC_REX_UNIF
AC_REX_SN
AC_REX_TCAUCH–S
AC_REX_TCAUCH–L
UC_RAM_UNIF
UC_RAM_SN
UC_RAM_TCAUCH–S
UC_RAM_TCAUCH–L
UC_REX_UNIF
UC_REX_SN
UC_REX_TCAUCH–S
UC_REX_TCAUCH–L
3500
3000
2500
2000
t_n141
AC_RAM_UNIF
AC_RAM_SN
AC_RAM_TCAUCH–S
AC_RAM_TCAUCH–L
AC_REX_UNIF
AC_REX_SN
AC_REX_TCAUCH–S
AC_REX_TCAUCH–L
UC_RAM_UNIF
UC_RAM_SN
UC_RAM_TCAUCH–S
UC_RAM_TCAUCH–L
UC_REX_UNIF
UC_REX_SN
UC_REX_TCAUCH–S
UC_REX_TCAUCH–L
3500
3000
2500
2000
150
t_n138 *different bipartition in RAM/REX
AC_RAM_UNIF
AC_RAM_SN
AC_RAM_TCAUCH–S
AC_RAM_TCAUCH–L
AC_REX_UNIF
AC_REX_SN
AC_REX_TCAUCH–S
AC_REX_TCAUCH–L
UC_RAM_UNIF
UC_RAM_SN
UC_RAM_TCAUCH–S
UC_RAM_TCAUCH–L
UC_REX_UNIF
UC_REX_SN
UC_REX_TCAUCH–S
UC_REX_TCAUCH–L
3500
3000
2500
2000
t_n142
AC_RAM_UNIF
AC_RAM_SN
AC_RAM_TCAUCH–S
AC_RAM_TCAUCH–L
AC_REX_UNIF
AC_REX_SN
AC_REX_TCAUCH–S
AC_REX_TCAUCH–L
UC_RAM_UNIF
UC_RAM_SN
UC_RAM_TCAUCH–S
UC_RAM_TCAUCH–L
UC_REX_UNIF
UC_REX_SN
UC_REX_TCAUCH–S
UC_REX_TCAUCH–L
3000
2500
2000
1500
t_n139
AC_RAM_UNIF
AC_RAM_SN
AC_RAM_TCAUCH–S
AC_RAM_TCAUCH–L
AC_REX_UNIF
AC_REX_SN
AC_REX_TCAUCH–S
AC_REX_TCAUCH–L
UC_RAM_UNIF
UC_RAM_SN
UC_RAM_TCAUCH–S
UC_RAM_TCAUCH–L
UC_REX_UNIF
UC_REX_SN
UC_REX_TCAUCH–S
UC_REX_TCAUCH–L
3500
3000
2500
2000
t_n143
AC_RAM_UNIF
AC_RAM_SN
AC_RAM_TCAUCH–S
AC_RAM_TCAUCH–L
AC_REX_UNIF
AC_REX_SN
AC_REX_TCAUCH–S
AC_REX_TCAUCH–L
UC_RAM_UNIF
UC_RAM_SN
UC_RAM_TCAUCH–S
UC_RAM_TCAUCH–L
UC_REX_UNIF
UC_REX_SN
UC_REX_TCAUCH–S
UC_REX_TCAUCH–L
3000
2500
2000
1500
t_n140
AC_RAM_UNIF
AC_RAM_SN
AC_RAM_TCAUCH–S
AC_RAM_TCAUCH–L
AC_REX_UNIF
AC_REX_SN
AC_REX_TCAUCH–S
AC_REX_TCAUCH–L
UC_RAM_UNIF
UC_RAM_SN
UC_RAM_TCAUCH–S
UC_RAM_TCAUCH–L
UC_REX_UNIF
UC_REX_SN
UC_REX_TCAUCH–S
UC_REX_TCAUCH–L
3500
3000
2500
2000
t_n144
AC_RAM_UNIF
AC_RAM_SN
AC_RAM_TCAUCH–S
AC_RAM_TCAUCH–L
AC_REX_UNIF
AC_REX_SN
AC_REX_TCAUCH–S
AC_REX_TCAUCH–L
UC_RAM_UNIF
UC_RAM_SN
UC_RAM_TCAUCH–S
UC_RAM_TCAUCH–L
UC_REX_UNIF
UC_REX_SN
UC_REX_TCAUCH–S
UC_REX_TCAUCH–L
3000
2500
2000
1500

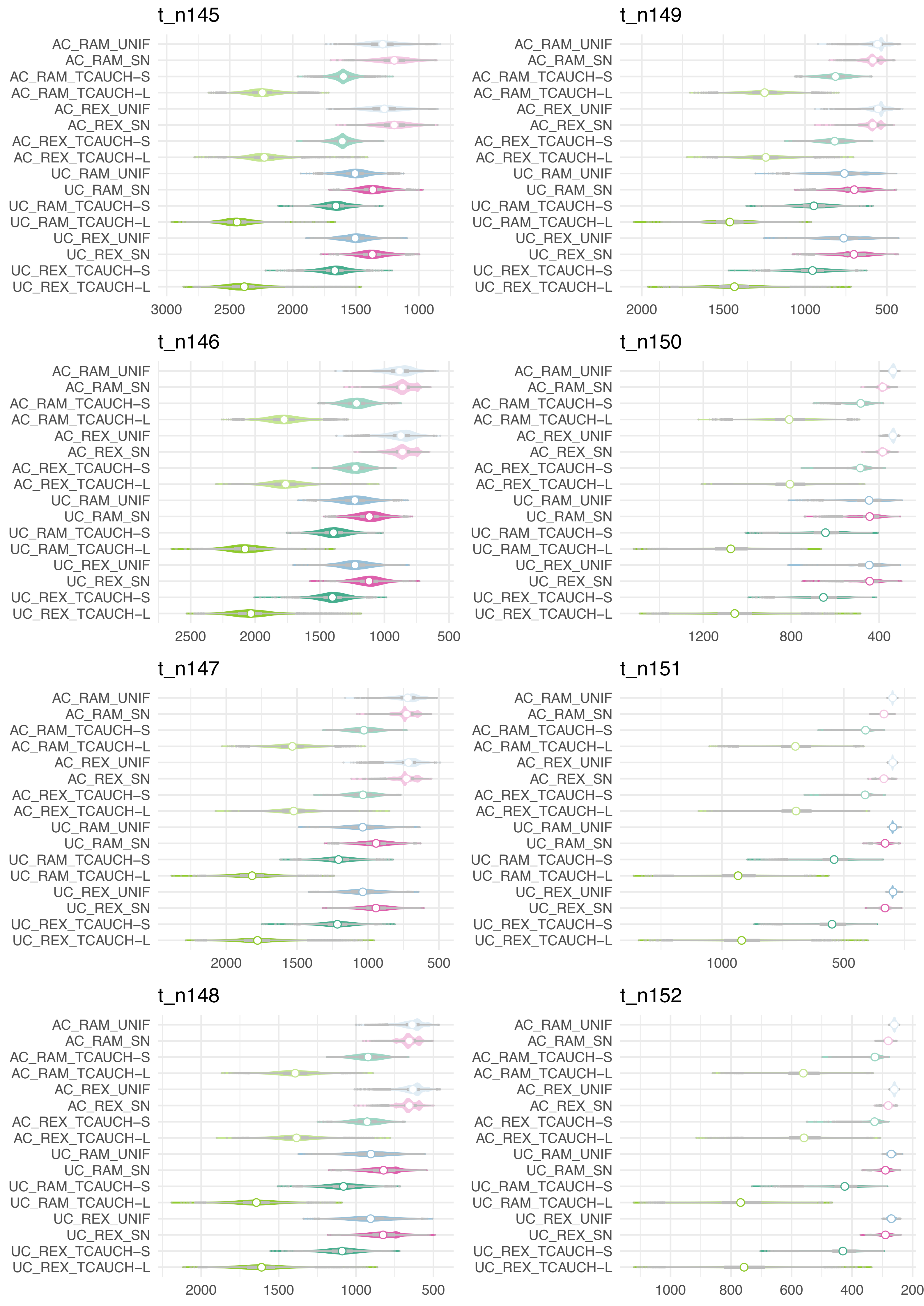
t_n145
AC_RAM_UNIF
AC_RAM_SN
AC_RAM_TCAUCH–S
AC_RAM_TCAUCH–L
AC_REX_UNIF
AC_REX_SN
AC_REX_TCAUCH–S
AC_REX_TCAUCH–L
UC_RAM_UNIF
UC_RAM_SN
UC_RAM_TCAUCH–S
UC_RAM_TCAUCH–L
UC_REX_UNIF
UC_REX_SN
UC_REX_TCAUCH–S
UC_REX_TCAUCH–L
3000
2500
2000
1500
1000
t_n146
AC_RAM_UNIF
AC_RAM_SN
AC_RAM_TCAUCH–S
AC_RAM_TCAUCH–L
AC_REX_UNIF
AC_REX_SN
AC_REX_TCAUCH–S
AC_REX_TCAUCH–L
UC_RAM_UNIF
UC_RAM_SN
UC_RAM_TCAUCH–S
UC_RAM_TCAUCH–L
UC_REX_UNIF
UC_REX_SN
UC_REX_TCAUCH–S
UC_REX_TCAUCH–L
2500
2000
1500
1000
500
t_n147
AC_RAM_UNIF
AC_RAM_SN
AC_RAM_TCAUCH–S
AC_RAM_TCAUCH–L
AC_REX_UNIF
AC_REX_SN
AC_REX_TCAUCH–S
AC_REX_TCAUCH–L
UC_RAM_UNIF
UC_RAM_SN
UC_RAM_TCAUCH–S
UC_RAM_TCAUCH–L
UC_REX_UNIF
UC_REX_SN
UC_REX_TCAUCH–S
UC_REX_TCAUCH–L
2000
1500
1000
500
t_n148
AC_RAM_UNIF
AC_RAM_SN
AC_RAM_TCAUCH–S
AC_RAM_TCAUCH–L
AC_REX_UNIF
AC_REX_SN
AC_REX_TCAUCH–S
AC_REX_TCAUCH–L
UC_RAM_UNIF
UC_RAM_SN
UC_RAM_TCAUCH–S
UC_RAM_TCAUCH–L
UC_REX_UNIF
UC_REX_SN
UC_REX_TCAUCH–S
UC_REX_TCAUCH–L
2000
1500
1000
500
t_n149
AC_RAM_UNIF
AC_RAM_SN
AC_RAM_TCAUCH–S
AC_RAM_TCAUCH–L
AC_REX_UNIF
AC_REX_SN
AC_REX_TCAUCH–S
AC_REX_TCAUCH–L
UC_RAM_UNIF
UC_RAM_SN
UC_RAM_TCAUCH–S
UC_RAM_TCAUCH–L
UC_REX_UNIF
UC_REX_SN
UC_REX_TCAUCH–S
UC_REX_TCAUCH–L
2000
1500
1000
500
t_n150
AC_RAM_UNIF
AC_RAM_SN
AC_RAM_TCAUCH–S
AC_RAM_TCAUCH–L
AC_REX_UNIF
AC_REX_SN
AC_REX_TCAUCH–S
AC_REX_TCAUCH–L
UC_RAM_UNIF
UC_RAM_SN
UC_RAM_TCAUCH–S
UC_RAM_TCAUCH–L
UC_REX_UNIF
UC_REX_SN
UC_REX_TCAUCH–S
UC_REX_TCAUCH–L
1200
800
400
t_n151
AC_RAM_UNIF
AC_RAM_SN
AC_RAM_TCAUCH–S
AC_RAM_TCAUCH–L
AC_REX_UNIF
AC_REX_SN
AC_REX_TCAUCH–S
AC_REX_TCAUCH–L
UC_RAM_UNIF
UC_RAM_SN
UC_RAM_TCAUCH–S
UC_RAM_TCAUCH–L
UC_REX_UNIF
UC_REX_SN
UC_REX_TCAUCH–S
UC_REX_TCAUCH–L
1000
500
t_n152
AC_RAM_UNIF
AC_RAM_SN
AC_RAM_TCAUCH–S
AC_RAM_TCAUCH–L
AC_REX_UNIF
AC_REX_SN
AC_REX_TCAUCH–S
AC_REX_TCAUCH–L
UC_RAM_UNIF
UC_RAM_SN
UC_RAM_TCAUCH–S
UC_RAM_TCAUCH–L
UC_REX_UNIF
UC_REX_SN
UC_REX_TCAUCH–S
UC_REX_TCAUCH–L
1000
800
600
400
200

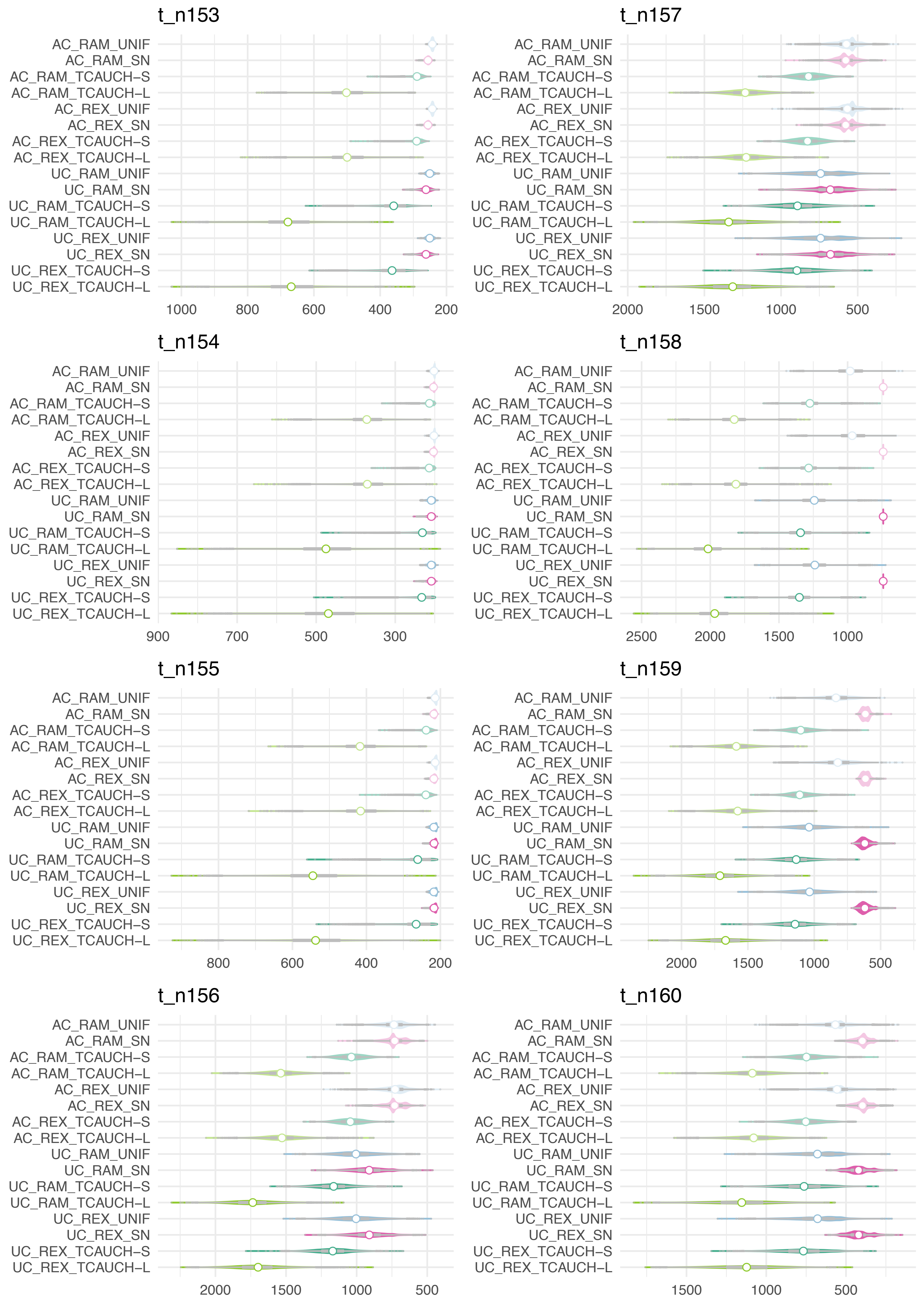

t_n153
AC_RAM_UNIF
AC_RAM_SN
AC_RAM_TCAUCH-S
AC_RAM_TCAUCH-L
AC_REX_UNIF
AC_REX_SN
AC_REX_TCAUCH-S
AC_REX_TCAUCH-L
UC_RAM_UNIF
UC_RAM_SN
UC_RAM_TCAUCH-S
UC_RAM_TCAUCH-L
UC_REX_UNIF
UC_REX_SN
UC_REX_TCAUCH-S
UC_REX_TCAUCH-L
1000
800
600
400
200
t_n154
AC_RAM_UNIF
AC_RAM_SN
AC_RAM_TCAUCH-S
AC_RAM_TCAUCH-L
AC_REX_UNIF
AC_REX_SN
AC_REX_TCAUCH-S
AC_REX_TCAUCH-L
UC_RAM_UNIF
UC_RAM_SN
UC_RAM_TCAUCH-S
UC_RAM_TCAUCH-L
UC_REX_UNIF
UC_REX_SN
UC_REX_TCAUCH-S
UC_REX_TCAUCH-L
900
700
500
300
t_n155
AC_RAM_UNIF
AC_RAM_SN
AC_RAM_TCAUCH-S
AC_RAM_TCAUCH-L
AC_REX_UNIF
AC_REX_SN
AC_REX_TCAUCH-S
AC_REX_TCAUCH-L
UC_RAM_UNIF
UC_RAM_SN
UC_RAM_TCAUCH-S
UC_RAM_TCAUCH-L
UC_REX_UNIF
UC_REX_SN
UC_REX_TCAUCH-S
UC_REX_TCAUCH-L
800
600
400
200
t_n156
AC_RAM_UNIF
AC_RAM_SN
AC_RAM_TCAUCH-S
AC_RAM_TCAUCH-L
AC_REX_UNIF
AC_REX_SN
AC_REX_TCAUCH-S
AC_REX_TCAUCH-L
UC_RAM_UNIF
UC_RAM_SN
UC_RAM_TCAUCH-S
UC_RAM_TCAUCH-L
UC_REX_UNIF
UC_REX_SN
UC_REX_TCAUCH-S
UC_REX_TCAUCH-L
2000
1500
1000
500
t_n157
AC_RAM_UNIF
AC_RAM_SN
AC_RAM_TCAUCH-S
AC_RAM_TCAUCH-L
AC_REX_UNIF
AC_REX_SN
AC_REX_TCAUCH-S
AC_REX_TCAUCH-L
UC_RAM_UNIF
UC_RAM_SN
UC_RAM_TCAUCH-S
UC_RAM_TCAUCH-L
UC_REX_UNIF
UC_REX_SN
UC_REX_TCAUCH-S
UC_REX_TCAUCH-L
2000
1500
1000
500
t_n158
AC_RAM_UNIF
AC_RAM_SN
AC_RAM_TCAUCH-S
AC_RAM_TCAUCH-L
AC_REX_UNIF
AC_REX_SN
AC_REX_TCAUCH-S
AC_REX_TCAUCH-L
UC_RAM_UNIF
UC_RAM_SN
UC_RAM_TCAUCH-S
UC_RAM_TCAUCH-L
UC_REX_UNIF
UC_REX_SN
UC_REX_TCAUCH-S
UC_REX_TCAUCH-L
2500
2000
1500
1000
t_n159
AC_RAM_UNIF
AC_RAM_SN
AC_RAM_TCAUCH-S
AC_RAM_TCAUCH-L
AC_REX_UNIF
AC_REX_SN
AC_REX_TCAUCH-S
AC_REX_TCAUCH-L
UC_RAM_UNIF
UC_RAM_SN
UC_RAM_TCAUCH-S
UC_RAM_TCAUCH-L
UC_REX_UNIF
UC_REX_SN
UC_REX_TCAUCH-S
UC_REX_TCAUCH-L
2000
1500
1000
500
t_n160
AC_RAM_UNIF
AC_RAM_SN
AC_RAM_TCAUCH-S
AC_RAM_TCAUCH-L
AC_REX_UNIF
AC_REX_SN
AC_REX_TCAUCH-S
AC_REX_TCAUCH-L
UC_RAM_UNIF
UC_RAM_SN
UC_RAM_TCAUCH-S
UC_RAM_TCAUCH-L
UC_REX_UNIF
UC_REX_SN
UC_REX_TCAUCH-S
UC_REX_TCAUCH-L
1500
1000
500

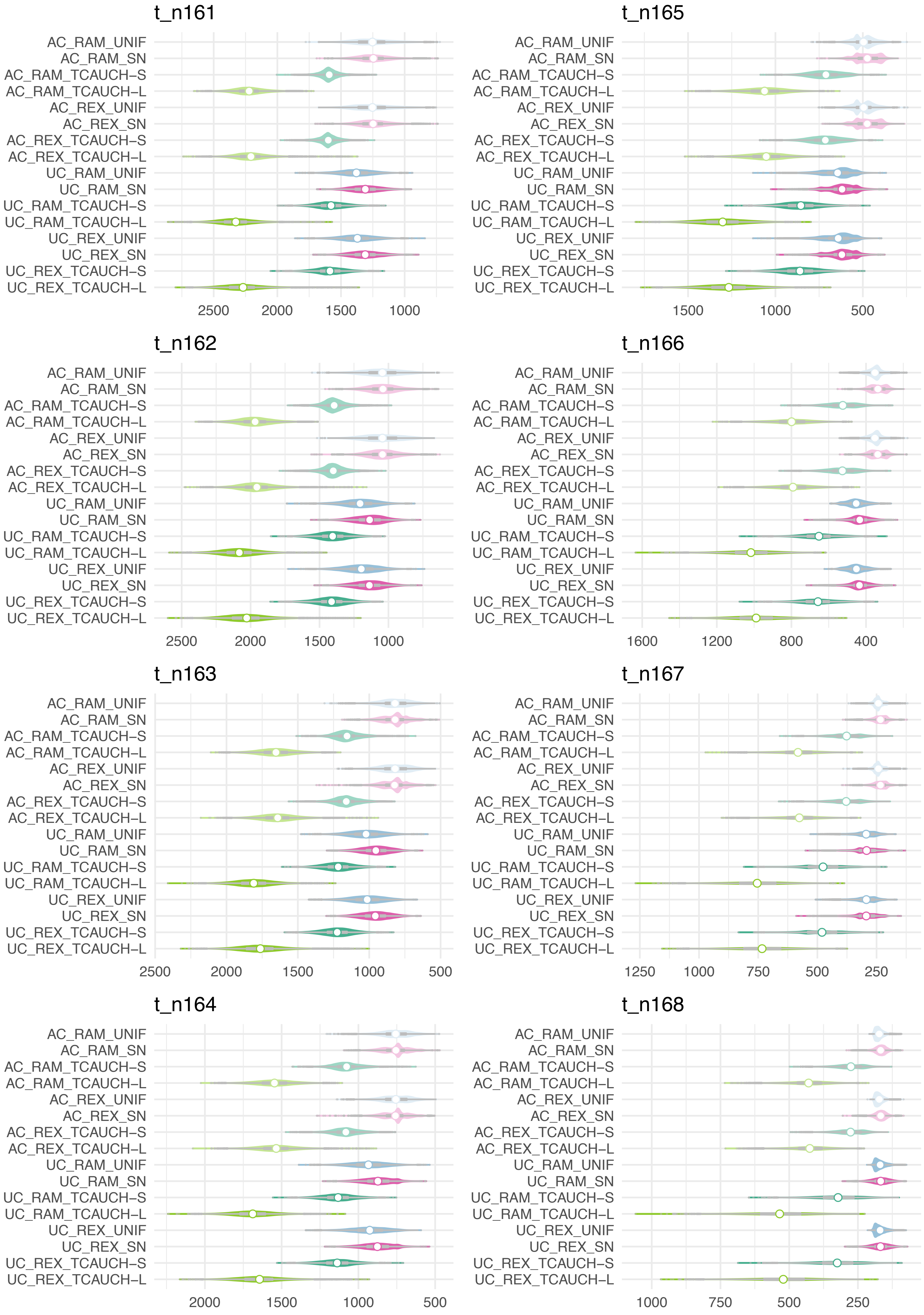
t_n161
t_n165
t_n162
t_n166
t_n163
t_n167
t_n164
t_n168
AC_RAM_UNIF
AC_RAM_SN
AC_RAM_TCAUCH–S
AC_RAM_TCAUCH–L
AC_REX_UNIF
AC_REX_SN
AC_REX_TCAUCH–S
AC_REX_TCAUCH–L
UC_RAM_UNIF
UC_RAM_SN
UC_RAM_TCAUCH–S
UC_RAM_TCAUCH–L
UC_REX_UNIF
UC_REX_SN
UC_REX_TCAUCH–S
UC_REX_TCAUCH–L
2500 2000 1500 1000
1500 1000 500
2500 2000 1500 1000
1600 1200 800 400
2500 2000 1500 1000 500
1250 1000 750 500 250
2000 1500 1000 500
1000 750 500 250

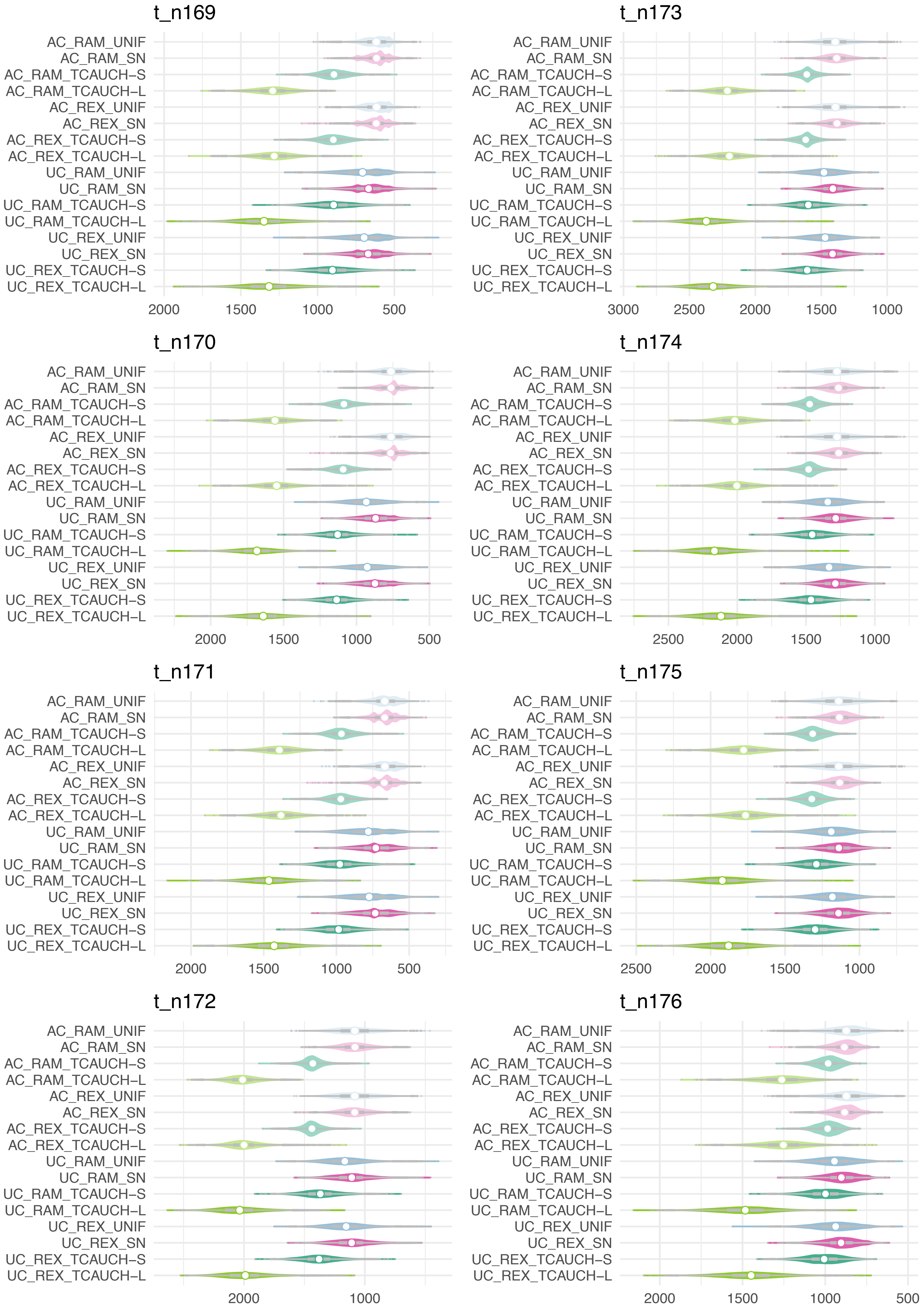

t_n169
AC_RAM_UNIF
AC_RAM_SN
AC_RAM_TCAUCH-S
AC_RAM_TCAUCH-L
AC_REX_UNIF
AC_REX_SN
AC_REX_TCAUCH-S
AC_REX_TCAUCH-L
UC_RAM_UNIF
UC_RAM_SN
UC_RAM_TCAUCH-S
UC_RAM_TCAUCH-L
UC_REX_UNIF
UC_REX_SN
UC_REX_TCAUCH-S
UC_REX_TCAUCH-L
2000
1500
1000
500
t_n173
AC_RAM_UNIF
AC_RAM_SN
AC_RAM_TCAUCH-S
AC_RAM_TCAUCH-L
AC_REX_UNIF
AC_REX_SN
AC_REX_TCAUCH-S
AC_REX_TCAUCH-L
UC_RAM_UNIF
UC_RAM_SN
UC_RAM_TCAUCH-S
UC_RAM_TCAUCH-L
UC_REX_UNIF
UC_REX_SN
UC_REX_TCAUCH-S
UC_REX_TCAUCH-L
3000
2500
2000
1500
1000
t_n170
AC_RAM_UNIF
AC_RAM_SN
AC_RAM_TCAUCH-S
AC_RAM_TCAUCH-L
AC_REX_UNIF
AC_REX_SN
AC_REX_TCAUCH-S
AC_REX_TCAUCH-L
UC_RAM_UNIF
UC_RAM_SN
UC_RAM_TCAUCH-S
UC_RAM_TCAUCH-L
UC_REX_UNIF
UC_REX_SN
UC_REX_TCAUCH-S
UC_REX_TCAUCH-L
2000
1500
1000
500
t_n174
AC_RAM_UNIF
AC_RAM_SN
AC_RAM_TCAUCH-S
AC_RAM_TCAUCH-L
AC_REX_UNIF
AC_REX_SN
AC_REX_TCAUCH-S
AC_REX_TCAUCH-L
UC_RAM_UNIF
UC_RAM_SN
UC_RAM_TCAUCH-S
UC_RAM_TCAUCH-L
UC_REX_UNIF
UC_REX_SN
UC_REX_TCAUCH-S
UC_REX_TCAUCH-L
2500
2000
1500
1000
t_n171
AC_RAM_UNIF
AC_RAM_SN
AC_RAM_TCAUCH-S
AC_RAM_TCAUCH-L
AC_REX_UNIF
AC_REX_SN
AC_REX_TCAUCH-S
AC_REX_TCAUCH-L
UC_RAM_UNIF
UC_RAM_SN
UC_RAM_TCAUCH-S
UC_RAM_TCAUCH-L
UC_REX_UNIF
UC_REX_SN
UC_REX_TCAUCH-S
UC_REX_TCAUCH-L
2000
1500
1000
500
t_n175
AC_RAM_UNIF
AC_RAM_SN
AC_RAM_TCAUCH-S
AC_RAM_TCAUCH-L
AC_REX_UNIF
AC_REX_SN
AC_REX_TCAUCH-S
AC_REX_TCAUCH-L
UC_RAM_UNIF
UC_RAM_SN
UC_RAM_TCAUCH-S
UC_RAM_TCAUCH-L
UC_REX_UNIF
UC_REX_SN
UC_REX_TCAUCH-S
UC_REX_TCAUCH-L
2500
2000
1500
1000
t_n172
AC_RAM_UNIF
AC_RAM_SN
AC_RAM_TCAUCH-S
AC_RAM_TCAUCH-L
AC_REX_UNIF
AC_REX_SN
AC_REX_TCAUCH-S
AC_REX_TCAUCH-L
UC_RAM_UNIF
UC_RAM_SN
UC_RAM_TCAUCH-S
UC_RAM_TCAUCH-L
UC_REX_UNIF
UC_REX_SN
UC_REX_TCAUCH-S
UC_REX_TCAUCH-L
2000
1000
t_n176
AC_RAM_UNIF
AC_RAM_SN
AC_RAM_TCAUCH-S
AC_RAM_TCAUCH-L
AC_REX_UNIF
AC_REX_SN
AC_REX_TCAUCH-S
AC_REX_TCAUCH-L
UC_RAM_UNIF
UC_RAM_SN
UC_RAM_TCAUCH-S
UC_RAM_TCAUCH-L
UC_REX_UNIF
UC_REX_SN
UC_REX_TCAUCH-S
UC_REX_TCAUCH-L
2000
1500
1000
500

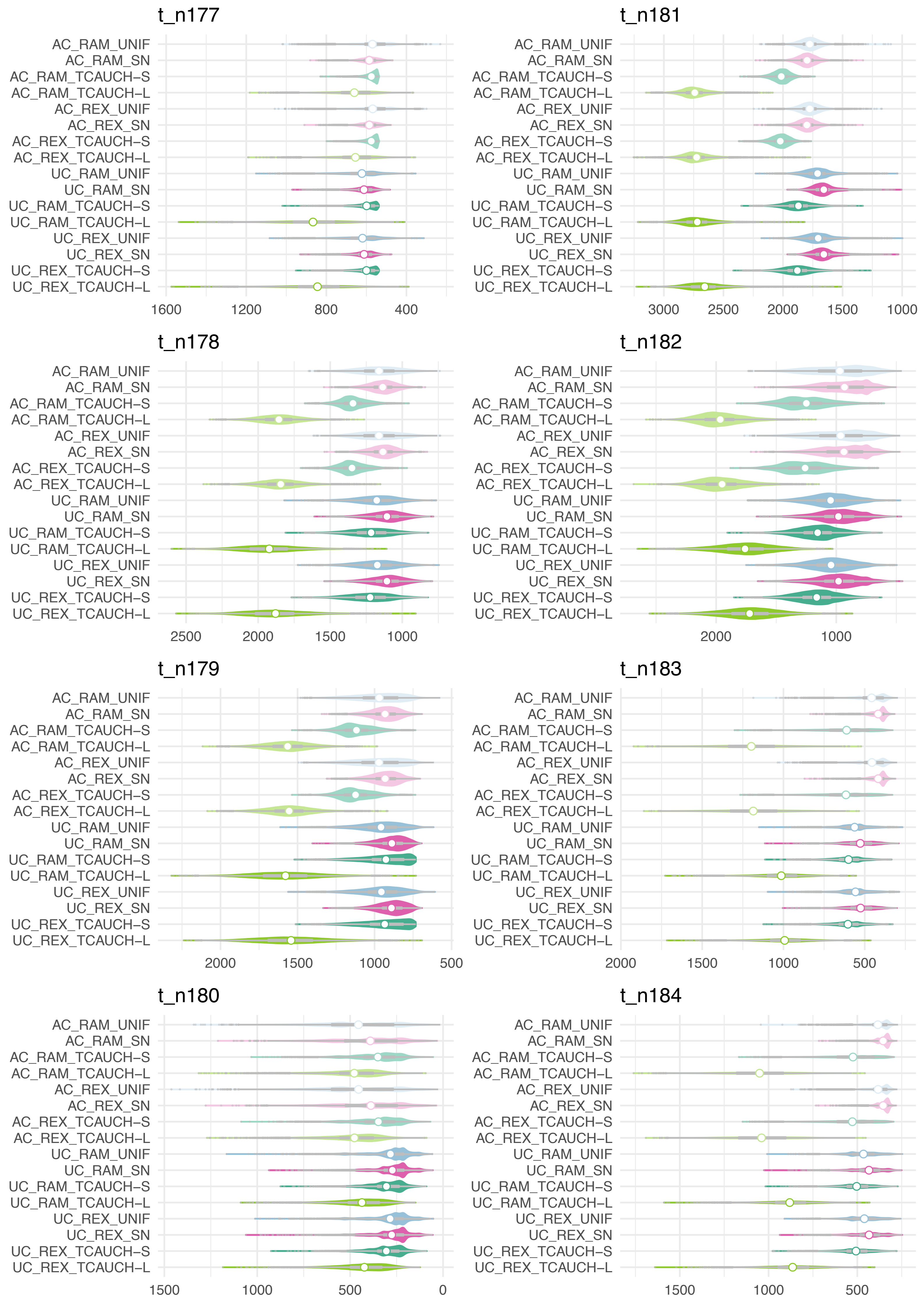
t_n177
AC_RAM_UNIF
AC_RAM_SN
AC_RAM_TCAUCH–S
AC_RAM_TCAUCH–L
AC_REX_UNIF
AC_REX_SN
AC_REX_TCAUCH–S
AC_REX_TCAUCH–L
UC_RAM_UNIF
UC_RAM_SN
UC_RAM_TCAUCH–S
UC_RAM_TCAUCH–L
UC_REX_UNIF
UC_REX_SN
UC_REX_TCAUCH–S
UC_REX_TCAUCH–L
1600
1200
800
400
t_n178
AC_RAM_UNIF
AC_RAM_SN
AC_RAM_TCAUCH–S
AC_RAM_TCAUCH–L
AC_REX_UNIF
AC_REX_SN
AC_REX_TCAUCH–S
AC_REX_TCAUCH–L
UC_RAM_UNIF
UC_RAM_SN
UC_RAM_TCAUCH–S
UC_RAM_TCAUCH–L
UC_REX_UNIF
UC_REX_SN
UC_REX_TCAUCH–S
UC_REX_TCAUCH–L
2500
2000
1500
1000
t_n179
AC_RAM_UNIF
AC_RAM_SN
AC_RAM_TCAUCH–S
AC_RAM_TCAUCH–L
AC_REX_UNIF
AC_REX_SN
AC_REX_TCAUCH–S
AC_REX_TCAUCH–L
UC_RAM_UNIF
UC_RAM_SN
UC_RAM_TCAUCH–S
UC_RAM_TCAUCH–L
UC_REX_UNIF
UC_REX_SN
UC_REX_TCAUCH–S
UC_REX_TCAUCH–L
2000
1500
1000
500
t_n180
AC_RAM_UNIF
AC_RAM_SN
AC_RAM_TCAUCH–S
AC_RAM_TCAUCH–L
AC_REX_UNIF
AC_REX_SN
AC_REX_TCAUCH–S
AC_REX_TCAUCH–L
UC_RAM_UNIF
UC_RAM_SN
UC_RAM_TCAUCH–S
UC_RAM_TCAUCH–L
UC_REX_UNIF
UC_REX_SN
UC_REX_TCAUCH–S
UC_REX_TCAUCH–L
1500
1000
500
0
t_n181
AC_RAM_UNIF
AC_RAM_SN
AC_RAM_TCAUCH–S
AC_RAM_TCAUCH–L
AC_REX_UNIF
AC_REX_SN
AC_REX_TCAUCH–S
AC_REX_TCAUCH–L
UC_RAM_UNIF
UC_RAM_SN
UC_RAM_TCAUCH–S
UC_RAM_TCAUCH–L
UC_REX_UNIF
UC_REX_SN
UC_REX_TCAUCH–S
UC_REX_TCAUCH–L
3000
2500
2000
1500
1000
t_n182
AC_RAM_UNIF
AC_RAM_SN
AC_RAM_TCAUCH–S
AC_RAM_TCAUCH–L
AC_REX_UNIF
AC_REX_SN
AC_REX_TCAUCH–S
AC_REX_TCAUCH–L
UC_RAM_UNIF
UC_RAM_SN
UC_RAM_TCAUCH–S
UC_RAM_TCAUCH–L
UC_REX_UNIF
UC_REX_SN
UC_REX_TCAUCH–S
UC_REX_TCAUCH–L
2000
1000
t_n183
AC_RAM_UNIF
AC_RAM_SN
AC_RAM_TCAUCH–S
AC_RAM_TCAUCH–L
AC_REX_UNIF
AC_REX_SN
AC_REX_TCAUCH–S
AC_REX_TCAUCH–L
UC_RAM_UNIF
UC_RAM_SN
UC_RAM_TCAUCH–S
UC_RAM_TCAUCH–L
UC_REX_UNIF
UC_REX_SN
UC_REX_TCAUCH–S
UC_REX_TCAUCH–L
2000
1500
1000
500
t_n184
AC_RAM_UNIF
AC_RAM_SN
AC_RAM_TCAUCH–S
AC_RAM_TCAUCH–L
AC_REX_UNIF
AC_REX_SN
AC_REX_TCAUCH–S
AC_REX_TCAUCH–L
UC_RAM_UNIF
UC_RAM_SN
UC_RAM_TCAUCH–S
UC_RAM_TCAUCH–L
UC_REX_UNIF
UC_REX_SN
UC_REX_TCAUCH–S
UC_REX_TCAUCH–L
1500
1000
500

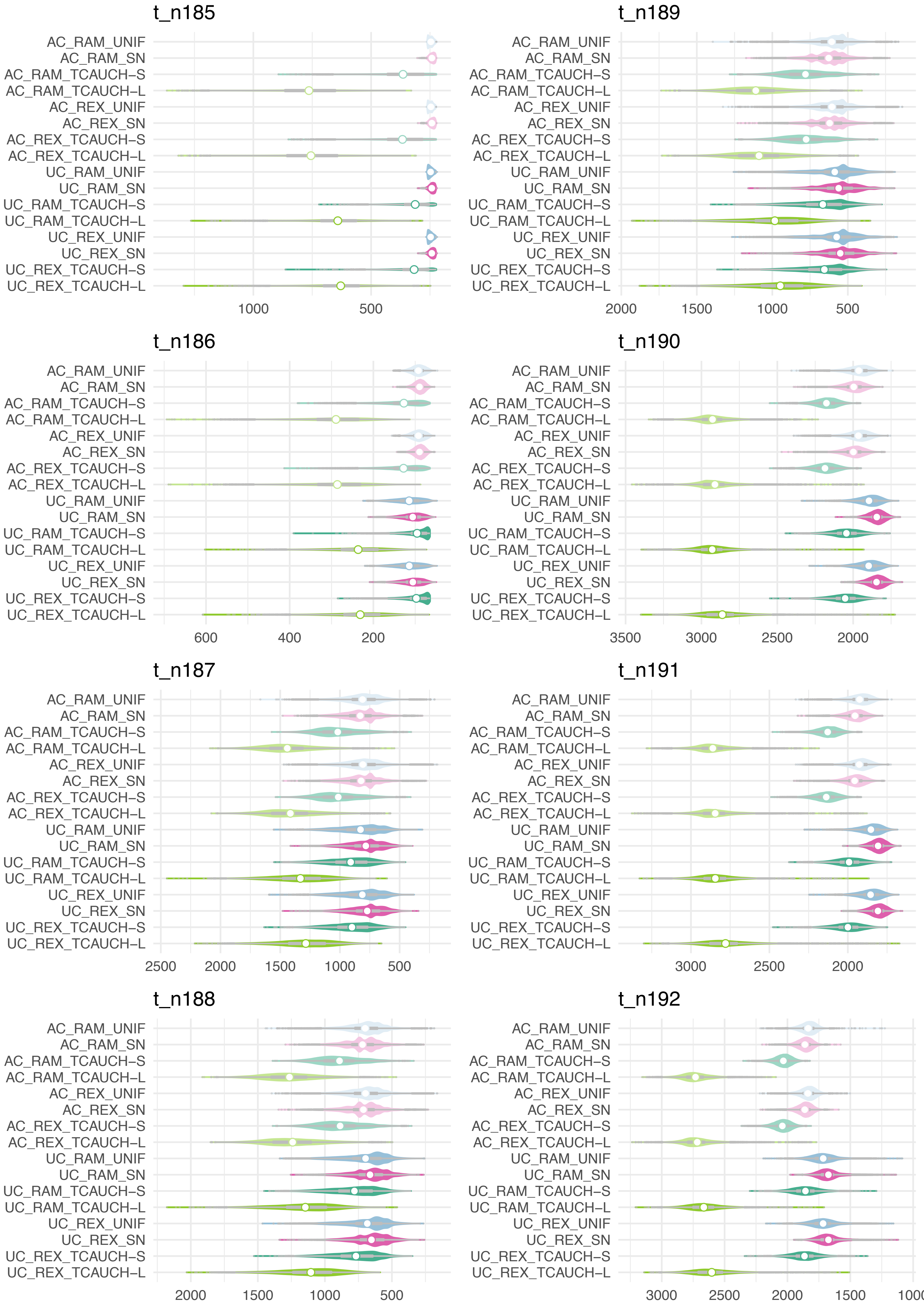

t_n185
t_n189
t_n186
t_n190
t_n187
t_n191
t_n188
t_n192
AC_RAM_UNIF
AC_RAM_SN
AC_RAM_TCAUCH–S
AC_RAM_TCAUCH–L
AC_REX_UNIF
AC_REX_SN
AC_REX_TCAUCH–S
AC_REX_TCAUCH–L
UC_RAM_UNIF
UC_RAM_SN
UC_RAM_TCAUCH–S
UC_RAM_TCAUCH–L
UC_REX_UNIF
UC_REX_SN
UC_REX_TCAUCH–S
UC_REX_TCAUCH–L
1000 500
2000 1500 1000 500
600 400 200
3500 3000 2500 2000
2500 2000 1500 1000 500
3000 2500 2000
2000 1500 1000 500
3000 2500 2000 1500 1000

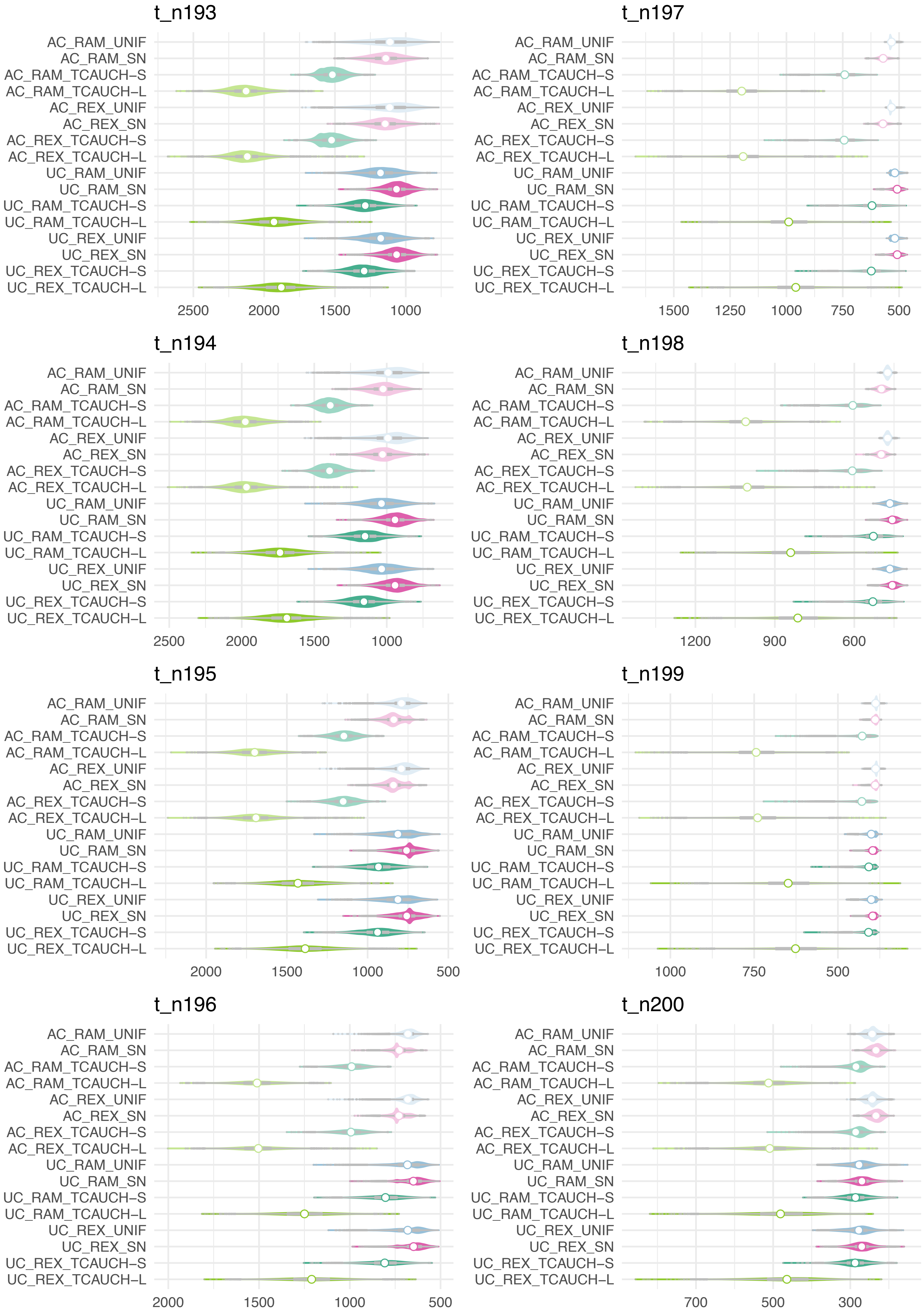
t_n193
AC_RAM_UNIF
AC_RAM_SN
AC_RAM_TCAUCH–S
AC_RAM_TCAUCH–L
AC_REX_UNIF
AC_REX_SN
AC_REX_TCAUCH–S
AC_REX_TCAUCH–L
UC_RAM_UNIF
UC_RAM_SN
UC_RAM_TCAUCH–S
UC_RAM_TCAUCH–L
UC_REX_UNIF
UC_REX_SN
UC_REX_TCAUCH–S
UC_REX_TCAUCH–L
2500
2000
1500
1000
t_n197
AC_RAM_UNIF
AC_RAM_SN
AC_RAM_TCAUCH–S
AC_RAM_TCAUCH–L
AC_REX_UNIF
AC_REX_SN
AC_REX_TCAUCH–S
AC_REX_TCAUCH–L
UC_RAM_UNIF
UC_RAM_SN
UC_RAM_TCAUCH–S
UC_RAM_TCAUCH–L
UC_REX_UNIF
UC_REX_SN
UC_REX_TCAUCH–S
UC_REX_TCAUCH–L
1500
1250
1000
750
500
t_n194
AC_RAM_UNIF
AC_RAM_SN
AC_RAM_TCAUCH–S
AC_RAM_TCAUCH–L
AC_REX_UNIF
AC_REX_SN
AC_REX_TCAUCH–S
AC_REX_TCAUCH–L
UC_RAM_UNIF
UC_RAM_SN
UC_RAM_TCAUCH–S
UC_RAM_TCAUCH–L
UC_REX_UNIF
UC_REX_SN
UC_REX_TCAUCH–S
UC_REX_TCAUCH–L
2500
2000
1500
1000
t_n198
AC_RAM_UNIF
AC_RAM_SN
AC_RAM_TCAUCH–S
AC_RAM_TCAUCH–L
AC_REX_UNIF
AC_REX_SN
AC_REX_TCAUCH–S
AC_REX_TCAUCH–L
UC_RAM_UNIF
UC_RAM_SN
UC_RAM_TCAUCH–S
UC_RAM_TCAUCH–L
UC_REX_UNIF
UC_REX_SN
UC_REX_TCAUCH–S
UC_REX_TCAUCH–L
1200
900
600
t_n195
AC_RAM_UNIF
AC_RAM_SN
AC_RAM_TCAUCH–S
AC_RAM_TCAUCH–L
AC_REX_UNIF
AC_REX_SN
AC_REX_TCAUCH–S
AC_REX_TCAUCH–L
UC_RAM_UNIF
UC_RAM_SN
UC_RAM_TCAUCH–S
UC_RAM_TCAUCH–L
UC_REX_UNIF
UC_REX_SN
UC_REX_TCAUCH–S
UC_REX_TCAUCH–L
2000
1500
1000
500
t_n199
AC_RAM_UNIF
AC_RAM_SN
AC_RAM_TCAUCH–S
AC_RAM_TCAUCH–L
AC_REX_UNIF
AC_REX_SN
AC_REX_TCAUCH–S
AC_REX_TCAUCH–L
UC_RAM_UNIF
UC_RAM_SN
UC_RAM_TCAUCH–S
UC_RAM_TCAUCH–L
UC_REX_UNIF
UC_REX_SN
UC_REX_TCAUCH–S
UC_REX_TCAUCH–L
1000
750
500
t_n196
AC_RAM_UNIF
AC_RAM_SN
AC_RAM_TCAUCH–S
AC_RAM_TCAUCH–L
AC_REX_UNIF
AC_REX_SN
AC_REX_TCAUCH–S
AC_REX_TCAUCH–L
UC_RAM_UNIF
UC_RAM_SN
UC_RAM_TCAUCH–S
UC_RAM_TCAUCH–L
UC_REX_UNIF
UC_REX_SN
UC_REX_TCAUCH–S
UC_REX_TCAUCH–L
2000
1500
1000
500
t_n200
AC_RAM_UNIF
AC_RAM_SN
AC_RAM_TCAUCH–S
AC_RAM_TCAUCH–L
AC_REX_UNIF
AC_REX_SN
AC_REX_TCAUCH–S
AC_REX_TCAUCH–L
UC_RAM_UNIF
UC_RAM_SN
UC_RAM_TCAUCH–S
UC_RAM_TCAUCH–L
UC_REX_UNIF
UC_REX_SN
UC_REX_TCAUCH–S
UC_REX_TCAUCH–L
700
500
300

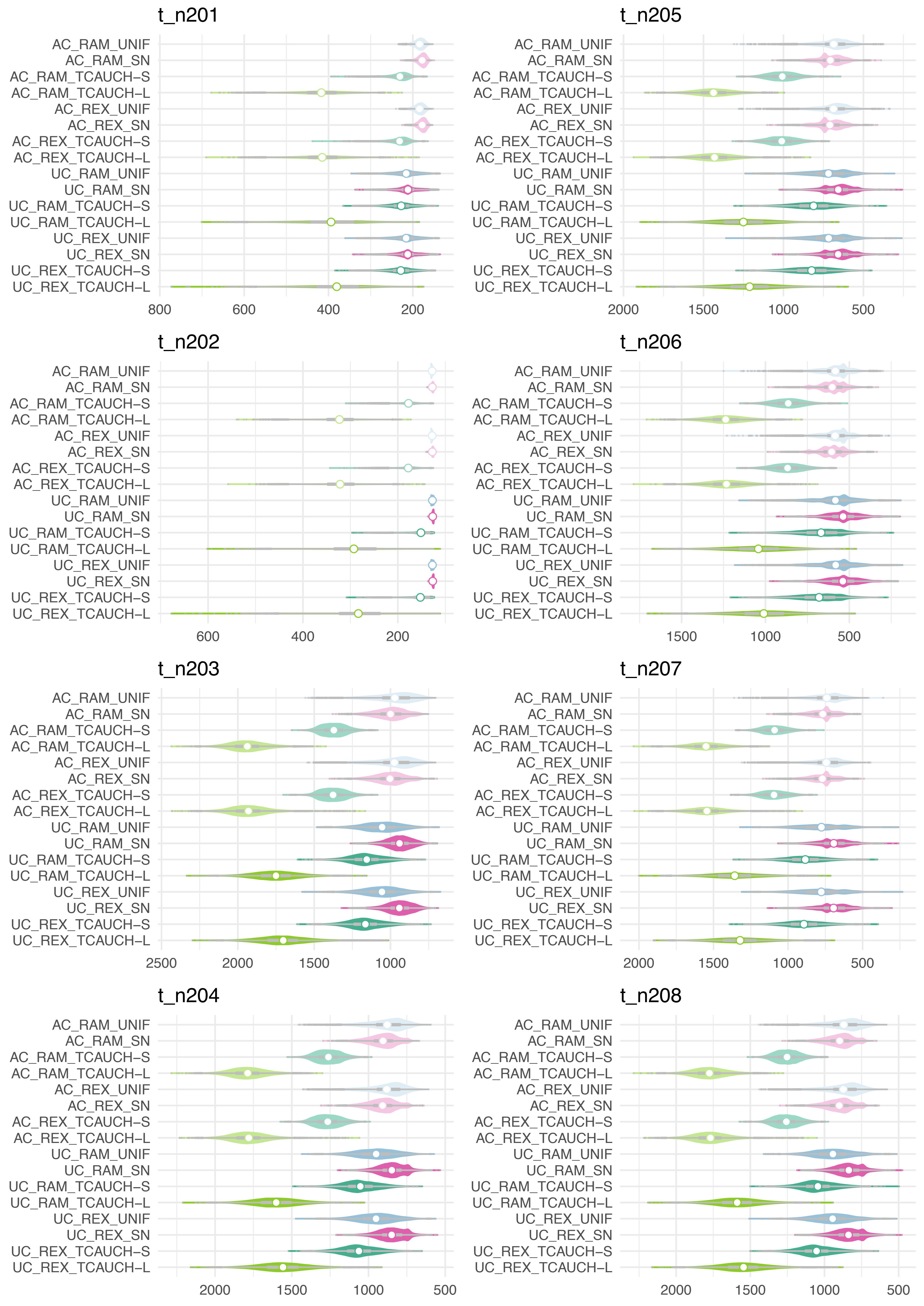
t_n201
AC_RAM_UNIF
AC_RAM_SN
AC_RAM_TCAUCH−S
AC_RAM_TCAUCH−L
AC_REX_UNIF
AC_REX_SN
AC_REX_TCAUCH−S
AC_REX_TCAUCH−L
UC_RAM_UNIF
UC_RAM_SN
UC_RAM_TCAUCH−S
UC_RAM_TCAUCH−L
UC_REX_UNIF
UC_REX_SN
UC_REX_TCAUCH−S
UC_REX_TCAUCH−L
800
600
400
200
t_n205
AC_RAM_UNIF
AC_RAM_SN
AC_RAM_TCAUCH−S
AC_RAM_TCAUCH−L
AC_REX_UNIF
AC_REX_SN
AC_REX_TCAUCH−S
AC_REX_TCAUCH−L
UC_RAM_UNIF
UC_RAM_SN
UC_RAM_TCAUCH−S
UC_RAM_TCAUCH−L
UC_REX_UNIF
UC_REX_SN
UC_REX_TCAUCH−S
UC_REX_TCAUCH−L
2000
1500
1000
500
t_n202
AC_RAM_UNIF
AC_RAM_SN
AC_RAM_TCAUCH−S
AC_RAM_TCAUCH−L
AC_REX_UNIF
AC_REX_SN
AC_REX_TCAUCH−S
AC_REX_TCAUCH−L
UC_RAM_UNIF
UC_RAM_SN
UC_RAM_TCAUCH−S
UC_RAM_TCAUCH−L
UC_REX_UNIF
UC_REX_SN
UC_REX_TCAUCH−S
UC_REX_TCAUCH−L
600
400
200
t_n206
AC_RAM_UNIF
AC_RAM_SN
AC_RAM_TCAUCH−S
AC_RAM_TCAUCH−L
AC_REX_UNIF
AC_REX_SN
AC_REX_TCAUCH−S
AC_REX_TCAUCH−L
UC_RAM_UNIF
UC_RAM_SN
UC_RAM_TCAUCH−S
UC_RAM_TCAUCH−L
UC_REX_UNIF
UC_REX_SN
UC_REX_TCAUCH−S
UC_REX_TCAUCH−L
1500
1000
500
t_n203
AC_RAM_UNIF
AC_RAM_SN
AC_RAM_TCAUCH−S
AC_RAM_TCAUCH−L
AC_REX_UNIF
AC_REX_SN
AC_REX_TCAUCH−S
AC_REX_TCAUCH−L
UC_RAM_UNIF
UC_RAM_SN
UC_RAM_TCAUCH−S
UC_RAM_TCAUCH−L
UC_REX_UNIF
UC_REX_SN
UC_REX_TCAUCH−S
UC_REX_TCAUCH−L
2500
2000
1500
1000
t_n207
AC_RAM_UNIF
AC_RAM_SN
AC_RAM_TCAUCH−S
AC_RAM_TCAUCH−L
AC_REX_UNIF
AC_REX_SN
AC_REX_TCAUCH−S
AC_REX_TCAUCH−L
UC_RAM_UNIF
UC_RAM_SN
UC_RAM_TCAUCH−S
UC_RAM_TCAUCH−L
UC_REX_UNIF
UC_REX_SN
UC_REX_TCAUCH−S
UC_REX_TCAUCH−L
2000
1500
1000
500
t_n204
AC_RAM_UNIF
AC_RAM_SN
AC_RAM_TCAUCH−S
AC_RAM_TCAUCH−L
AC_REX_UNIF
AC_REX_SN
AC_REX_TCAUCH−S
AC_REX_TCAUCH−L
UC_RAM_UNIF
UC_RAM_SN
UC_RAM_TCAUCH−S
UC_RAM_TCAUCH−L
UC_REX_UNIF
UC_REX_SN
UC_REX_TCAUCH−S
UC_REX_TCAUCH−L
2000
1500
1000
500
t_n208
AC_RAM_UNIF
AC_RAM_SN
AC_RAM_TCAUCH−S
AC_RAM_TCAUCH−L
AC_REX_UNIF
AC_REX_SN
AC_REX_TCAUCH−S
AC_REX_TCAUCH−L
UC_RAM_UNIF
UC_RAM_SN
UC_RAM_TCAUCH−S
UC_RAM_TCAUCH−L
UC_REX_UNIF
UC_REX_SN
UC_REX_TCAUCH−S
UC_REX_TCAUCH−L
2000
1500
1000
500

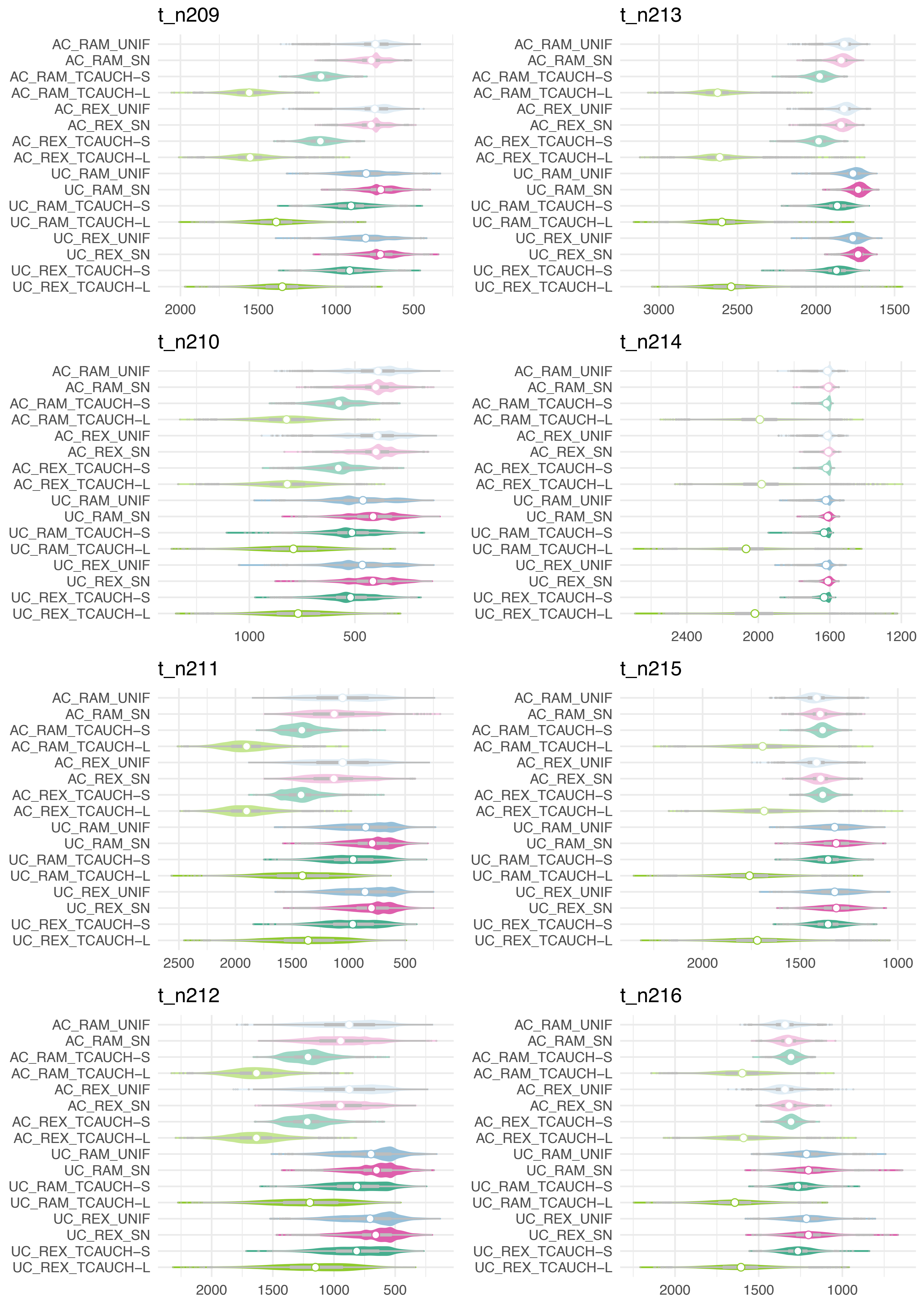

t_n209
AC_RAM_UNIF
AC_RAM_SN
AC_RAM_TCAUCH–S
AC_RAM_TCAUCH–L
AC_REX_UNIF
AC_REX_SN
AC_REX_TCAUCH–S
AC_REX_TCAUCH–L
UC_RAM_UNIF
UC_RAM_SN
UC_RAM_TCAUCH–S
UC_RAM_TCAUCH–L
UC_REX_UNIF
UC_REX_SN
UC_REX_TCAUCH–S
UC_REX_TCAUCH–L
2000
1500
1000
500
t_n213
3000
2500
2000
1500
t_n210
1000
500
t_n214
2400
2000
1600
1200
t_n211
2500
2000
1500
1000
500
t_n215
2000
1500
1000
t_n212
2000
1500
1000
500
t_n216
2000
1500
1000

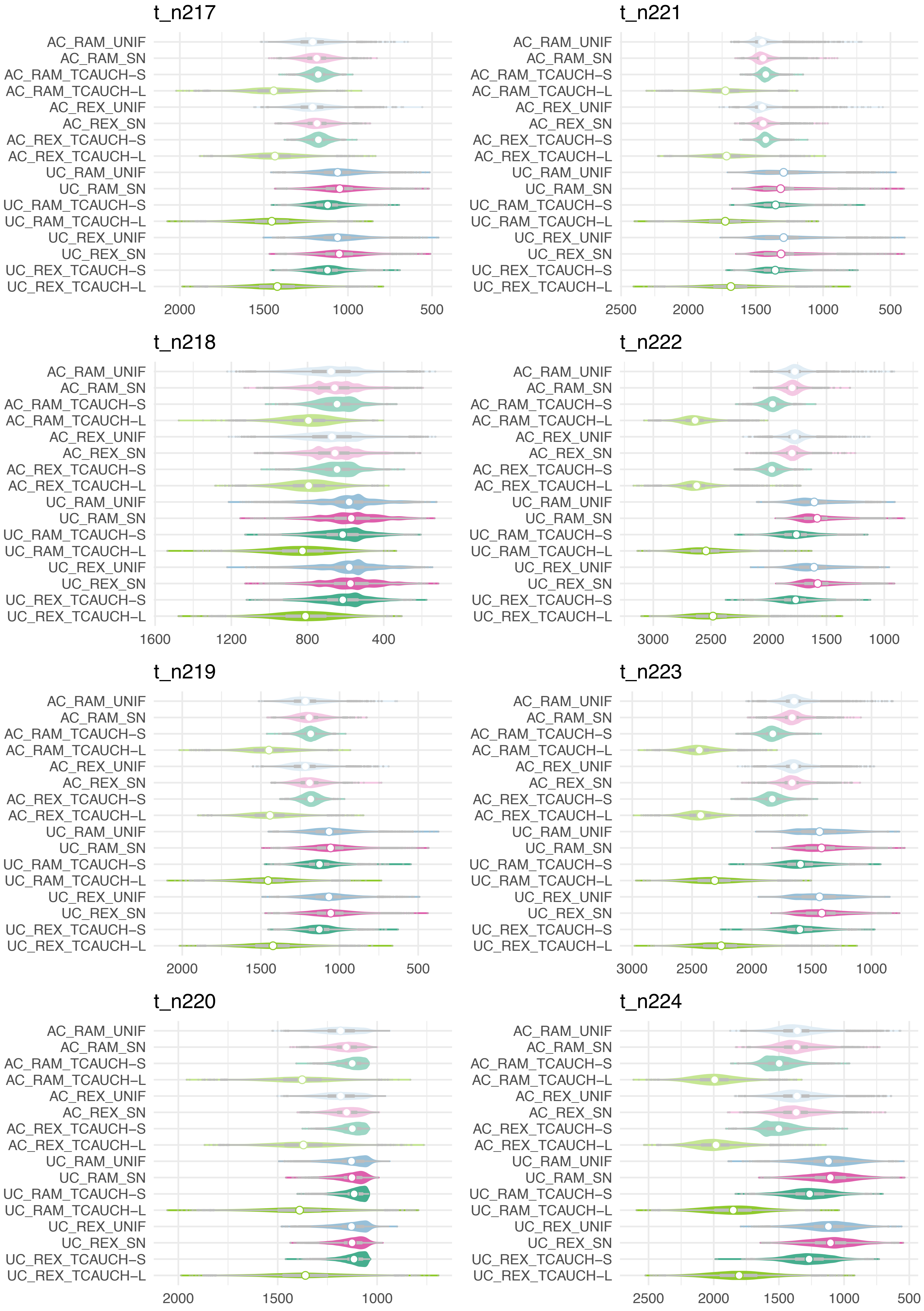

t_n217
AC_RAM_UNIF
AC_RAM_SN
AC_RAM_TCAUCH−S
AC_RAM_TCAUCH−L
AC_REX_UNIF
AC_REX_SN
AC_REX_TCAUCH−S
AC_REX_TCAUCH−L
UC_RAM_UNIF
UC_RAM_SN
UC_RAM_TCAUCH−S
UC_RAM_TCAUCH−L
UC_REX_UNIF
UC_REX_SN
UC_REX_TCAUCH−S
UC_REX_TCAUCH−L
2000 1500 1000 500
t_n221
2500 2000 1500 1000 500
t_n218
1600 1200 800 400
t_n222
3000 2500 2000 1500 1000
t_n219
2000 1500 1000 500
t_n223
3000 2500 2000 1500 1000
t_n220
2000 1500 1000
t_n224
2500 2000 1500 1000 500

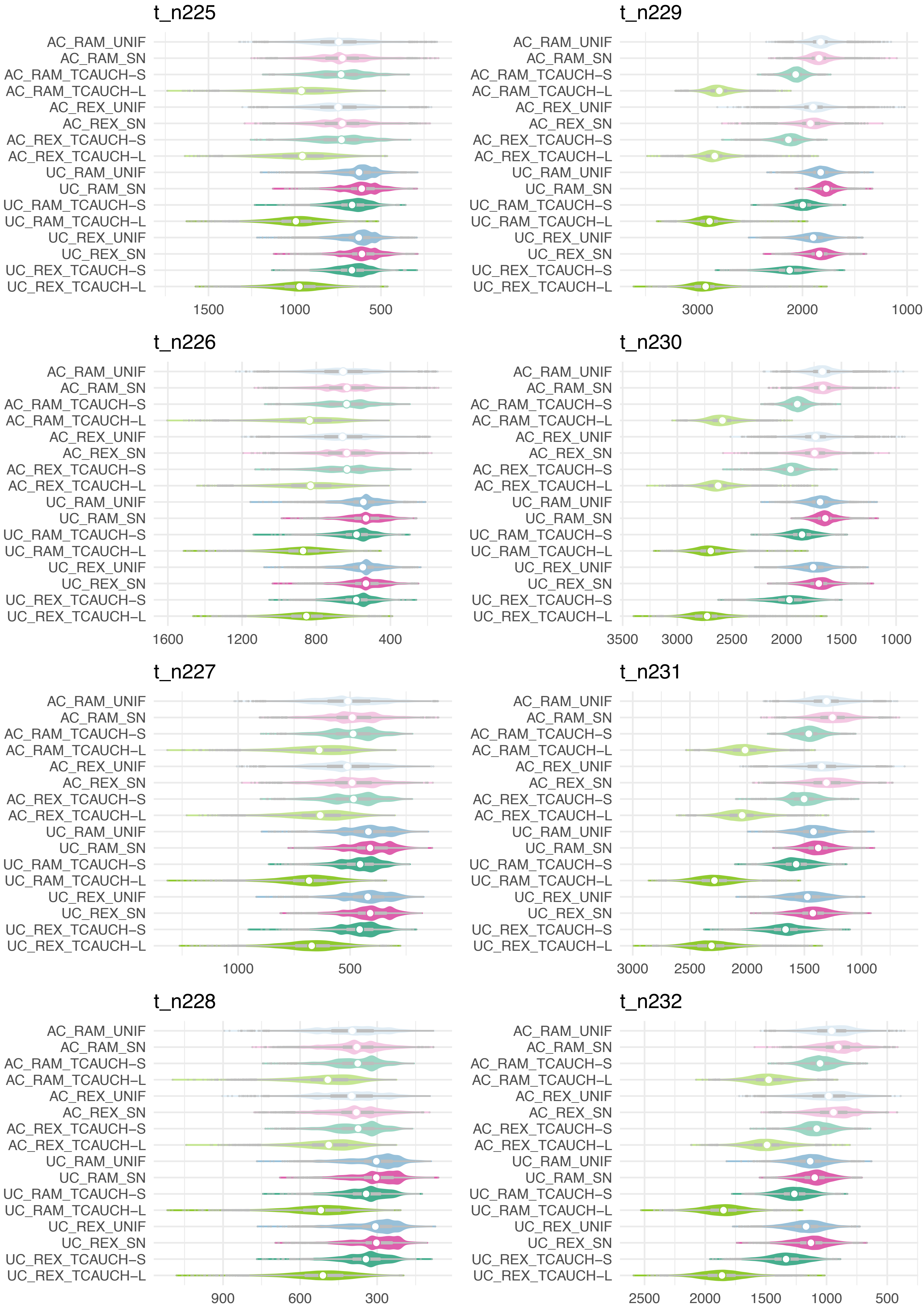
t_n225
AC_RAM_UNIF
AC_RAM_SN
AC_RAM_TCAUCH-S
AC_RAM_TCAUCH-L
AC_REX_UNIF
AC_REX_SN
AC_REX_TCAUCH-S
AC_REX_TCAUCH-L
UC_RAM_UNIF
UC_RAM_SN
UC_RAM_TCAUCH-S
UC_RAM_TCAUCH-L
UC_REX_UNIF
UC_REX_SN
UC_REX_TCAUCH-S
UC_REX_TCAUCH-L
1500
1000
500
t_n229
AC_RAM_UNIF
AC_RAM_SN
AC_RAM_TCAUCH-S
AC_RAM_TCAUCH-L
AC_REX_UNIF
AC_REX_SN
AC_REX_TCAUCH-S
AC_REX_TCAUCH-L
UC_RAM_UNIF
UC_RAM_SN
UC_RAM_TCAUCH-S
UC_RAM_TCAUCH-L
UC_REX_UNIF
UC_REX_SN
UC_REX_TCAUCH-S
UC_REX_TCAUCH-L
3000
2000
1000
t_n226
AC_RAM_UNIF
AC_RAM_SN
AC_RAM_TCAUCH-S
AC_RAM_TCAUCH-L
AC_REX_UNIF
AC_REX_SN
AC_REX_TCAUCH-S
AC_REX_TCAUCH-L
UC_RAM_UNIF
UC_RAM_SN
UC_RAM_TCAUCH-S
UC_RAM_TCAUCH-L
UC_REX_UNIF
UC_REX_SN
UC_REX_TCAUCH-S
UC_REX_TCAUCH-L
1600
1200
800
400
t_n230
AC_RAM_UNIF
AC_RAM_SN
AC_RAM_TCAUCH-S
AC_RAM_TCAUCH-L
AC_REX_UNIF
AC_REX_SN
AC_REX_TCAUCH-S
AC_REX_TCAUCH-L
UC_RAM_UNIF
UC_RAM_SN
UC_RAM_TCAUCH-S
UC_RAM_TCAUCH-L
UC_REX_UNIF
UC_REX_SN
UC_REX_TCAUCH-S
UC_REX_TCAUCH-L
3500
3000
2500
2000
1500
1000
t_n227
AC_RAM_UNIF
AC_RAM_SN
AC_RAM_TCAUCH-S
AC_RAM_TCAUCH-L
AC_REX_UNIF
AC_REX_SN
AC_REX_TCAUCH-S
AC_REX_TCAUCH-L
UC_RAM_UNIF
UC_RAM_SN
UC_RAM_TCAUCH-S
UC_RAM_TCAUCH-L
UC_REX_UNIF
UC_REX_SN
UC_REX_TCAUCH-S
UC_REX_TCAUCH-L
1000
500
t_n231
AC_RAM_UNIF
AC_RAM_SN
AC_RAM_TCAUCH-S
AC_RAM_TCAUCH-L
AC_REX_UNIF
AC_REX_SN
AC_REX_TCAUCH-S
AC_REX_TCAUCH-L
UC_RAM_UNIF
UC_RAM_SN
UC_RAM_TCAUCH-S
UC_RAM_TCAUCH-L
UC_REX_UNIF
UC_REX_SN
UC_REX_TCAUCH-S
UC_REX_TCAUCH-L
3000
2500
2000
1500
1000
t_n228
AC_RAM_UNIF
AC_RAM_SN
AC_RAM_TCAUCH-S
AC_RAM_TCAUCH-L
AC_REX_UNIF
AC_REX_SN
AC_REX_TCAUCH-S
AC_REX_TCAUCH-L
UC_RAM_UNIF
UC_RAM_SN
UC_RAM_TCAUCH-S
UC_RAM_TCAUCH-L
UC_REX_UNIF
UC_REX_SN
UC_REX_TCAUCH-S
UC_REX_TCAUCH-L
900
600
300
t_n232
AC_RAM_UNIF
AC_RAM_SN
AC_RAM_TCAUCH-S
AC_RAM_TCAUCH-L
AC_REX_UNIF
AC_REX_SN
AC_REX_TCAUCH-S
AC_REX_TCAUCH-L
UC_RAM_UNIF
UC_RAM_SN
UC_RAM_TCAUCH-S
UC_RAM_TCAUCH-L
UC_REX_UNIF
UC_REX_SN
UC_REX_TCAUCH-S
UC_REX_TCAUCH-L
2500
2000
1500
1000
500

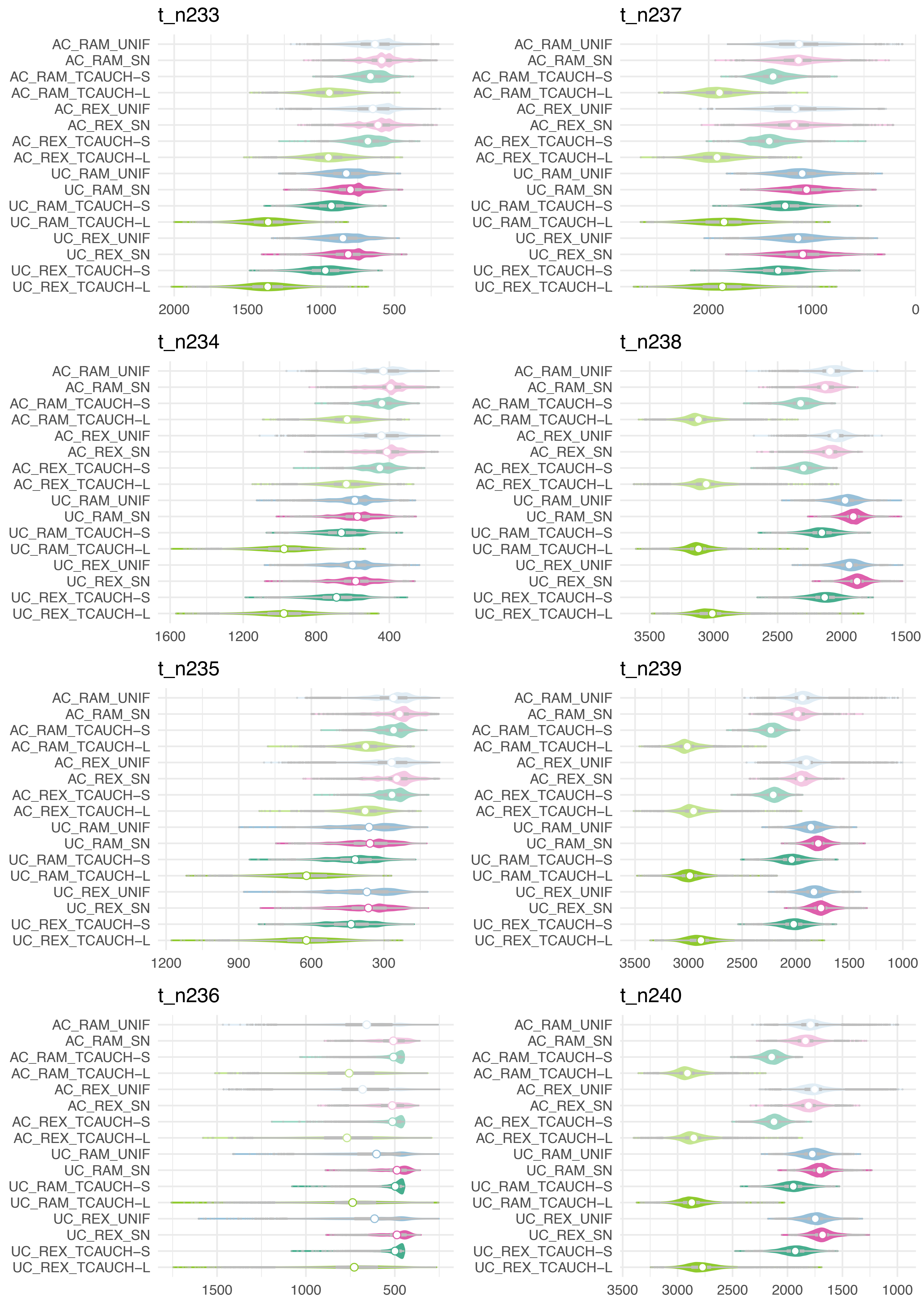
t_n233
AC_RAM_UNIF
AC_RAM_SN
AC_RAM_TCAUCH–S
AC_RAM_TCAUCH–L
AC_REX_UNIF
AC_REX_SN
AC_REX_TCAUCH–S
AC_REX_TCAUCH–L
UC_RAM_UNIF
UC_RAM_SN
UC_RAM_TCAUCH–S
UC_RAM_TCAUCH–L
UC_REX_UNIF
UC_REX_SN
UC_REX_TCAUCH–S
UC_REX_TCAUCH–L
2000
1500
1000
500
t_n237
AC_RAM_UNIF
AC_RAM_SN
AC_RAM_TCAUCH–S
AC_RAM_TCAUCH–L
AC_REX_UNIF
AC_REX_SN
AC_REX_TCAUCH–S
AC_REX_TCAUCH–L
UC_RAM_UNIF
UC_RAM_SN
UC_RAM_TCAUCH–S
UC_RAM_TCAUCH–L
UC_REX_UNIF
UC_REX_SN
UC_REX_TCAUCH–S
UC_REX_TCAUCH–L
2000
1000
0
t_n234
AC_RAM_UNIF
AC_RAM_SN
AC_RAM_TCAUCH–S
AC_RAM_TCAUCH–L
AC_REX_UNIF
AC_REX_SN
AC_REX_TCAUCH–S
AC_REX_TCAUCH–L
UC_RAM_UNIF
UC_RAM_SN
UC_RAM_TCAUCH–S
UC_RAM_TCAUCH–L
UC_REX_UNIF
UC_REX_SN
UC_REX_TCAUCH–S
UC_REX_TCAUCH–L
1600
1200
800
400
t_n238
AC_RAM_UNIF
AC_RAM_SN
AC_RAM_TCAUCH–S
AC_RAM_TCAUCH–L
AC_REX_UNIF
AC_REX_SN
AC_REX_TCAUCH–S
AC_REX_TCAUCH–L
UC_RAM_UNIF
UC_RAM_SN
UC_RAM_TCAUCH–S
UC_RAM_TCAUCH–L
UC_REX_UNIF
UC_REX_SN
UC_REX_TCAUCH–S
UC_REX_TCAUCH–L
3500
3000
2500
2000
1500
t_n235
AC_RAM_UNIF
AC_RAM_SN
AC_RAM_TCAUCH–S
AC_RAM_TCAUCH–L
AC_REX_UNIF
AC_REX_SN
AC_REX_TCAUCH–S
AC_REX_TCAUCH–L
UC_RAM_UNIF
UC_RAM_SN
UC_RAM_TCAUCH–S
UC_RAM_TCAUCH–L
UC_REX_UNIF
UC_REX_SN
UC_REX_TCAUCH–S
UC_REX_TCAUCH–L
1200
900
600
300
t_n239
AC_RAM_UNIF
AC_RAM_SN
AC_RAM_TCAUCH–S
AC_RAM_TCAUCH–L
AC_REX_UNIF
AC_REX_SN
AC_REX_TCAUCH–S
AC_REX_TCAUCH–L
UC_RAM_UNIF
UC_RAM_SN
UC_RAM_TCAUCH–S
UC_RAM_TCAUCH–L
UC_REX_UNIF
UC_REX_SN
UC_REX_TCAUCH–S
UC_REX_TCAUCH–L
3500
3000
2500
2000
1500
1000
t_n236
AC_RAM_UNIF
AC_RAM_SN
AC_RAM_TCAUCH–S
AC_RAM_TCAUCH–L
AC_REX_UNIF
AC_REX_SN
AC_REX_TCAUCH–S
AC_REX_TCAUCH–L
UC_RAM_UNIF
UC_RAM_SN
UC_RAM_TCAUCH–S
UC_RAM_TCAUCH–L
UC_REX_UNIF
UC_REX_SN
UC_REX_TCAUCH–S
UC_REX_TCAUCH–L
1500
1000
500
t_n240
AC_RAM_UNIF
AC_RAM_SN
AC_RAM_TCAUCH–S
AC_RAM_TCAUCH–L
AC_REX_UNIF
AC_REX_SN
AC_REX_TCAUCH–S
AC_REX_TCAUCH–L
UC_RAM_UNIF
UC_RAM_SN
UC_RAM_TCAUCH–S
UC_RAM_TCAUCH–L
UC_REX_UNIF
UC_REX_SN
UC_REX_TCAUCH–S
UC_REX_TCAUCH–L
3500
3000
2500
2000
1500
1000

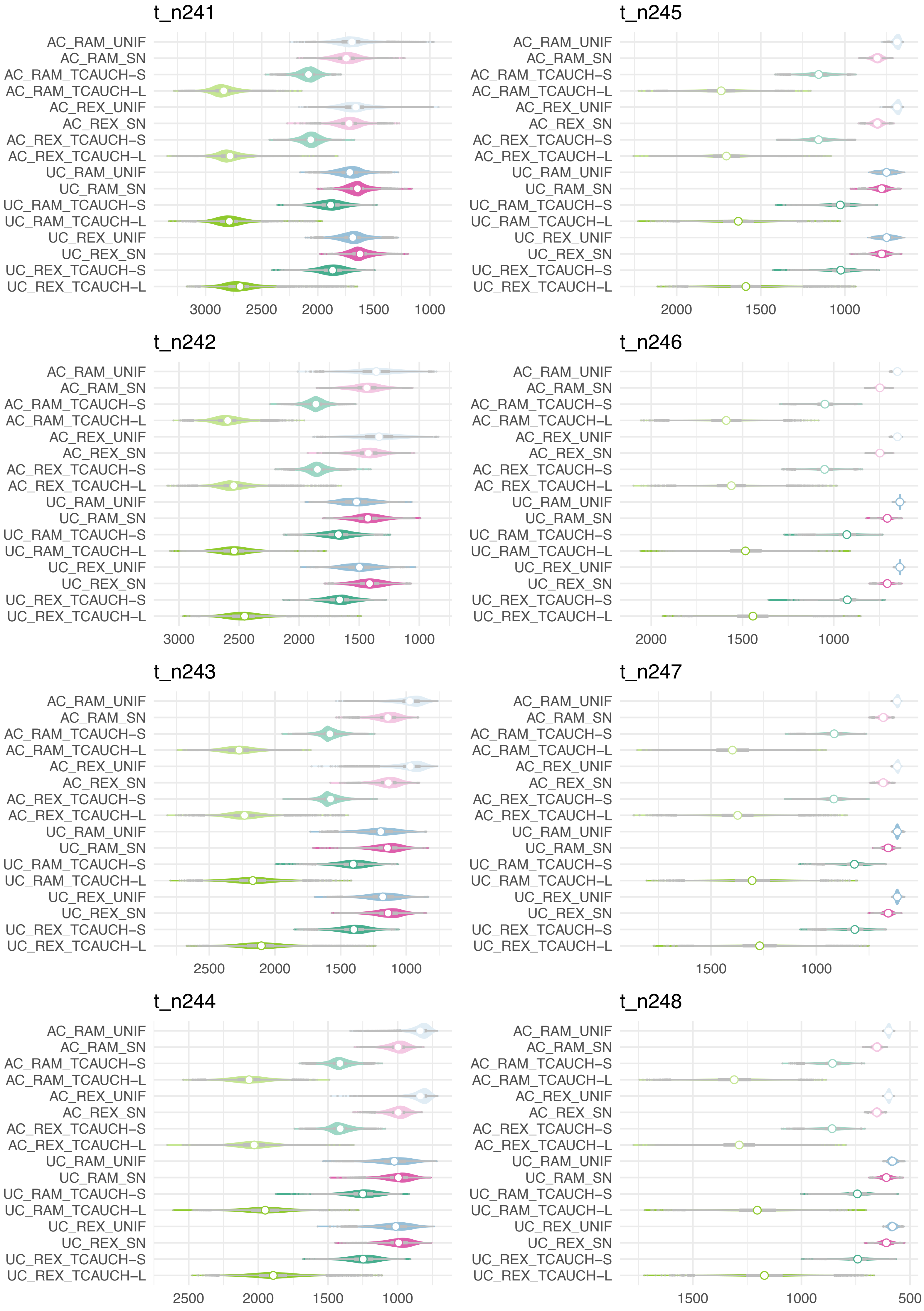
t_n241
AC_RAM_UNIF
AC_RAM_SN
AC_RAM_TCAUCH–S
AC_RAM_TCAUCH–L
AC_REX_UNIF
AC_REX_SN
AC_REX_TCAUCH–S
AC_REX_TCAUCH–L
UC_RAM_UNIF
UC_RAM_SN
UC_RAM_TCAUCH–S
UC_RAM_TCAUCH–L
UC_REX_UNIF
UC_REX_SN
UC_REX_TCAUCH–S
UC_REX_TCAUCH–L
3000
2500
2000
1500
1000
t_n245
AC_RAM_UNIF
AC_RAM_SN
AC_RAM_TCAUCH–S
AC_RAM_TCAUCH–L
AC_REX_UNIF
AC_REX_SN
AC_REX_TCAUCH–S
AC_REX_TCAUCH–L
UC_RAM_UNIF
UC_RAM_SN
UC_RAM_TCAUCH–S
UC_RAM_TCAUCH–L
UC_REX_UNIF
UC_REX_SN
UC_REX_TCAUCH–S
UC_REX_TCAUCH–L
2000
1500
1000
t_n242
AC_RAM_UNIF
AC_RAM_SN
AC_RAM_TCAUCH–S
AC_RAM_TCAUCH–L
AC_REX_UNIF
AC_REX_SN
AC_REX_TCAUCH–S
AC_REX_TCAUCH–L
UC_RAM_UNIF
UC_RAM_SN
UC_RAM_TCAUCH–S
UC_RAM_TCAUCH–L
UC_REX_UNIF
UC_REX_SN
UC_REX_TCAUCH–S
UC_REX_TCAUCH–L
3000
2500
2000
1500
1000
t_n246
AC_RAM_UNIF
AC_RAM_SN
AC_RAM_TCAUCH–S
AC_RAM_TCAUCH–L
AC_REX_UNIF
AC_REX_SN
AC_REX_TCAUCH–S
AC_REX_TCAUCH–L
UC_RAM_UNIF
UC_RAM_SN
UC_RAM_TCAUCH–S
UC_RAM_TCAUCH–L
UC_REX_UNIF
UC_REX_SN
UC_REX_TCAUCH–S
UC_REX_TCAUCH–L
2000
1500
1000
t_n243
AC_RAM_UNIF
AC_RAM_SN
AC_RAM_TCAUCH–S
AC_RAM_TCAUCH–L
AC_REX_UNIF
AC_REX_SN
AC_REX_TCAUCH–S
AC_REX_TCAUCH–L
UC_RAM_UNIF
UC_RAM_SN
UC_RAM_TCAUCH–S
UC_RAM_TCAUCH–L
UC_REX_UNIF
UC_REX_SN
UC_REX_TCAUCH–S
UC_REX_TCAUCH–L
2500
2000
1500
1000
t_n247
AC_RAM_UNIF
AC_RAM_SN
AC_RAM_TCAUCH–S
AC_RAM_TCAUCH–L
AC_REX_UNIF
AC_REX_SN
AC_REX_TCAUCH–S
AC_REX_TCAUCH–L
UC_RAM_UNIF
UC_RAM_SN
UC_RAM_TCAUCH–S
UC_RAM_TCAUCH–L
UC_REX_UNIF
UC_REX_SN
UC_REX_TCAUCH–S
UC_REX_TCAUCH–L
1500
1000
t_n244
AC_RAM_UNIF
AC_RAM_SN
AC_RAM_TCAUCH–S
AC_RAM_TCAUCH–L
AC_REX_UNIF
AC_REX_SN
AC_REX_TCAUCH–S
AC_REX_TCAUCH–L
UC_RAM_UNIF
UC_RAM_SN
UC_RAM_TCAUCH–S
UC_RAM_TCAUCH–L
UC_REX_UNIF
UC_REX_SN
UC_REX_TCAUCH–S
UC_REX_TCAUCH–L
2500
2000
1500
1000
t_n248
AC_RAM_UNIF
AC_RAM_SN
AC_RAM_TCAUCH–S
AC_RAM_TCAUCH–L
AC_REX_UNIF
AC_REX_SN
AC_REX_TCAUCH–S
AC_REX_TCAUCH–L
UC_RAM_UNIF
UC_RAM_SN
UC_RAM_TCAUCH–S
UC_RAM_TCAUCH–L
UC_REX_UNIF
UC_REX_SN
UC_REX_TCAUCH–S
UC_REX_TCAUCH–L
1500
1000
500

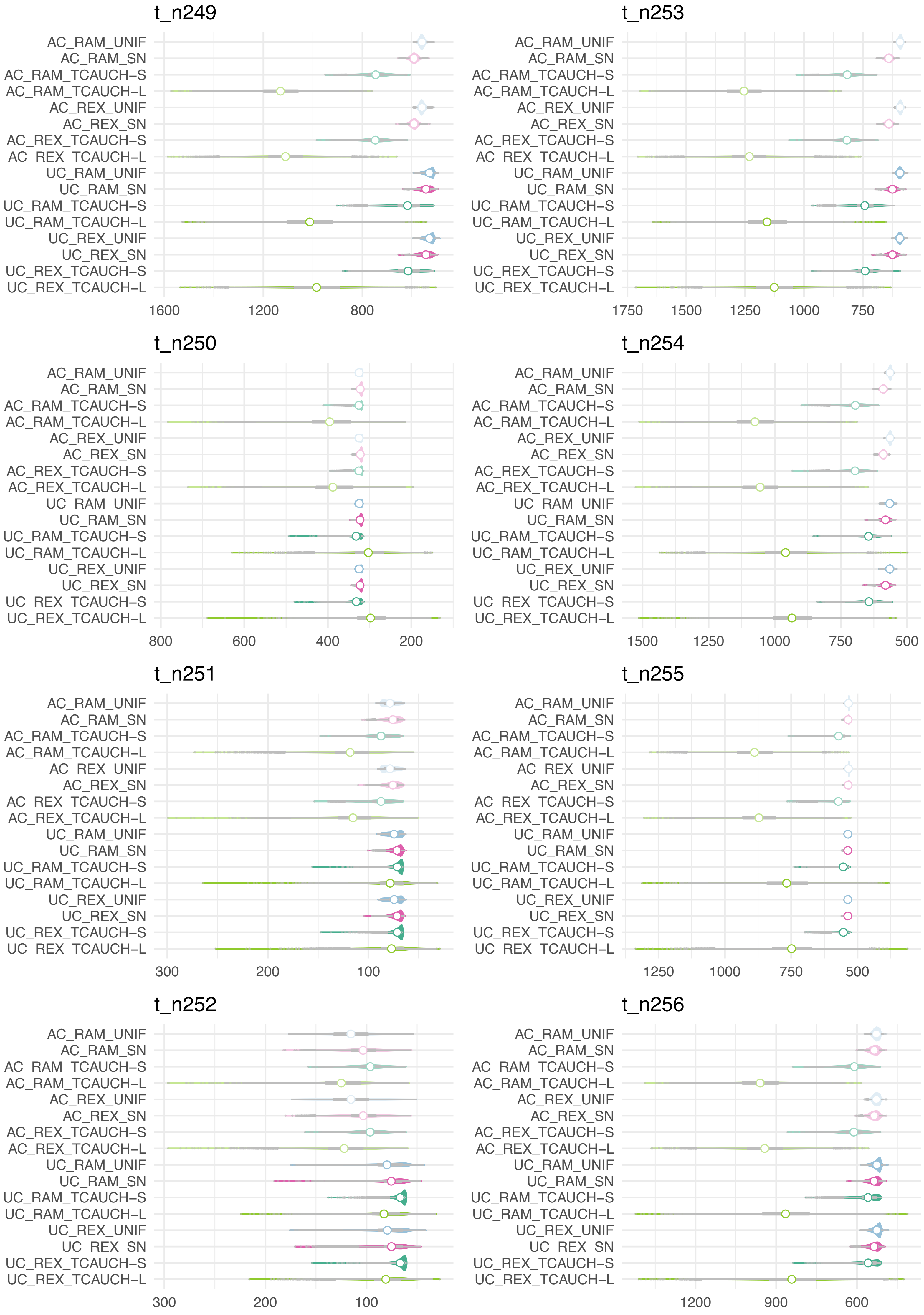

t_n249
AC_RAM_UNIF
AC_RAM_SN
AC_RAM_TCAUCH−S
AC_RAM_TCAUCH−L
AC_REX_UNIF
AC_REX_SN
AC_REX_TCAUCH−S
AC_REX_TCAUCH−L
UC_RAM_UNIF
UC_RAM_SN
UC_RAM_TCAUCH−S
UC_RAM_TCAUCH−L
UC_REX_UNIF
UC_REX_SN
UC_REX_TCAUCH−S
UC_REX_TCAUCH−L
1600
1200
800
t_n253
1750
1500
1250
1000
750
t_n250
800
600
400
200
t_n254
1500
1250
1000
750
500
t_n251
300
200
100
t_n255
1250
1000
750
500
t_n252
300
200
100
t_n256
1200
900
600

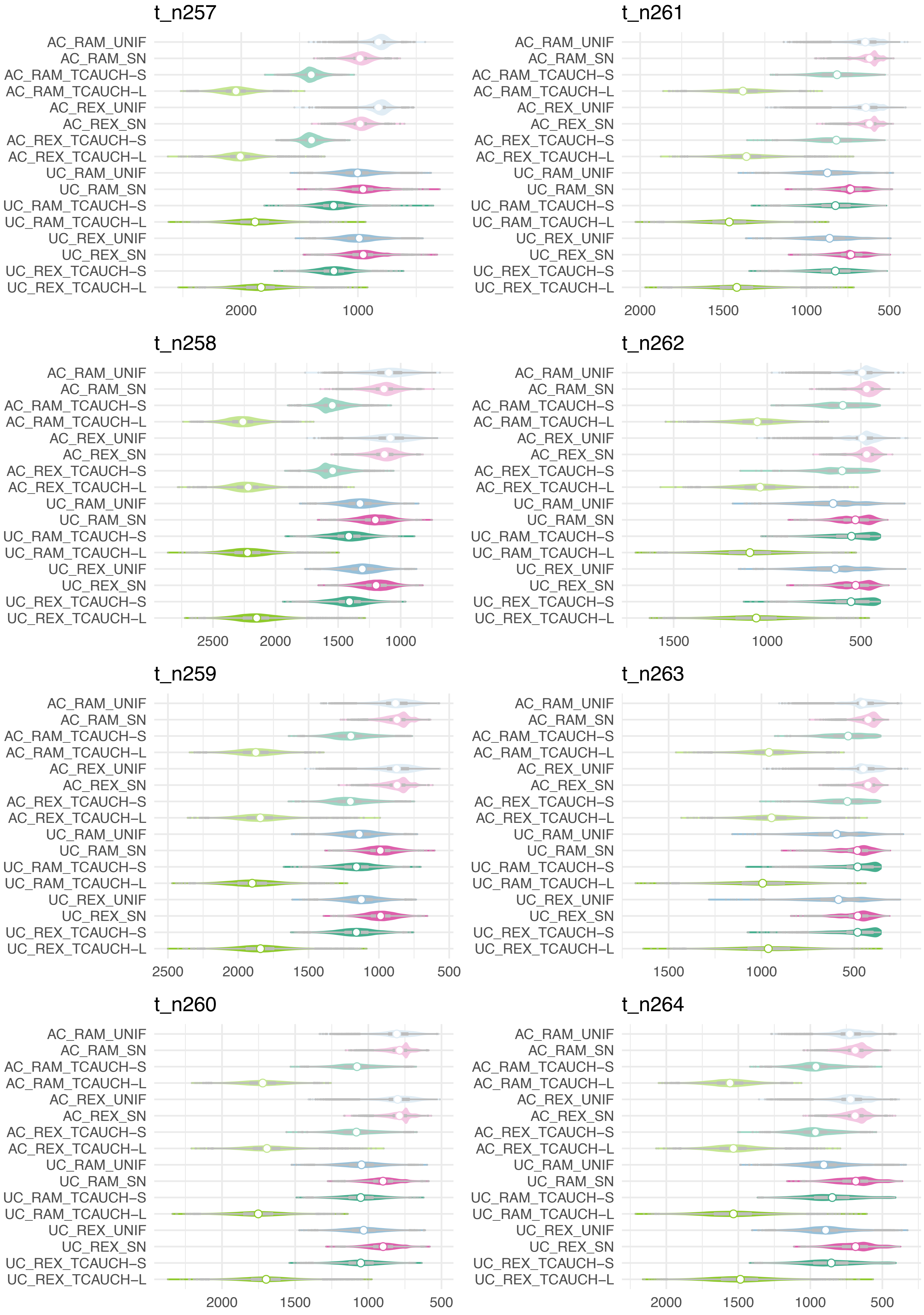

t_n257
AC_RAM_UNIF
AC_RAM_SN
AC_RAM_TCAUCH-S
AC_RAM_TCAUCH-L
AC_REX_UNIF
AC_REX_SN
AC_REX_TCAUCH-S
AC_REX_TCAUCH-L
UC_RAM_UNIF
UC_RAM_SN
UC_RAM_TCAUCH-S
UC_RAM_TCAUCH-L
UC_REX_UNIF
UC_REX_SN
UC_REX_TCAUCH-S
UC_REX_TCAUCH-L
2000
1000
t_n261
AC_RAM_UNIF
AC_RAM_SN
AC_RAM_TCAUCH-S
AC_RAM_TCAUCH-L
AC_REX_UNIF
AC_REX_SN
AC_REX_TCAUCH-S
AC_REX_TCAUCH-L
UC_RAM_UNIF
UC_RAM_SN
UC_RAM_TCAUCH-S
UC_RAM_TCAUCH-L
UC_REX_UNIF
UC_REX_SN
UC_REX_TCAUCH-S
UC_REX_TCAUCH-L
2000
1500
1000
500
t_n258
AC_RAM_UNIF
AC_RAM_SN
AC_RAM_TCAUCH-S
AC_RAM_TCAUCH-L
AC_REX_UNIF
AC_REX_SN
AC_REX_TCAUCH-S
AC_REX_TCAUCH-L
UC_RAM_UNIF
UC_RAM_SN
UC_RAM_TCAUCH-S
UC_RAM_TCAUCH-L
UC_REX_UNIF
UC_REX_SN
UC_REX_TCAUCH-S
UC_REX_TCAUCH-L
2500
2000
1500
1000
t_n262
AC_RAM_UNIF
AC_RAM_SN
AC_RAM_TCAUCH-S
AC_RAM_TCAUCH-L
AC_REX_UNIF
AC_REX_SN
AC_REX_TCAUCH-S
AC_REX_TCAUCH-L
UC_RAM_UNIF
UC_RAM_SN
UC_RAM_TCAUCH-S
UC_RAM_TCAUCH-L
UC_REX_UNIF
UC_REX_SN
UC_REX_TCAUCH-S
UC_REX_TCAUCH-L
1500
1000
500
t_n259
AC_RAM_UNIF
AC_RAM_SN
AC_RAM_TCAUCH-S
AC_RAM_TCAUCH-L
AC_REX_UNIF
AC_REX_SN
AC_REX_TCAUCH-S
AC_REX_TCAUCH-L
UC_RAM_UNIF
UC_RAM_SN
UC_RAM_TCAUCH-S
UC_RAM_TCAUCH-L
UC_REX_UNIF
UC_REX_SN
UC_REX_TCAUCH-S
UC_REX_TCAUCH-L
2500
2000
1500
1000
500
t_n263
AC_RAM_UNIF
AC_RAM_SN
AC_RAM_TCAUCH-S
AC_RAM_TCAUCH-L
AC_REX_UNIF
AC_REX_SN
AC_REX_TCAUCH-S
AC_REX_TCAUCH-L
UC_RAM_UNIF
UC_RAM_SN
UC_RAM_TCAUCH-S
UC_RAM_TCAUCH-L
UC_REX_UNIF
UC_REX_SN
UC_REX_TCAUCH-S
UC_REX_TCAUCH-L
1500
1000
500
t_n260
AC_RAM_UNIF
AC_RAM_SN
AC_RAM_TCAUCH-S
AC_RAM_TCAUCH-L
AC_REX_UNIF
AC_REX_SN
AC_REX_TCAUCH-S
AC_REX_TCAUCH-L
UC_RAM_UNIF
UC_RAM_SN
UC_RAM_TCAUCH-S
UC_RAM_TCAUCH-L
UC_REX_UNIF
UC_REX_SN
UC_REX_TCAUCH-S
UC_REX_TCAUCH-L
2000
1500
1000
500
t_n264
AC_RAM_UNIF
AC_RAM_SN
AC_RAM_TCAUCH-S
AC_RAM_TCAUCH-L
AC_REX_UNIF
AC_REX_SN
AC_REX_TCAUCH-S
AC_REX_TCAUCH-L
UC_RAM_UNIF
UC_RAM_SN
UC_RAM_TCAUCH-S
UC_RAM_TCAUCH-L
UC_REX_UNIF
UC_REX_SN
UC_REX_TCAUCH-S
UC_REX_TCAUCH-L
2000
1500
1000
500

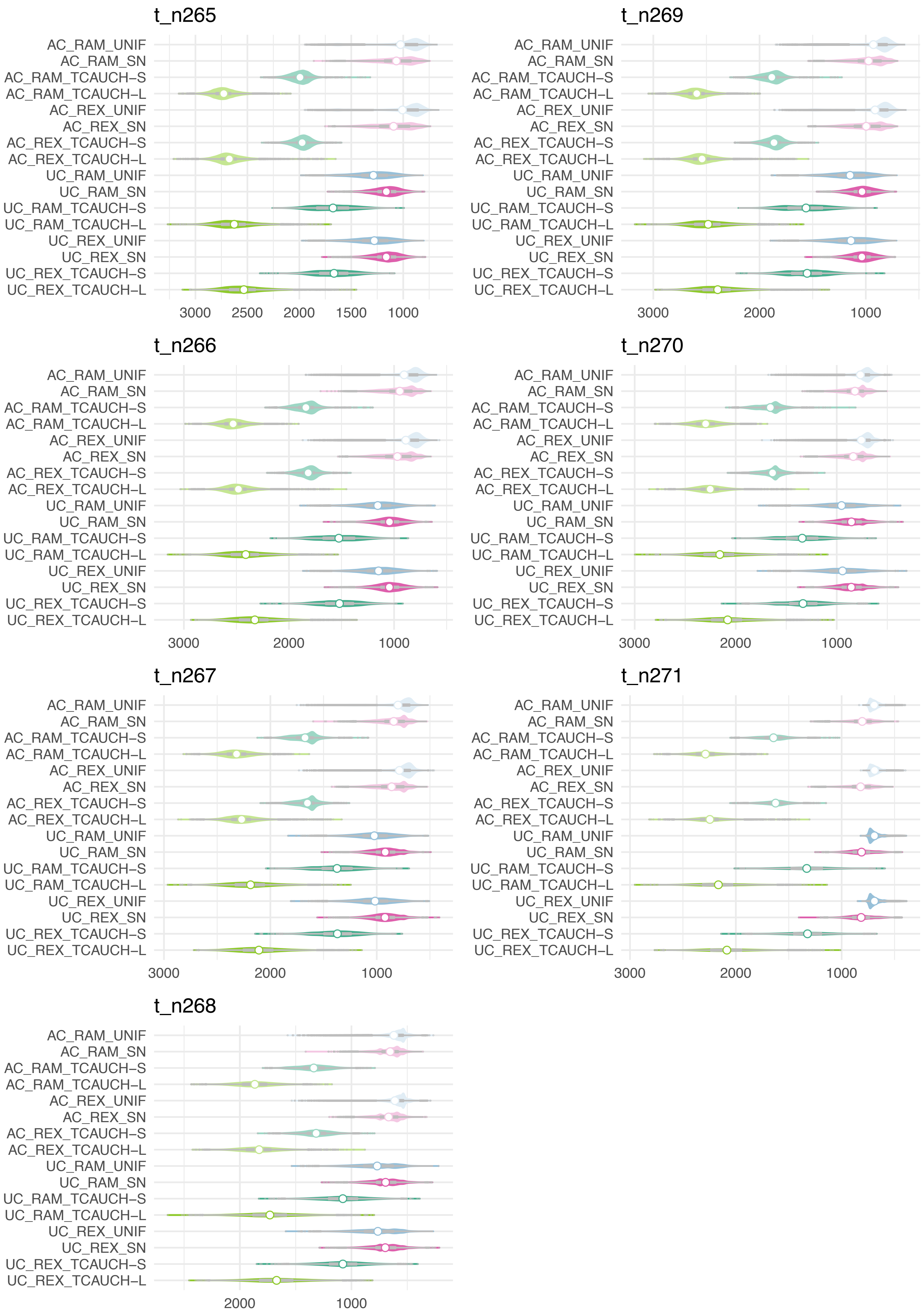
t_n265
AC_RAM_UNIF
AC_RAM_SN
AC_RAM_TCAUCH–S
AC_RAM_TCAUCH–L
AC_REX_UNIF
AC_REX_SN
AC_REX_TCAUCH–S
AC_REX_TCAUCH–L
UC_RAM_UNIF
UC_RAM_SN
UC_RAM_TCAUCH–S
UC_RAM_TCAUCH–L
UC_REX_UNIF
UC_REX_SN
UC_REX_TCAUCH–S
UC_REX_TCAUCH–L
3000
2500
2000
1500
1000
t_n269
AC_RAM_UNIF
AC_RAM_SN
AC_RAM_TCAUCH–S
AC_RAM_TCAUCH–L
AC_REX_UNIF
AC_REX_SN
AC_REX_TCAUCH–S
AC_REX_TCAUCH–L
UC_RAM_UNIF
UC_RAM_SN
UC_RAM_TCAUCH–S
UC_RAM_TCAUCH–L
UC_REX_UNIF
UC_REX_SN
UC_REX_TCAUCH–S
UC_REX_TCAUCH–L
3000
2000
1000
t_n266
AC_RAM_UNIF
AC_RAM_SN
AC_RAM_TCAUCH–S
AC_RAM_TCAUCH–L
AC_REX_UNIF
AC_REX_SN
AC_REX_TCAUCH–S
AC_REX_TCAUCH–L
UC_RAM_UNIF
UC_RAM_SN
UC_RAM_TCAUCH–S
UC_RAM_TCAUCH–L
UC_REX_UNIF
UC_REX_SN
UC_REX_TCAUCH–S
UC_REX_TCAUCH–L
3000
2000
1000
t_n270
AC_RAM_UNIF
AC_RAM_SN
AC_RAM_TCAUCH–S
AC_RAM_TCAUCH–L
AC_REX_UNIF
AC_REX_SN
AC_REX_TCAUCH–S
AC_REX_TCAUCH–L
UC_RAM_UNIF
UC_RAM_SN
UC_RAM_TCAUCH–S
UC_RAM_TCAUCH–L
UC_REX_UNIF
UC_REX_SN
UC_REX_TCAUCH–S
UC_REX_TCAUCH–L
3000
2000
1000
t_n267
AC_RAM_UNIF
AC_RAM_SN
AC_RAM_TCAUCH–S
AC_RAM_TCAUCH–L
AC_REX_UNIF
AC_REX_SN
AC_REX_TCAUCH–S
AC_REX_TCAUCH–L
UC_RAM_UNIF
UC_RAM_SN
UC_RAM_TCAUCH–S
UC_RAM_TCAUCH–L
UC_REX_UNIF
UC_REX_SN
UC_REX_TCAUCH–S
UC_REX_TCAUCH–L
3000
2000
1000
t_n271
AC_RAM_UNIF
AC_RAM_SN
AC_RAM_TCAUCH–S
AC_RAM_TCAUCH–L
AC_REX_UNIF
AC_REX_SN
AC_REX_TCAUCH–S
AC_REX_TCAUCH–L
UC_RAM_UNIF
UC_RAM_SN
UC_RAM_TCAUCH–S
UC_RAM_TCAUCH–L
UC_REX_UNIF
UC_REX_SN
UC_REX_TCAUCH–S
UC_REX_TCAUCH–L
3000
2000
1000
t_n268
AC_RAM_UNIF
AC_RAM_SN
AC_RAM_TCAUCH–S
AC_RAM_TCAUCH–L
AC_REX_UNIF
AC_REX_SN
AC_REX_TCAUCH–S
AC_REX_TCAUCH–L
UC_RAM_UNIF
UC_RAM_SN
UC_RAM_TCAUCH–S
UC_RAM_TCAUCH–L
UC_REX_UNIF
UC_REX_SN
UC_REX_TCAUCH–S
UC_REX_TCAUCH–L
2000
1000

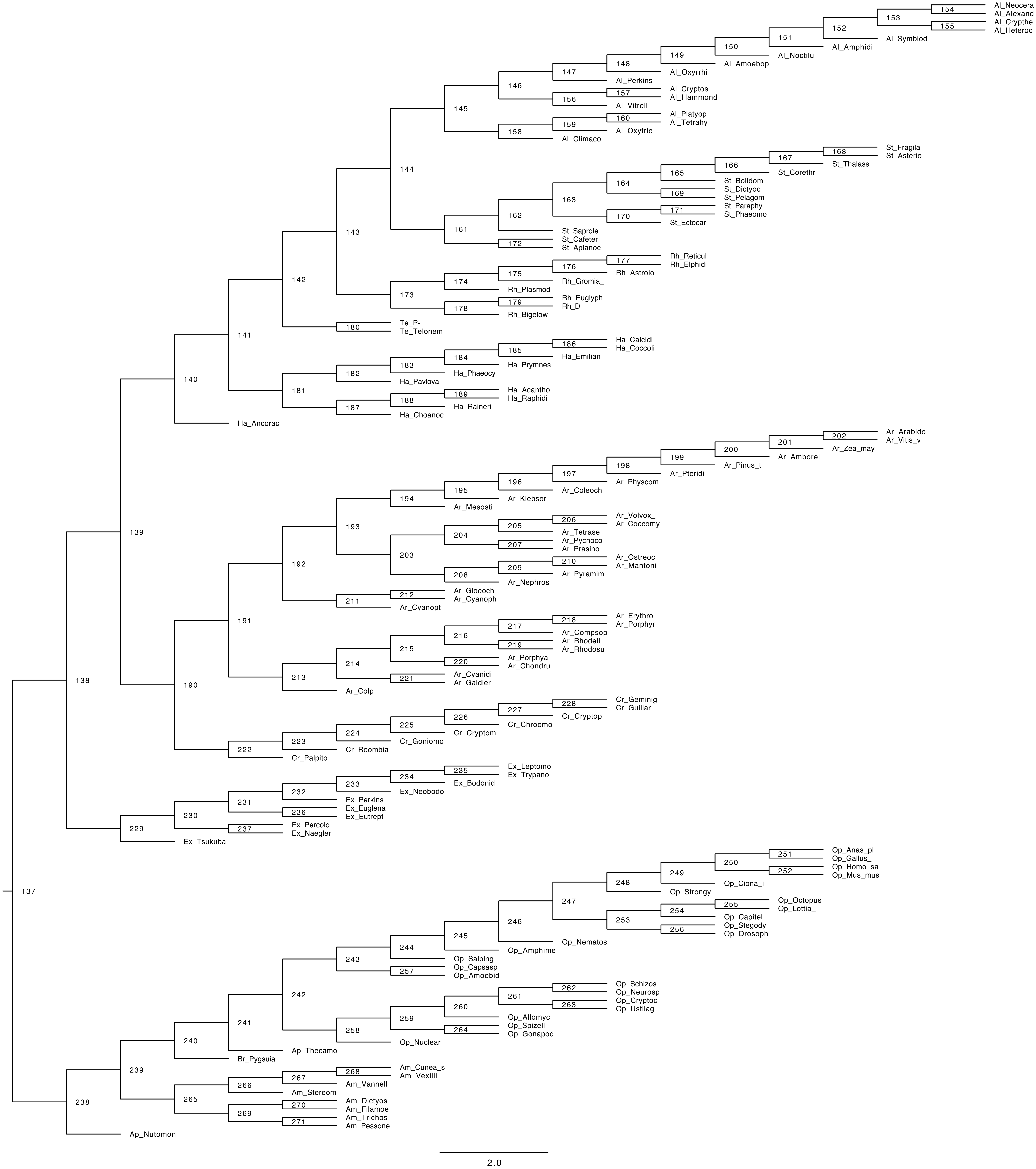
154
Al_Neocera
Al_Alexand
153
155
Al_Crypthe
Al_Heteroc
152
Al_Symbiod
151
Al_Amphidi
150
Al_Noctilu
149
Al_Amoebop
148
Al_Oxyrrhi
147
Al_Perkins
146
157
Al_Cryptos
Al_Hammond
156
Al_Vitrell
145
160
Al_Platyop
Al_Tetrahy
159
Al_Oxytric
158
Al_Climaco
168
St_Fragila
St_Asterio
167
St_Thalass
166
St_Corethr
165
St_Bolidom
164
169
St_Dictyoc
St_Pelagom
163
171
St_Paraphy
St_Phaeomo
170
St_Ectocar
162
144
St_Saprole
161
172
St_Cafeter
St_Aplanoc
143
177
Rh_Reticul
Rh_Elphidi
176
Rh_Astrolo
175
Rh_Gromia_
174
Rh_Plasmod
142
173
179
Rh_Euglyph
Rh_D
178
Rh_Bigelow
180
Te_P-
Te_Telonem
141
186
Ha_Calcidi
Ha_Coccoli
185
Ha_Emilian
184
Ha_Prymnes
183
Ha_Phaeocy
182
Ha_Pavlova
140
181
189
Ha_Acantho
Ha_Raphidi
188
Ha_Raineri
187
Ha_Choanoc
Ha_Ancorac
202
Ar_Arabido
Ar_Vitis_v
201
Ar_Zea_may
200
Ar_Amborel
199
Ar_Pinus_t
198
Ar_Pteridi
197
Ar_Physcom
196
Ar_Coleoch
195
Ar_Klebsor
194
Ar_Mesosti
193
206
Ar_Volvox_
Ar_Coccomy
205
Ar_Tetrase
204
207
Ar_Pycnoco
Ar_Prasino
139
203
210
Ar_Ostreoc
Ar_Mantoni
209
Ar_Pyramim
192
208
Ar_Nephros
212
Ar_Gloeoch
Ar_Cyanoph
211
Ar_Cyanopt
191
218
Ar_Erythro
Ar_Porphyr
217
Ar_Compsop
216
219
Ar_Rhodell
Ar_Rhodosu
215
220
Ar_Porphya
Ar_Chondru
214
221
Ar_Cyanidi
Ar_Galdier
138
213
Ar_Colp
190
228
Cr_Geminig
Cr_Guillar
227
Cr_Cryptop
226
Cr_Chroomo
225
Cr_Cryptom
224
Cr_Goniomo
223
Cr_Roombia
222
Cr_Palpito
235
Ex_Leptomo
Ex_Trypano
234
Ex_Bodonid
233
Ex_Neobodo
232
Ex_Perkins
231
236
Ex_Euglena
Ex_Eutrept
230
237
Ex_Percolo
Ex_Naegler
229
Ex_Tsukuba
251
Op_Anas_pl
Op_Gallus_
250
252
Op_Homo_sa
Op_Mus_mus
249
Op_Ciona_i
248
Op_Strongy
137
247
255
Op_Octopus
Op_Lottia_
254
Op_Capitel
253
256
Op_Stegody
Op_Drosoph
246
Op_Nematos
245
Op_Amphime
244
Op_Salping
243
257
Op_Capsasp
Op_Amoebid
262
Op_Schizos
Op_Neurosp
261
263
Op_Cryptoc
Op_Ustilag
242
260
Op_Allomyc
259
264
Op_Spizell
Op_Gonapod
241
258
Op_Nuclear
240
Ap_Thecamo
Br_Pygsuia
239
268
Am_Cunea_s
Am_Vexilli
267
Am_Vannell
266
Am_Stereom
265
270
Am_Dictyos
Am_Filamoe
269
271
Am_Trichos
Am_Pessone
238
Ap_Nutomon
2.0
