## Supplementary Data 2 for "A molecular timescale for the origin of red algal-derived plastids": README.pdf

**Single_genes.zip**, 320 ×
Gene.fa -> Raw sequences
Gene.filtered.ginsi.bmge.merged.fa -> PREQUAL-filtered, MAFFT G-INS-i-aligned, BMGE-trimmed, merged partial sequences belonging to same taxon

**Completeness.txt** -> Completeness of all taxa used for building the supermatrix, from which the initial ML tree (Fig. S1) has been inferred.
**136_OTUs.txt** -> Taxa and chimera that were used as OTUs.

**Supermatrix.fa** -> Concatenation of all 320 Gene.filtered.ginsi.bmge.merged.fa files.
**Supermatrix_320g136o.fa** -> Subsample of Supermatrix.fa; prior to concatenation, raw sequences were filtered, aligned, and trimmed as described in the manuscript. Note, this supermatrix contains also merged taxa (OTUs) for which the name structure does not follow the 8-column scheme that is described below.
**Supermatrix_320g63o.fa** -> Subsample of Supermatrix_320g136o.fa; real subsample, i.e., not newly aligned.

**Taxon naming (general notes):** Each taxon is named as follows: Group_Phylum_Class_Order_Family_Genus_SpeciesEpithet_Strain. I.e., underscores can be used as delimiter. The only exception from this is the strain "column" ($8^{th}$ column), where underscores can occur too. Strain names like js001, js002, js003 etc. were used for several taxa, for which strain information was not available or could not be reconstructed anymore. The $7^{th}$ column can contain further information that is added with a dash (e.g., -nucleomorph, -endo, -LGT). -endo stands for endosymbiont, -LGT stands for lateral gene transfer (excluded from the supermatrix!). To facilitate the detection of paralogs, wrong gene copy sequences of some taxa were added to several gene files (excluded from the supermatrix!). The information about these wrong copies is also added to column 7 (e.g., -psmb1 or simply -paralog). To several single genes, prokaryotes were added as well (see "Prokaryota" in the 1st column) to facilitated the detection of prokaryotic contaminants (excluded from the supermatrix!). Finally, highly underrepresented taxa (less than 5% data) were excluded from the supermatrix.
