## Supplementary Information (Figs. S1-S7, Table S1) for "A molecular timescale for the origin of red algal-derived plastids"

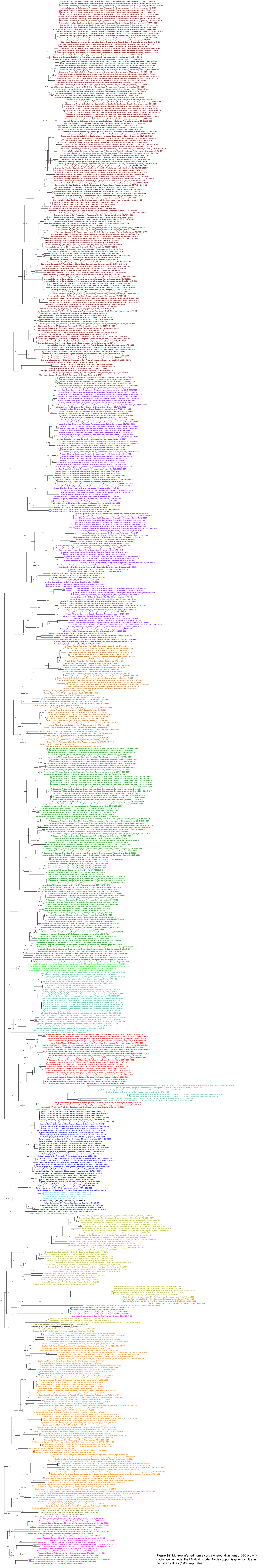

**Figure S1:** ML tree inferred from a concatenated alignment of 320 protein-coding genes under the LG+G+F model. Node support is given by ultrafast bootstrap values (1,000 replicates).

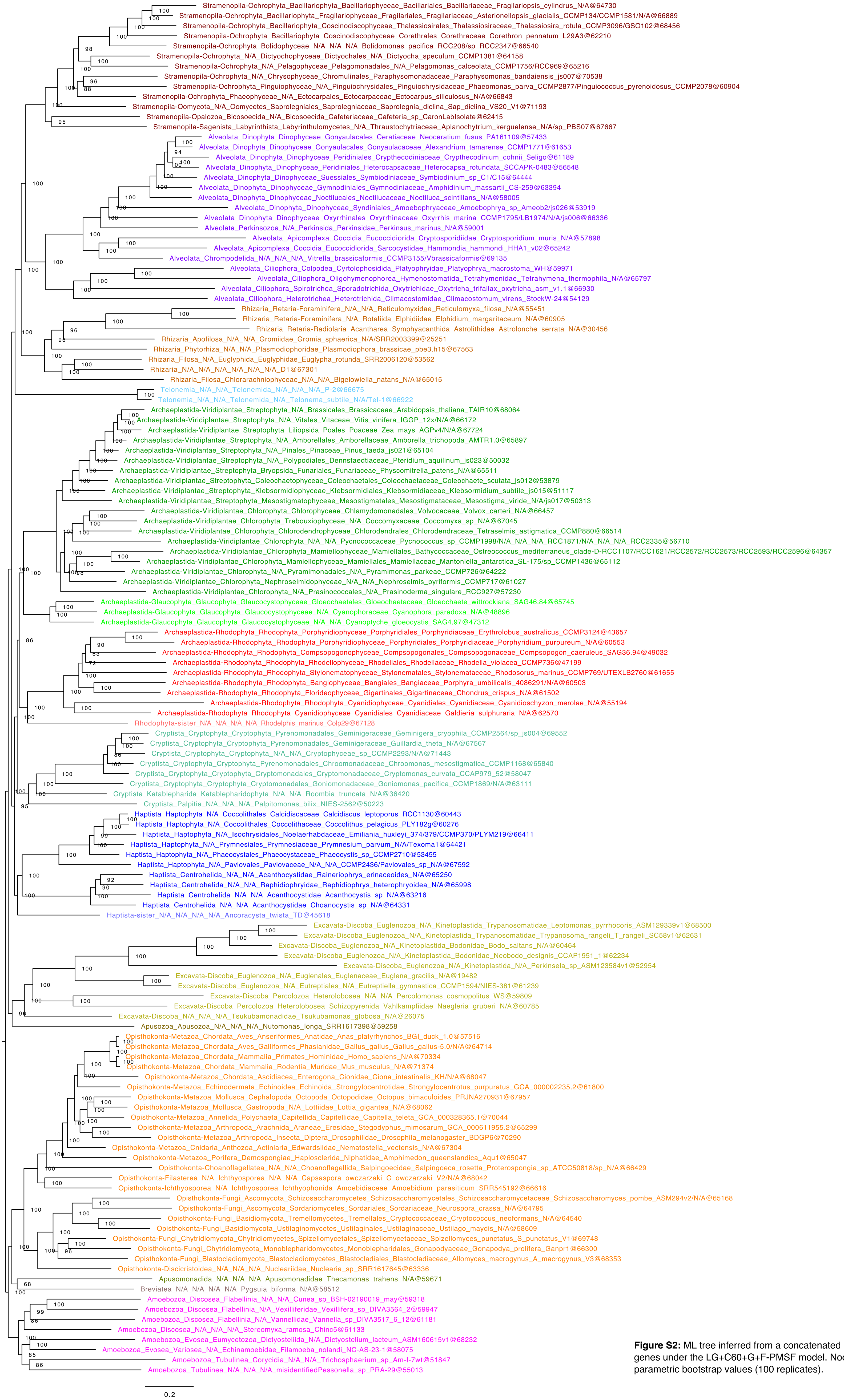


**Figure S2:** ML tree inferred from a concatenated alignment of 320 protein-coding genes under the LG+C60+G+F-PMSF model. Node support is given by non-parametric bootstrap values (100 replicates).

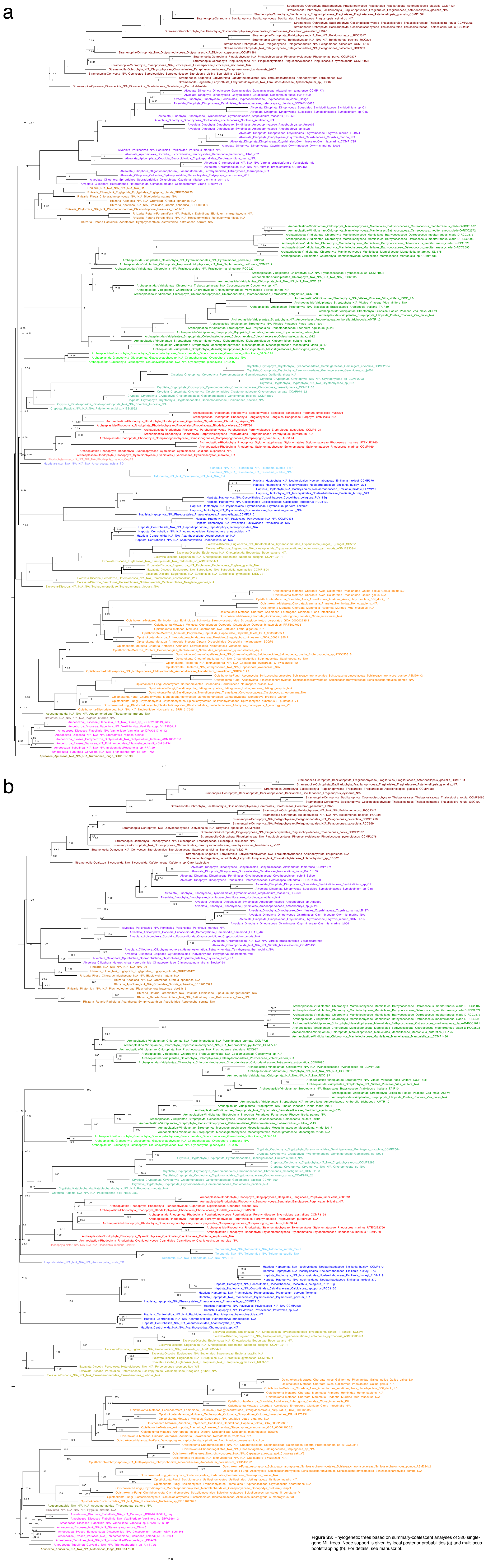

**Figure S3:** Phylogenetic trees based on summary-coalescent analyses of 320 single-gene ML trees. Node support is given by local posterior probabilities (a) and multilocus bootstrapping (b). For details, see manuscript.

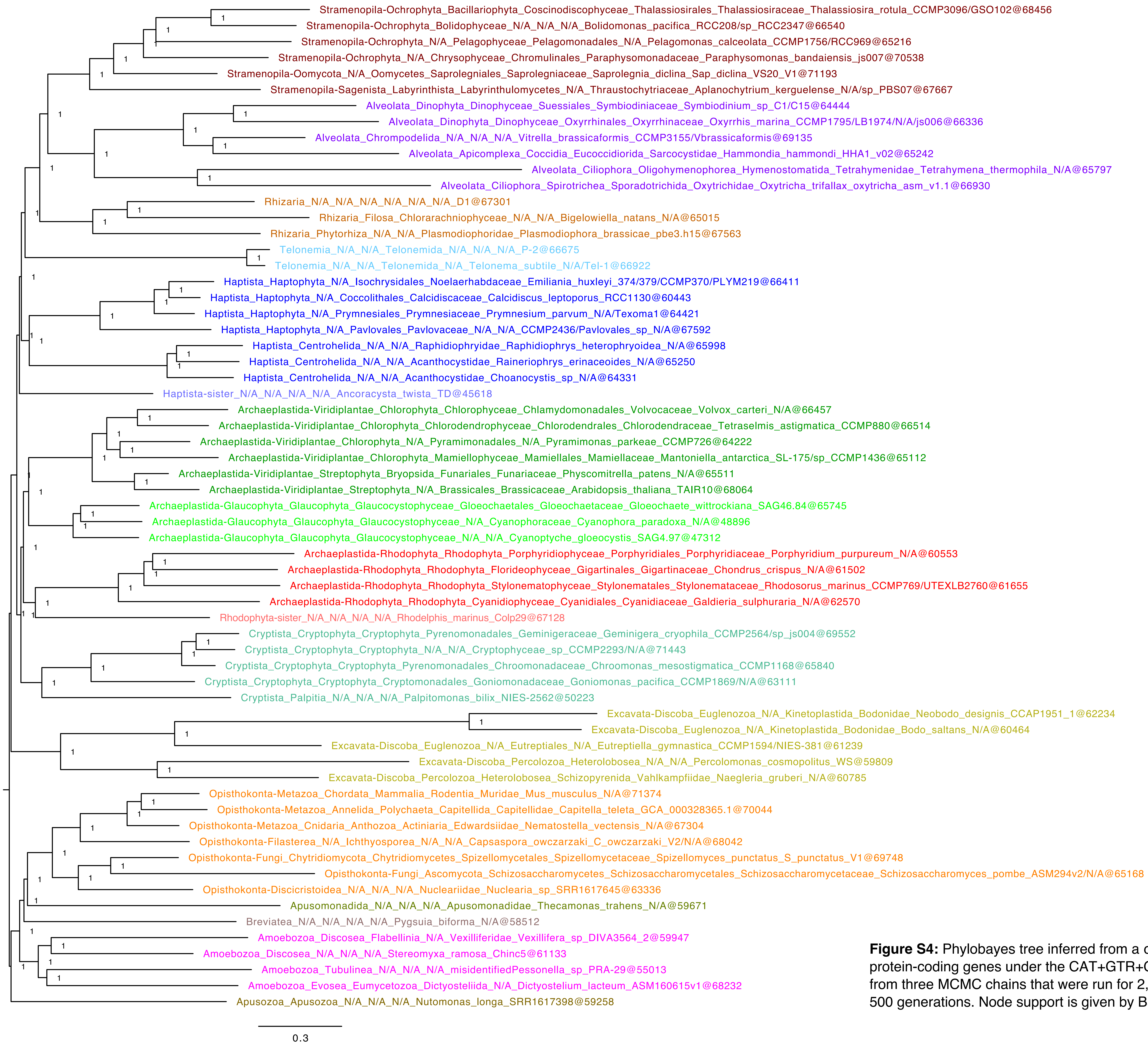


**Figure S4:** Phylobayes tree inferred from a concatenated alignment of 320 protein-coding genes under the CAT+GTR+G model. The tree is a consensus from three MCMC chains that were run for 2,598 generations with a burnin of 500 generations. Node support is given by Bayesian posterior probabilities.

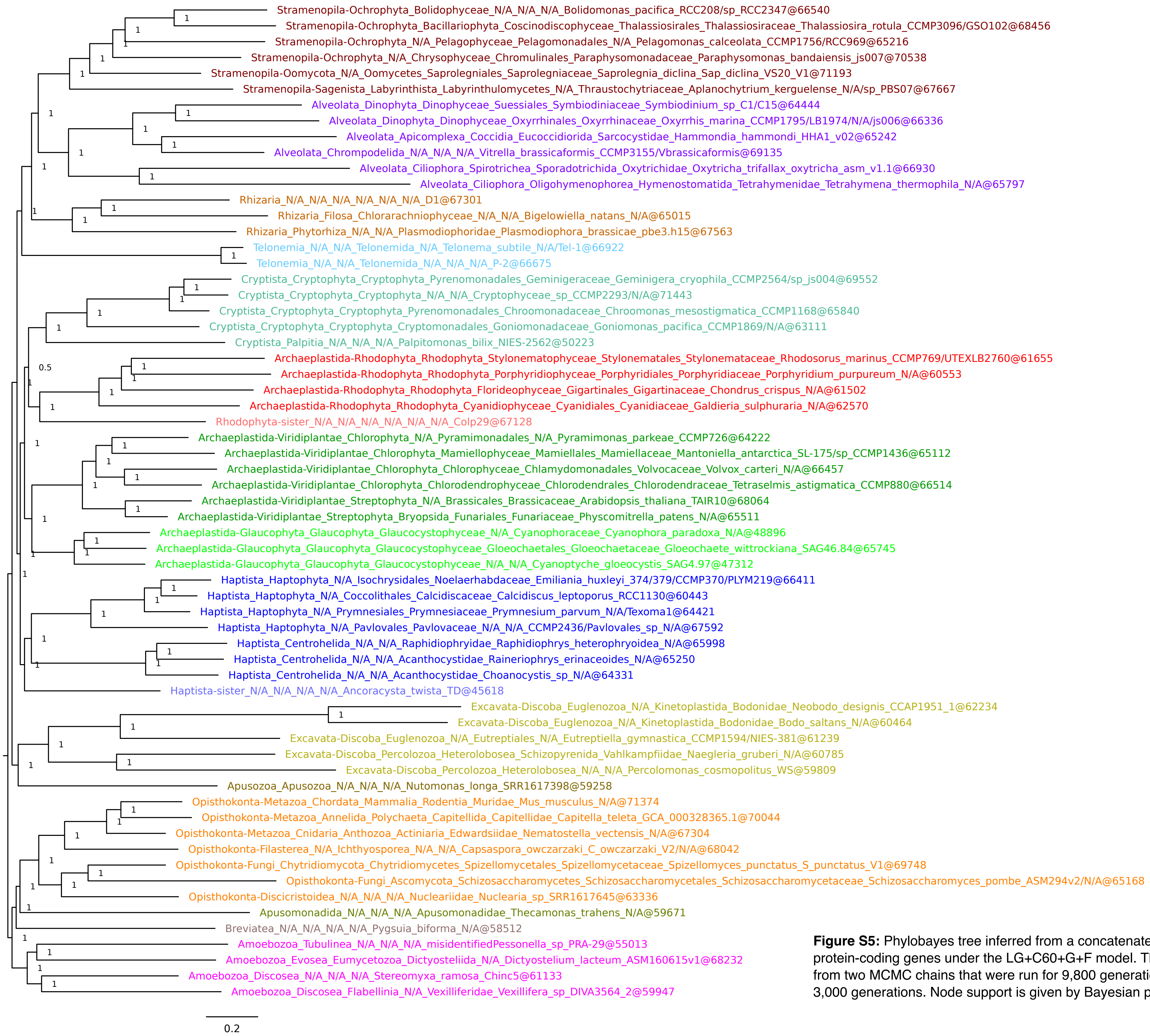


**Figure S5:** Phylobayes tree inferred from a concatenated alignment of 320 protein-coding genes under the LG+C60+G+F model. The tree is a consensus from two MCMC chains that were run for 9,800 generations with a burnin of 3,000 generations. Node support is given by Bayesian posterior probabilities.

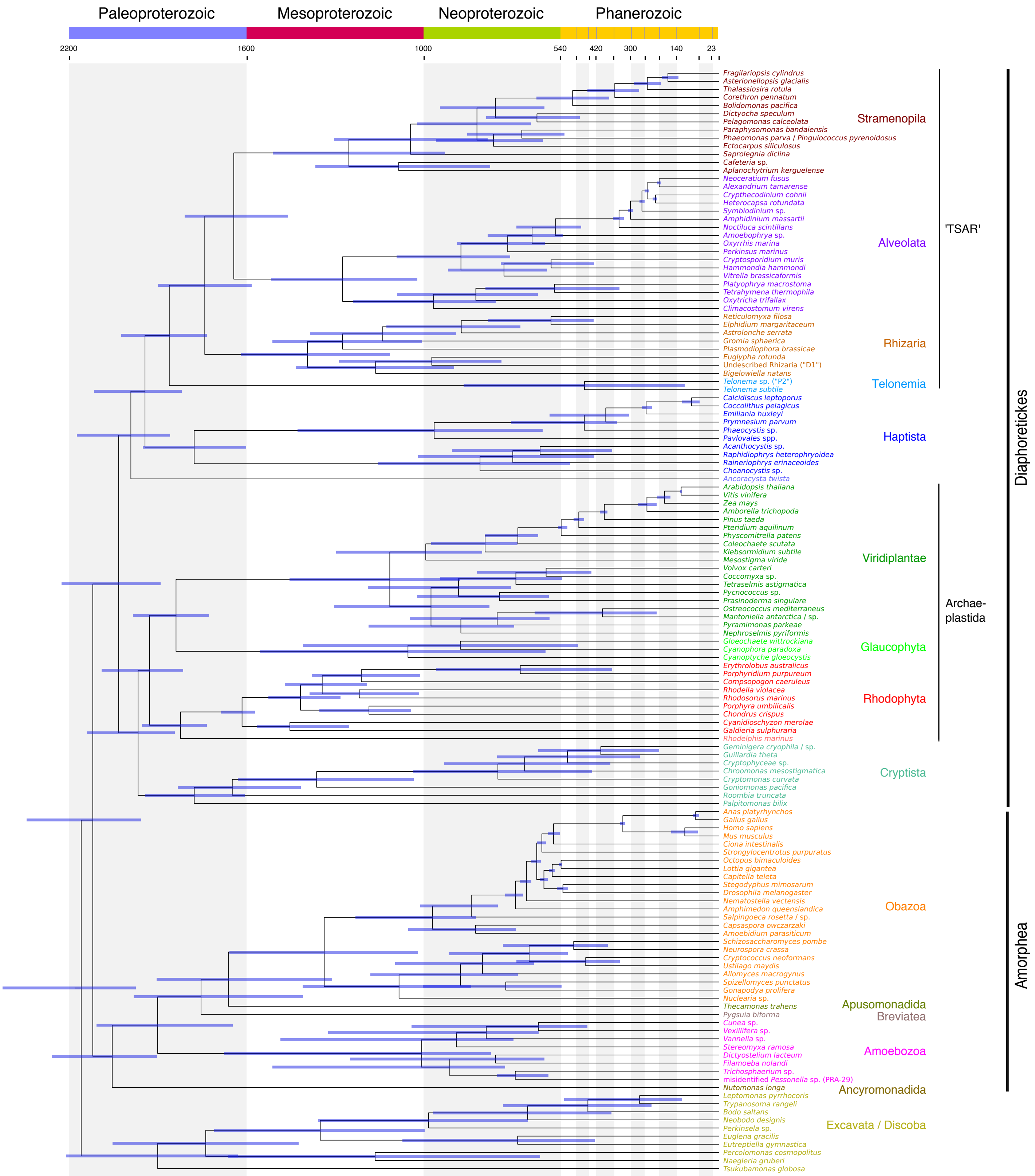


**Figure S6:** Time-calibrated phylogeny of extant eukaryotes. Divergence times were inferred with MCMCTree under an autocorrelated relaxed clock model and 33 fossil calibration points as soft-bound uniform priors (Table 1). The tree topology was reconstructed using IQ-TREE under the LG+C60+G+F model and a constrained tree search following the OTU-reduced CAT+GTR+G topology (Supplementary Fig. 4). Approximate likelihood calculations on the 320 gene concatenation under LG+G and a birth-death tree prior were used. Bars at nodes are 95% HPD. An absolute time scale in Ma and a geological time scale are shown. The tree depicted here was rooted on Excavata. An equivalent time-calibrated tree rooted on Amorphea is shown in Fig. 2.

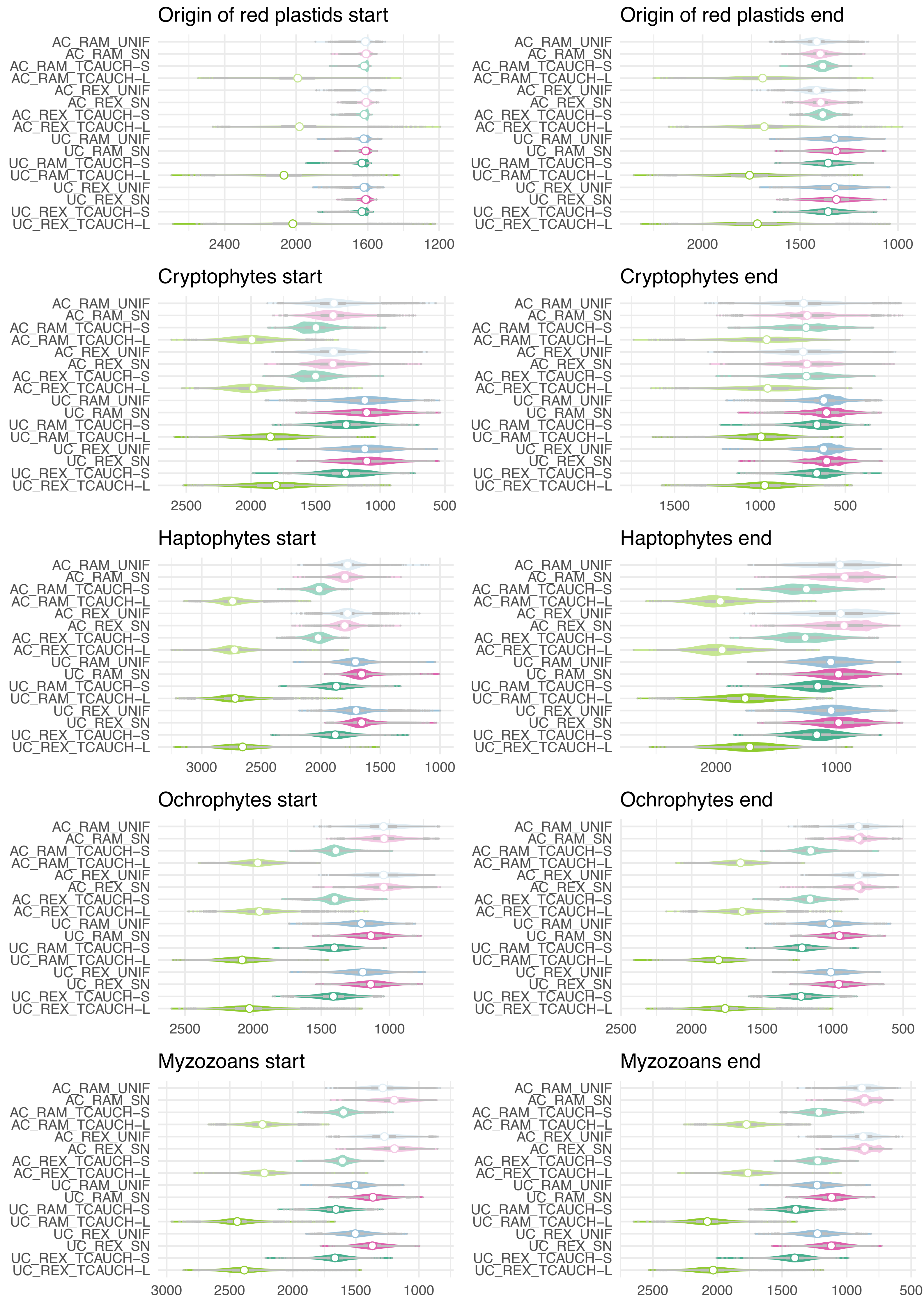


**Figure S7:** Divergence times for the ten relevant nodes that bracket the age of the five endosymbiosis events for secondary red algal-derived plastids under a broad range of conditions (MCMCTree analyses). Violin plots show the full posterior distribution. Mean ages are indicated with a white circle. AC – autocorrelated clock model, UC – uncorrelated clock model, RAM – root Amorphea, REX – root Excavata; calibration densities: UNIF – uniform, SN – skew-normal, TCAUCH–S/L – t-cauchy with short/long tails.

**Table S1:** Results of the posterior predictive analyses (PPA).

| | Site-specific AA heterogeneity* | | | Lineage-specific AA heterogeneity* | |
|---|---|---|---|---|---|
| **Model** | **div** | **conv** | **var** | **max** | **mean** |
| CAT+GTR+G | 9.85567 | 8.93904 | 9.60434 | 215.16 | 280.945 |
| LG+C60+G | 56.6831 | 34.5833 | 33.1079 | 256.79 | 318.385 |

*z-scores
